# Structured biological evidence enables general-purpose language models to predict bacteriophage–host interactions

**DOI:** 10.64898/2026.06.10.727770

**Authors:** Yao-zhong Zhang, Long Xu, Seiya Imoto

## Abstract

Biological interaction inference often requires integrating heterogeneous evidence that differs in biological meaning, provenance, specificity, and reliability. Existing computational approaches typically compress such evidence into numerical features, aggregate scores, or latent representations before prediction, obscuring the contribution of individual evidence records and limiting interpretation when evidence is conflicting or incomplete. Phage–host interaction prediction provides a demanding test case for this problem, requiring the integration of genomic, functional, and reference-derived evidence with differing coverage, specificity, and reliability. Here we introduce a structured-evidence inference paradigm that retains heterogeneous biological observations as modular, named, and experimentally perturbable records rather than compressing them before prediction. We implement this paradigm in PHI-Reason, which constructs structured textual profiles from genomic annotations, receptor-binding protein homology, nucleotide-neighbour relationships, alignment-free genomic similarity, and available CRISPR spacer information. A general-purpose large language model (LLM) performs inference directly over these structured profiles without task-specific training or manually engineered evidence-fusion rules.

Across phage–host benchmarks, PHI-Reason achieved species-level top-1 accuracies of 67.4% and 57.2% on RefSeq-634 and VHDB-3150, respectively, and a multi-host accuracy of 0.673 on the Hi-C cohort, outperforming established numerical methods. On 207 experimentally supported associations from 2025–2026 excluded from the reference corpus, it achieved 52.2% species-level and 90.8% genus-level top-1 accuracy. Evidence perturbations identified nucleotide-neighbour context as the dominant signal and quantified complementary or misleading evidence contributions. Target-conditioned local Jacobian and output-grounding analyses further characterized evidence-dependent inference and departures of generated rationales from the supplied evidence. Extending the framework to eukaryotic virus–host prediction preserved high predictive accuracy (67.0%), supporting cross-domain portability. Moreover, replacing the reasoning LLM with Jev, a typed-decision model, maintained broadly comparable performance across phage–host benchmarks while reducing mean per-query inference time by 11- to 40-fold, supporting the separation of evidence representation from downstream inference. These results establish structured biological evidence as a portable interface between heterogeneous biological observations and downstream inference. By separating evidence representation from the inference engine, this paradigm supports both language-model predictions with inspectable rationales and faster typed decisions for high-throughput prediction, while making the contribution and limits of evidential support directly testable.

## 1 Introduction

Many computational problems in biology are formulated as prediction tasks, yet their practical difficulty often lies in integrating evidence generated by disparate experiments, annotations and reference databases. Such evidence can differ in biological meaning, provenance, taxonomic specificity and reliability, and individual records may be missing, redundant or conflicting. Phage–host assignment provides a demanding test case for this problem. Bacteriophages shape bacterial population structure, horizontal gene transfer and microbial community function across natural, engineered and host-associated ecosystems [1], and identifying their bacterial hosts is critical for studying microbial ecology and developing phage-based interventions. However, host association is not determined by a single genomic feature. Instead, it must be inferred from complementary evidence–including evolutionary relatedness, adsorption-associated functions, bacterial defence records and similarity to previously characterized phages–each with uneven taxonomic coverage and variable reliability.

Current computational phage–host interaction (PHI) prediction methods include sequence-composition and similarity approaches [2–4], database-integrated and rule-based pipelines [5, 6], and machine-learning or representation-learning models [7–10]. These approaches enable large-scale host assignment and perform well when informative reference sequences or training examples are available. However, when multiple evidence sources are used, they are commonly integrated through numerical scores, fixed decision rules or learned latent representations. Although some pipelines retain individual evidence calls, their final predictions are derived from method-specific fusion procedures. Consequently, it is difficult to determine whether a prediction depends on a specific evidence–host relationship, redundant signals or properties of the representation itself. While conventional feature attribution and ablation can quantify dependence on model inputs, interventions on numerical variables do not necessarily preserve or isolate the semantics of named biological records. A complementary inference interface is therefore needed in which evidence remains individually identifiable and systematically perturbable while the prediction procedure is held fixed.

We propose treating structured biological evidence itself as the prediction interface. In this formulation, heterogeneous observations are retained as modular, named records rather than first being collapsed into aggregate scores or shared latent representations. This interface separates evidence representation from inference and can, in principle, be paired with different inference engines. A general-purpose large language model (LLM) provides one such inference layer because evidence with different biological semantics can be expressed within a shared textual schema without learning a new joint numerical representation for each combination of evidence types [11, 12]. However, the capacity of an LLM to generate fluent text does not guarantee reliable evidence integration. Pretrained models may rely on prior associations, overlook conflicting records or generate plausible claims that are unsupported by the supplied evidence. Our work demonstrates that a key challenge in scientific LLM use is disentangling profile grounding from predictive correctness: faithful rationales do not always yield correct predictions, and conversely, prediction errors often arise from incomplete evidence rather than unsupported model hallucinations.

Here we introduce PHI-Reason, an implementation of this structured-evidence inference paradigm for candidate-conditioned biological prediction using general-purpose LLMs. For each query, PHI-Reason constructs a structured profile containing modular fields–including genomic annotations, receptor-binding-protein homology, nucleotide-neighbour relationships, alignment-free genomic similarity, CRISPR-derived associations and candidate-host metadata. A frozen LLM integrates these records either to rank hosts within a species-level catalogue or to assess a nominated phage–host pair, without fitting model parameters or specifying numerical fusion weights. By preserving the identity and provenance of individual records, this separation of evidence construction from inference permits field-level interventions while keeping the model backbone fixed. Crucially, the modularity of this design ensures that the inference engine is decoupled from the biological domain, allowing the same contract to be applied to any interaction prediction task by simply swapping the domain-specific evidence fields.

To validate this paradigm, we evaluated PHI-Reason across distinct datasets, evidence regimes, biological interaction domains, and inference architectures. Across phage–host benchmarks, PHI-Reason achieved species-level top-1 accuracies of 67.4% and 57.2% on RefSeq-634 and VHDB-3150, respectively, and a multi-host accuracy of 0.673 on the Hi-C cohort, outperforming established numerical methods. A same-profile embedding-only control showed that, although the structured profiles contained substantial host-discriminative information, candidate-conditioned LLM inference extracted significantly greater predictive value. On an additional set of 207 experimentally supported phage–host associations reported in 2025–2026 and excluded from the reference corpus, PHI-Reason achieved species-level and genus-level top-1 accuracies of 52.2% and 90.8%, respectively. Systematic field removal and host-label perturbations on RefSeq-634 identified labelled nucleotide-neighbour context as the dominant evidence source and revealed complementary contributions from receptor-binding-protein and CRISPR evidence. Output-grounding analyses and a target-conditioned local Jacobian readout further characterized evidence-dependent inference at multiple levels, including when generated rationales departed from the supplied profiles.

Extending the framework to eukaryotic virus–host prediction with domain-appropriate evidence preserved high predictive accuracy (67.0%), supporting portability across biological interaction domains. Moreover, replacing the reasoning LLM with Jev, a typed-decision model, maintained broadly comparable predictive performance across the phage–host benchmarks while reducing mean per-query inference time by 11- to 40-fold, supporting the separation of evidence representation from downstream inference.

In summary, these results establish structured biological evidence as an explicit, perturbable, and portable interface for biological inference. By separating evidence construction from downstream inference, the framework accommodates new evidence modules without retraining and supports complementary inference engines for different operational settings, while making evidence contributions and the boundaries of reliable inference directly testable.

## 2 Results

### 2.1 Overview of the proposed framework

The proposed framework reformulates biological interaction prediction as candidate-conditioned inference over explicit, structured biological evidence profiles (Fig. 1). Rather than compressing heterogeneous biological observations into a single numerical feature vector, aggregate score or latent representation before prediction, PHI-Reason preserves distinct evidence sources as modular, named and independently perturbable records. This design separates biological evidence construction from downstream inference and treats the structured profile as a shared prediction interface that can be reused across different task formulations and inference architectures.

**Fig. 1.**
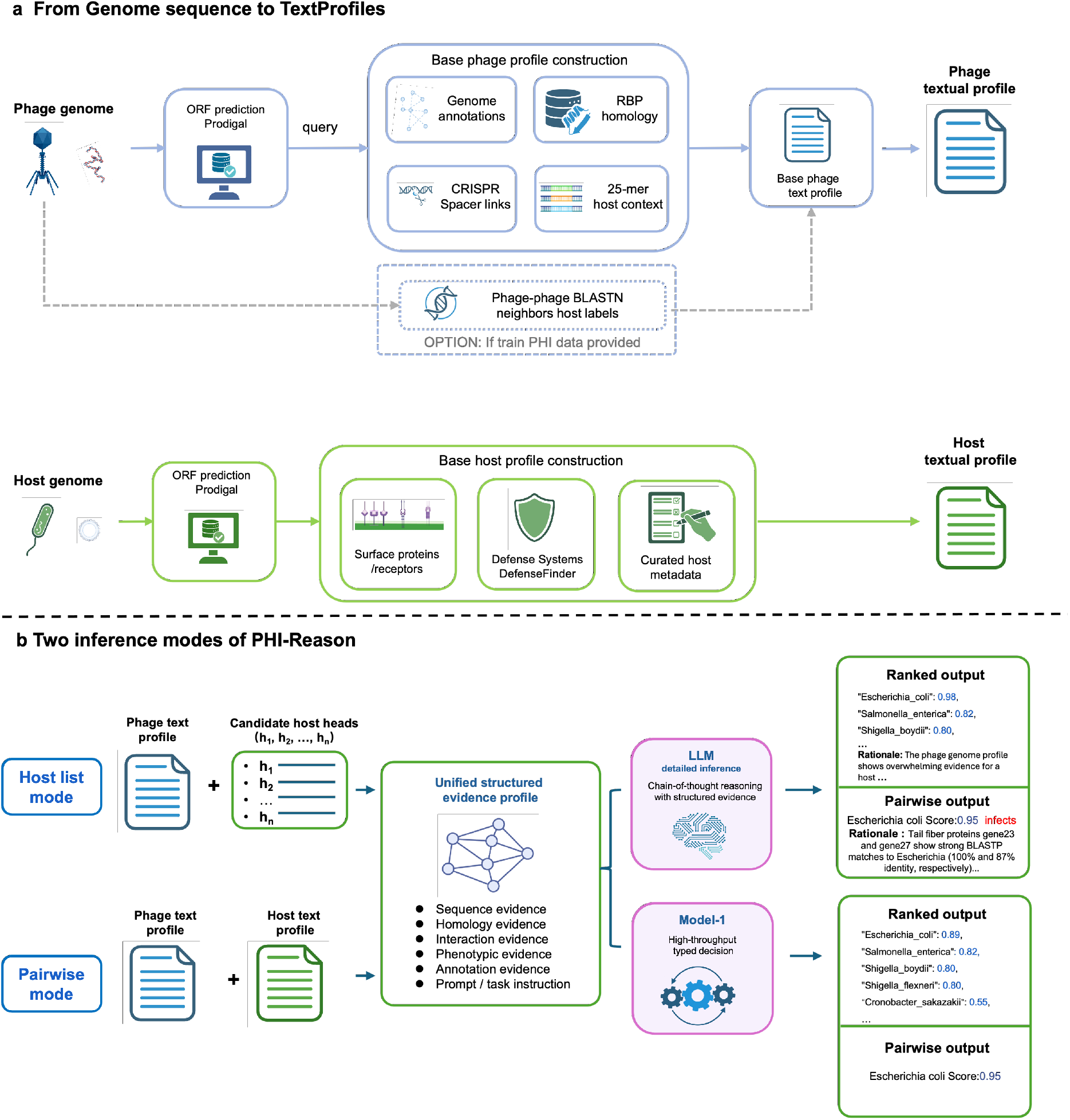
PHI-Reason converts genome-derived evidence into structured text profiles for host-list ranking and pairwise assessment. **a**, Deterministic pipelines convert phage and host genomes into named, perturbable evidence fields (annotations, receptor-binding-protein evidence and, when available, BLASTN reference-context blocks carrying host labels). **b**, The frozen language model runs in two modes: host-list ranking of all candidate hosts in one call, and pairwise scoring of a nominated phage–host pair. Created with BioRender.

For each query phage, PHI-Reason constructs a structured evidence profile containing base genome annotations, receptor-binding-protein (RBP) homology identified by BLASTP, labelled nucleotide-neighbour context derived from whole-genome BLASTN, alignment-free 25-mer genomic similarity and CRISPR-spacer associations when available. Candidate-host profiles summarize bacterial surface features, receptor-associated functions, defence repertoires and ecological or isolation-source information. Sequence-based searches further retrieve related reference phages, whose known host associations are incorporated into the profile as explicit contextual evidence.

Because these evidence records remain explicit, individual profile fields can be removed, isolated, relabelled or perturbed, creating a controlled evidence-perturbation platform. By holding the remaining evidence and task configuration fixed, the framework enables direct analysis of the contribution, complementarity, redundancy and potentially misleading effects of different evidence sources, while also providing a common basis for downstream evidence-grounding and error analyses.

PHI-Reason supports two complementary task modes over the same structured evidence interface (Fig. 1b). In the host-list mode, a query phage is evaluated against all species in a predefined candidate-host catalogue, producing an ordered host list for catalogue-wide prioritization. In the pairwise mode, a nominated phage–host pair is evaluated independently, yielding a focused compatibility score for that specific pair. The two modes therefore provide complementary views of the same prediction problem: global ranking over a candidate catalogue and focused assessment of predefined phage–host hypotheses.

The choice of task mode is independent of the choice of inference engine. In the primary analyses, a frozen general-purpose language model integrates the structured evidence and generates natural-language rationales, supporting detailed inference, evidence-grounding and error analysis. The same structured evidence interface can also be supplied to a typed-decision inference engine, which directly returns structured probabilistic judgments and thereby supports higher-throughput inference (Section 2.9). Thus, host-list and pairwise define how the prediction task is organized, whereas language-model and typed-decision inference define how the same structured evidence is converted into a decision.

We evaluated the framework along four complementary dimensions. Cross-benchmark comparisons, a same-profile embedding-only control, multi-backbone experiments, and post-cutoff evaluations on 207 newly reported experimental phage– host associations and 14 recently deposited phage genomes assessed predictive performance and how effectively different inference engines exploited the information contained in the structured profiles. Evidence-field removal, reference-label perturbation and identifier masking examined dependence on individual evidence sources and evidence–label correspondence, including evidence dependence in the post-cutoff setting. Output-grounding and intermediate-state analyses characterized how model outputs related to the supplied evidence. Finally, portability was evaluated along two axes: across biological domains by replacing domain-specific evidence modules, and across inference architectures by retaining the structured evidence while replacing the downstream inference engine.

### 2.2 A structured-profile perturbation platform dissects evidence dependence in host prediction

PHI-Reason represents each phage as a set of named evidence fields, including BLASTN-derived nucleotide-neighbour context, receptor-binding-protein (RBP) matches, alignment-free genomic similarity and host CRISPR-spacer records, which a frozen general-purpose language model integrates while jointly ranking a fixed catalogue of candidate host species, without fitting model parameters to phage-host labels (Fig. 2a). We used Qwen3.6:27b in its no-thinking configuration, an open-weight dense language model. Because the evidence fields and their associated host labels remain explicit, individual fields can be added, withheld, stripped of host labels or label-scrambled while the model, prompt, decoding settings and candidate catalogue remain fixed. This converts host prediction into a controlled evidence-intervention platform in which the contribution, redundancy and failure modes of individual evidence sources can be examined independently of model retraining. We applied this platform to RefSeq-634, the held-out partition of the CHERRY species-level phage–host collection, in which 634 evaluation phages were ranked against 223 candidate host species. A separate partition of 1,306 phages was used only as the reference collection for retrieving labelled nucleotide neighbours (Methods).

**Fig. 2.**
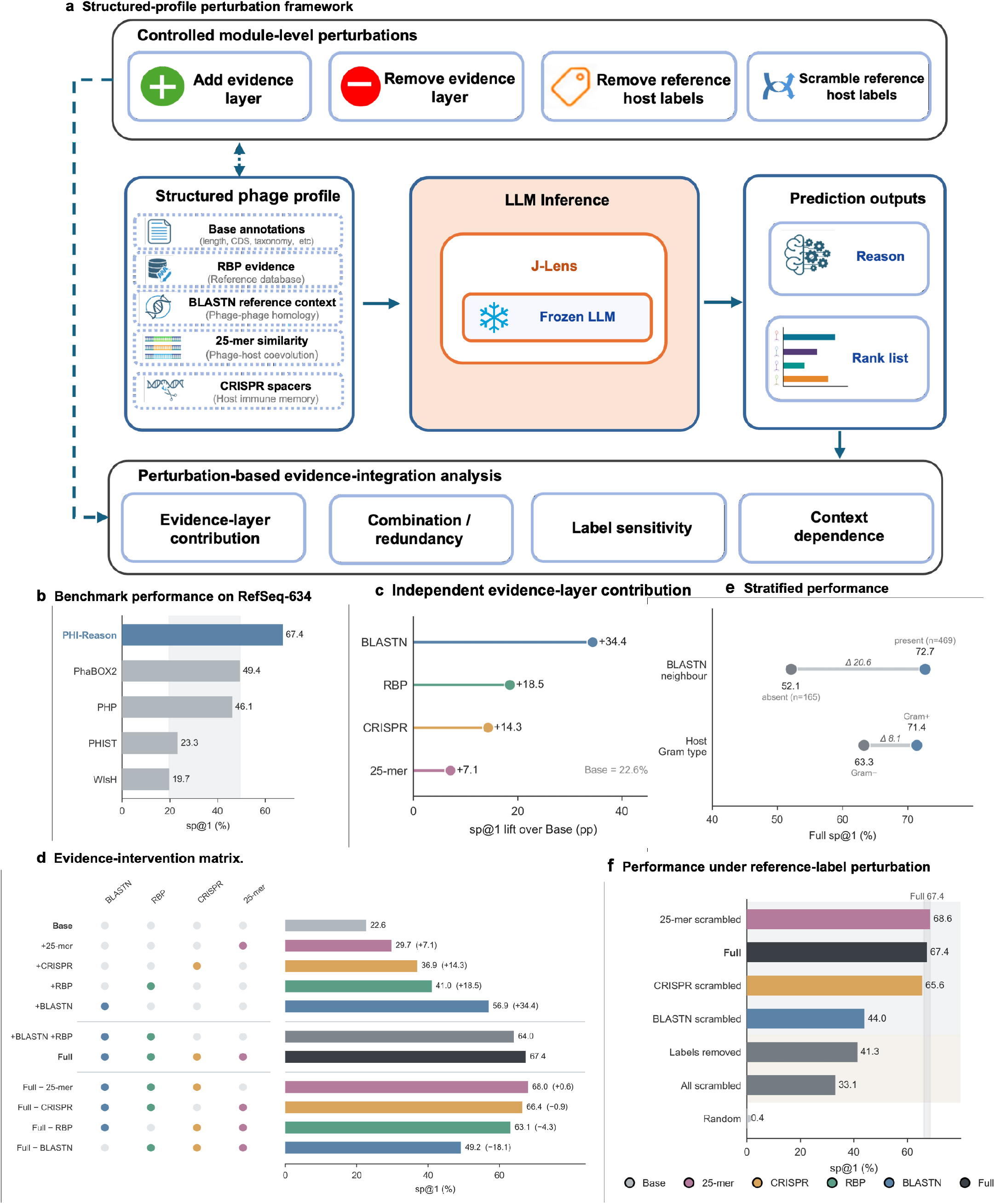
Structured-profile perturbations reveal overlapping and label-dependent evidence integration. **a**, Perturbation framework. Four labelled evidence blocks are added to an annotation-only base and passed to the frozen backbone, which returns a rationale and a ranked host list; the four module-level perturbations shown drive the analyses in panels **c**–**f**. Model, prompt, decoding settings and candidate catalogue are identical across all conditions. **b**, PHI-Reason reached sp@1 67.4%, above PhaBOX2 (49.4%), PHP (46.1%), PHIST (23.3%) and WIsH (19.7%). **c**, Single-field lift over Base (22.6%): BLASTN +34.4, RBP +18.5, CRISPR +14.3, 25-mer +7.1 pp. **d**, Contribution matrix: Base + BLASTN + RBP already reached 64.0% (near Full 67.4%); removing BLASTN caused the largest drop (*−*18.1 pp). **e**, Stratification: sp@1 72.7% with a labelled BLASTN neighbour (*n* = 469) versus 52.1% without (*n* = 165); Gram-positive 71.4% versus Gram-negative 63.3%. **f**, Label perturbation: scrambling all three labelled reference blocks lowered sp@1 to 33.1%, well above the *≈*0.4% random floor.

The full profile achieved species-level top-1 accuracy (sp@1) of 67.4% (95% confidence interval (CI), 63.6–70.9%; Wilson score interval; Fig. 2b), 18.0 percentage points (pp) above the strongest baseline evaluated on this benchmark (PhaBOX2, 49.4%). We used this full-profile configuration (Methods; configB) as the reference condition for the perturbation analyses below, including as the endpoint of the field-intervention ladder. A minimal Base profile containing candidate-host names and Prodigal-, eggNOG- and PHROG-derived genome annotations achieved sp@1 = 22.6%, whereas the full profile reached 67.4%, a gain of 44.8 pp. Adding one evidence field at a time to the Base profile revealed substantial differences in predictive contribution (Fig. 2c). Labelled BLASTN nucleotide-neighbour context produced the largest gain, increasing sp@1 to 56.9% (+34.4 pp). RBP, CRISPR and 25-mer evidence produced smaller gains, reaching 41.0% (+18.5 pp), 36.9% (+14.3 pp) and 29.7% (+7.1 pp), respectively. This ordering probably reflects differences in evidence origin, reference coverage, taxonomic representation and specificity rather than a universal hierarchy of biological importance. For example, 25-mer evidence was available for 534 of 634 phages but had limited host specificity, whereas CRISPR evidence was present for only 29% of queries but was frequently concordant with the true host genus when available (Supplementary Table 3).

Field-combination and leave-one-field interventions distinguished complementary from overlapping information (Fig. 2d). The Base+BLASTN+RBP profile achieved sp@1 = 64.0%, close to the 67.4% obtained with the full profile, indicating that these two fields captured most of the predictive information in the complete profile. Consistent with this result, removing BLASTN reduced accuracy to 49.2% (−18.2 pp), the largest leave-one-field effect. Removing RBP or CRISPR produced smaller reductions, to 63.1% (−4.3 pp) and 66.4% (−1.0 pp), respectively, whereas removing 25-mer evidence left accuracy essentially unchanged (68.0%, +0.6 pp, within the run-to-run sampling noise described above). Thus, BLASTN provided the dominant and least replaceable reference-conditioned signal, RBP and CRISPR supplied smaller complementary contributions, and 25-mer similarity added little once more specific evidence was available. Because the RBP field was derived from matches to an external structure-predicted reference corpus, its contribution should be interpreted as corpus-mediated evidence rather than as an independent de novo species-level predictor (Methods).

Evidence utility varied across queries (Fig. 2e). The 469 phages with at least one labelled BLASTN neighbour achieved sp@1 = 72.7% (95% CI, 68.5–76.5%), compared with 52.1% (95% CI, 44.5–59.6%) for the 165 phages without such a neighbour. The difference was 20.6 pp (95% CI, 12.0–29.1 pp; Newcombe method). This comparison is observational because phages lacking labelled neighbours may also be more evolutionarily novel or otherwise more difficult to assign. Nevertheless, the full profile retained appreciable performance in this subset. Among neighbour-absent phages, 25-mer and RBP evidence remained available for 88.5% and 72.1% of queries, respectively, although their separate contributions cannot be inferred from this stratified comparison. Full profile accuracy also differed by host Gram type, reaching 71.4% for Gram-positive hosts and 63.3% for Gram-negative hosts (Supplementary Note S1).

We next used label removal and scrambling to test whether prediction depended on correct evidence–host correspondence rather than merely on field presence or formatting (Fig. 2f). These interventions targeted the BLASTN, 25-mer and CRISPR fields, which contained explicit host labels, while preserving the RBP field and the remaining content and structure of the perturbed records. Removing host labels from all three fields reduced sp@1 to 41.3% (−26.1 pp), closely matching the Base+RBP condition (41.0%). Scrambling BLASTN labels produced the largest single-field reduction, lowering sp@1 to 44.0%, whereas scrambling CRISPR labels produced a smaller reduction to 65.6% and scrambling 25-mer labels did not reduce observed accuracy (68.6%). Simultaneously scrambling all three labelled fields reduced sp@1 to 33.1% (−34.3 pp). This remained above the approximately 0.4% random top-1 rate expected for 223 candidates, consistent with residual information in the Base annotations, candidate catalogue and unmodified RBP field. Relative to its own clean full-profile accuracy, BLASTN scrambling reduced performance by 23.4 percentage points, to 34.7% of the clean value, indicating that Qwen3.6:27b relies substantially on correctly labelled BLASTN evidence and is sensitive to this specific form of evidence corruption.

These interventions show that the predictive value of structured evidence was highly uneven and depended on both evidence availability and correct evidence–host correspondence. Labelled BLASTN neighbour context provided the dominant and least replaceable signal, RBP and CRISPR supplied smaller complementary contributions, and 25-mer similarity was broadly available but largely redundant in the full profile. Label removal and scrambling further showed that incorrect evidence–host associations actively redirected prediction. The structured-profile platform therefore resolves heterogeneous evidence into dominant, complementary, redundant, coverage-limited and misleading components rather than reducing their combined effect to a single aggregate score.

### 2.3 Evidence grounding distinguishes unsupported outputs from evidence-limited prediction errors

A central risk in applying large language models to scientific inference is that a generated rationale can appear biologically plausible while extending beyond the evidence supplied to the model. Because PHI-Reason uses a fixed evidence interface, both its rationale and its ranked host prediction can be evaluated directly against the corresponding structured profile. We defined an *evidence-interface hallucination* as an output that lacked support in the supplied profile, independently of whether the resulting prediction was biologically correct. We evaluated this departure at three levels (Fig. 3): claim-level grounding assessed whether individual rationale claims were supported by the profile; rationale-level grounding assessed whether a rationale contained one or more unsupported claims; and answer-level grounding assessed whether the selected host genus appeared in any host-supporting profile field. Predictive correctness was evaluated separately, allowing grounded errors to be distinguished from unsupported outputs and allowing unsupported content accompanying a correct prediction to remain detectable. These definitions are relative to the supplied profile rather than to the broader biological literature, and rationale grounding neither reveals the model’s internal inference process nor establishes causal evidence use.

**Fig. 3.**
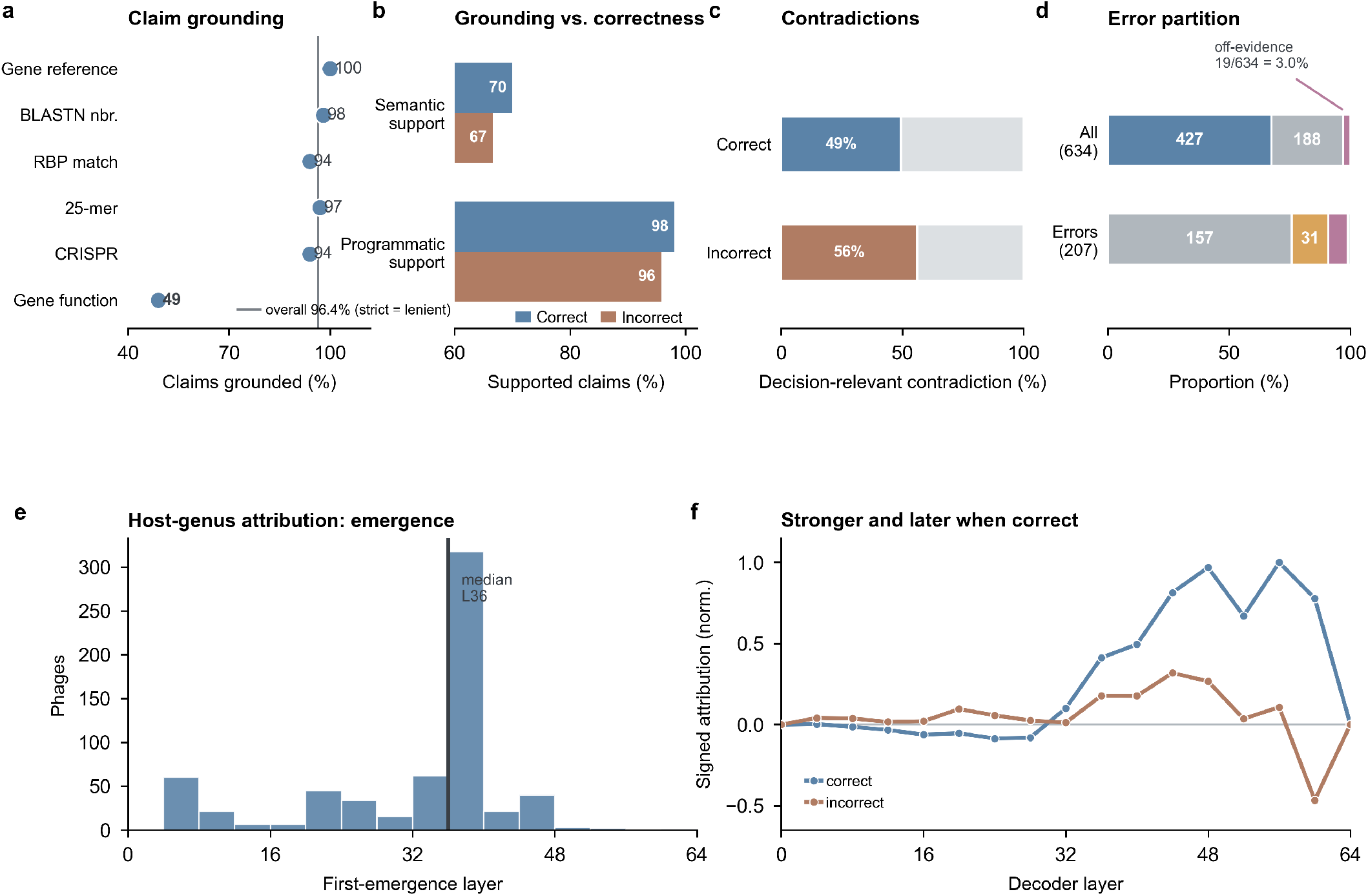
Evidence grounding separates unsupported claims from evidence-limited errors. **a**, Claim grounding: 96.4% overall, identical under the strict criterion; gene-function attribution claims show the lowest support (49%). **b**, Programmatic grounding does not track correctness (*<*3 pp difference between correct and incorrect predictions). **c**, Decision-relevant contradictions are weakly enriched among errors (56.0% versus 49.2%). **d**, Error partition: off-evidence calls are rare (19/634, 3.0%); most errors are non-decisive (157) or evidence-incomplete (48). **e**–**f**, Target-conditioned local Jacobian readout (matched bf16 model, non-causal): a positive first-order contribution to the confirmed host-genus logit emerges by mid-to-late network depth (median layer 36 of 64), later for correct than incorrect predictions (Supplementary Note S11).

Most extracted claims were supported by the supplied profiles, although support varied across evidence types. A programmatic verifier extracted 3,365 field-referenced claims from 634 rationales (5.3 claims per phage) and identified 96.4% micro-averaged support, with the stricter field-specific criterion yielding an identical value (Supplementary Note S8.2). These aggregate values were dominated by frequent and nearly completely grounded gene-reference and RBP-match claims. Support for sparser but decision-relevant host-linking claims reached 98% for BLASTN-neighbour hosts, 94% for CRISPR-associated hosts and 97% for 25-mer-associated hosts (Supplementary Table 16). A complementary blinded semantic assessment using Claude Opus 4.8 [13], applied with a fixed judging rubric, evaluated interpretive and unverifiable content in addition to field-referenced claims and classified 51.4% of rationales as containing at least one decision-relevant contradiction (Supplementary Note S8.3–S8.4), despite the higher programmatic faithfulness reported above; targeted re-auditing identified unsupported specific, checkable details, including invented gene counts, evidence-schema fields, neighbour-phage identifiers and candidate-host genera, that were not captured by the programmatic checks (Supplementary Note S8.4). Under the programmatic assessment, support differed by less than three percentage points between correct and incorrect predictions (98.1% versus 95.8%; Fig. 3b); under the semantic assessment the decision-relevant-contradiction rate differed by seven percentage points (56.0% incorrect versus 49.2% correct). Gene references were fully traceable (100%), whereas gene-function attributions showed the lowest support (49%), primarily because some rationales assigned molecular functions beyond those contained in the supplied annotations (Fig. 3a; Supplementary Note S8.2). Claim grounding therefore measured adherence to the evidence interface but only weakly discriminated prediction correctness.

Unsupported rationale content was associated with, but neither necessary nor sufficient for, prediction error. A decision-relevant contradiction concerning the selected host or the stated basis of the ranking occurred in 56.0% of incorrect predictions and 49.2% of correct predictions (Fig. 3c). Nevertheless, 91 of the 207 incorrect predictions (44%) contained no detected decision-relevant contradiction under this semantic assessment. Under the programmatic assessment, rationale-level grounding failures occurred in 46 of 207 incorrect predictions (22%) and in 36 of 427 correct predictions (8%). The semantic assessment produced higher rates because it additionally classified content that could not be verified from the profile (Supplementary Note S8.3). Conversely, an incorrect prediction could be accompanied by a fully grounded rationale when the available evidence was incomplete, ambiguous or misleading. Rationale grounding and predictive correctness therefore captured related but distinct properties of the model output.

To complement these input- and output-level analyses, we examined intermediate model states using a *target-conditioned local Jacobian readout* adapted from the Jacobian-lens approach [14] (Methods; Supplementary Note S11). For each phage, we supplied the frozen model with the structured profile and answer prefix and, at the position immediately preceding host-name generation, calculated the local first-order contribution of each decoder-block activation to the confirmed host-genus logit. This per-input, target-specific readout differs from the original context-averaged lens and produces neither a vocabulary ranking nor a reconstruction of the model’s reasoning. Across the 64 text-decoder blocks of Qwen3.6:27b, three-quarters of which use linear rather than full attention, the confirmed host-genus logit first acquired an appreciable positive contribution at a median layer of 36, corresponding to 56% of network depth (Fig. 3e). The threshold-crossing layer was stable across relative thresholds ranging from 10% to 50% of each phage’s positive peak (medians 32/36/36/44; Supplementary Note S11), and crossed markedly later among correct predictions (median layer 36) than incorrect predictions (median layer 20).

Correct and incorrect predictions also differed in the magnitude of the target-conditioned readout. Restricting the analysis to layers 32–53 of the 64-block Qwen3.6:27b model, correct predictions showed a substantially stronger mean positive attribution than incorrect predictions (0.88 versus 0.36; Supplementary Note S11; Fig. 3f). This comparison is not a matched contrast between correct and incorrect predictions at the per-phage level and should not be interpreted as evidence of a causal mechanism; the Jacobian-lens readout remains a non-causal, target-conditioned attribution (Methods).

At the answer level, assignments unsupported by all host-linking fields were uncommon. We defined an inference-time *off-evidence* flag when the selected host genus appeared nowhere in the host-supporting profile fields. This flag depended only on the profile and predicted host and was therefore independent of ground-truth correctness. It was raised for 19 of 634 predictions (3.0%; 95% CI, 1.9–4.6%; Wilson score interval), all of which were incorrect. These cases accounted for 9.2% of the 207 prediction errors and none of the 427 correct predictions. In 17 of the 19 cases, the confirmed genus was also absent from the profile; in the remaining two, the confirmed genus was represented but the model selected a genus unsupported by any host-linking field.

We partitioned the 207 errors into three mutually exclusive categories to distinguish answer-level departure from limitations of the supplied evidence (Fig. 3d; Supplementary Note S8.7). A total of 157 errors were *non-decisive*: both the confirmed and predicted genera were supported, but an evidence-supported competitor was ranked first. A further 48 were *evidence-incomplete*, with the confirmed genus absent from the profile; these comprised 17 off-evidence assignments and 31 errors in which the selected genus remained anchored to supplied evidence. The remaining two were *recoverable-profile* errors in which the confirmed genus was supported but the model selected an unsupported genus. Thus, most incorrect predictions arose when the profile omitted the confirmed host or supported multiple competing genera, whereas only 19 errors involved selection of a genus detached from all supplied host-linking evidence.

The top-1–top-2 score margin provided a complementary inference-time signal for prediction correctness, reaching an area under the receiver operating characteristic curve (AUROC) of 0.854 on RefSeq-634 (95% CI, 0.824–0.883) and 0.823 on the class-balanced Balanced-200 subset. Off-evidence errors had a smaller mean margin than correct predictions (0.030 versus 0.146; Supplementary Fig. 1). The margin was evaluated as a general correctness signal, however, and its association with off-evidence errors does not make it a specific detector of evidence departure. Resampling self-consistency was a relatively weak correctness indicator (AUROC = 0.672) and remained lower among answer-level-hallucination cases (mean genus self-consistency, 0.59) than among correct predictions (0.98). High self-consistency therefore did not certify that a prediction was supported by the available evidence.

In all, these analyses separate claim grounding, rationale-level unsupported content, answer-level evidence departure and predictive correctness. Most errors reflected incomplete or non-decisive evidence rather than selection of a host genus unsupported by the profile, and grounded rationales could accompany both correct and incorrect predictions. Evidence-interface hallucination is therefore a specific, operationally measurable departure from the supplied evidence boundary, rather than a synonym for ordinary prediction error, low confidence or variability across repeated generations.

### 2.4 Structured profiles support evidence-grounded species-level PHI prediction without a task-specific classifier

Having shown on RefSeq-634 that structured profiles expose how distinct evidence fields contribute to host ranking, we next asked whether the same representation supports species-level PHI prediction across broader evidence regimes. We evaluated PHI-Reason on VHDB-3150, derived from the VHDB collection [15], using 1,280 CHERRY-overlapping phages as the reference and training partition and 3,150 phages for evaluation. We also evaluated two Hi-C-derived metagenomic cohorts obtained from the proximity-ligation data of Bignaud et al. [16]. The Hi-C data were restricted to species-level host labels and comprised 52 host species and 406 phages, from which two evaluation subsets were considered: a multi-host-only subset, Sp-1 (*n* = 46), and a host-number-stratified subset, Sp-2 (*n* = 82). Across benchmarks, the profile schema, inference prompt and frozen model configuration were held fixed; unavailable evidence fields remained empty, and only the retrieved evidence and candidate-host catalogue changed with the dataset.

We compared PHI-Reason with sequence-, composition-, supervised- and database-based methods—PHIST [2], WIsH [3], PHP [4], DeepHost [10] and PhaBOX2 [6]— retraining trainable methods where applicable. PhaBOX2 was run in MAG mode against the candidate-host genomes while retaining access to its complete internal database and was evaluated on the isolate-derived benchmarks. DeepHost was evaluated only on the Hi-C cohorts, for which a leakage-controlled retraining configuration could be established. iPHoP [5], which calibrates host assignments at the genus rather than species level, was compared separately at its native taxonomic resolution (Supplementary Table 26). We report species-level top-1 accuracy (sp@1) for the isolate-derived benchmarks and species-level multi-host accuracy (MHA-sp) for the Hi-C cohorts, always using the complete dataset denominator, with abstentions, unresolved outputs and predictions outside the candidate-host catalogue counted as errors. Because the evaluated methods differed in reference access, training labels and prediction coverage, these comparisons provide a practical benchmark-specific performance reference rather than a controlled comparison of inference model classes.

On RefSeq-634, PHI-Reason achieved sp@1 = 67.4%, compared with 49.4% for PhaBOX2, 46.1% for PHP, 23.3% for PHIST and 19.7% for WIsH. On VHDB-3150, PHI-Reason achieved sp@1 = 57.2%, compared with 43.8% for PhaBOX2, 32.8% for PHP, 16.5% for PHIST and 13.5% for WIsH (Fig. 4a; Supplementary Table 1). Under these evidence regimes, PHI-Reason achieved the highest observed species-level accuracy among the evaluated methods on both isolate benchmarks, while using a frozen language model rather than a task-specific classifier. Prediction coverage differed across tools, but uncovered cases were retained in the denominator and counted as errors, so the reported values reflect complete-set performance rather than accuracy conditional on successful prediction (Supplementary Table 1).

**Fig. 4.**
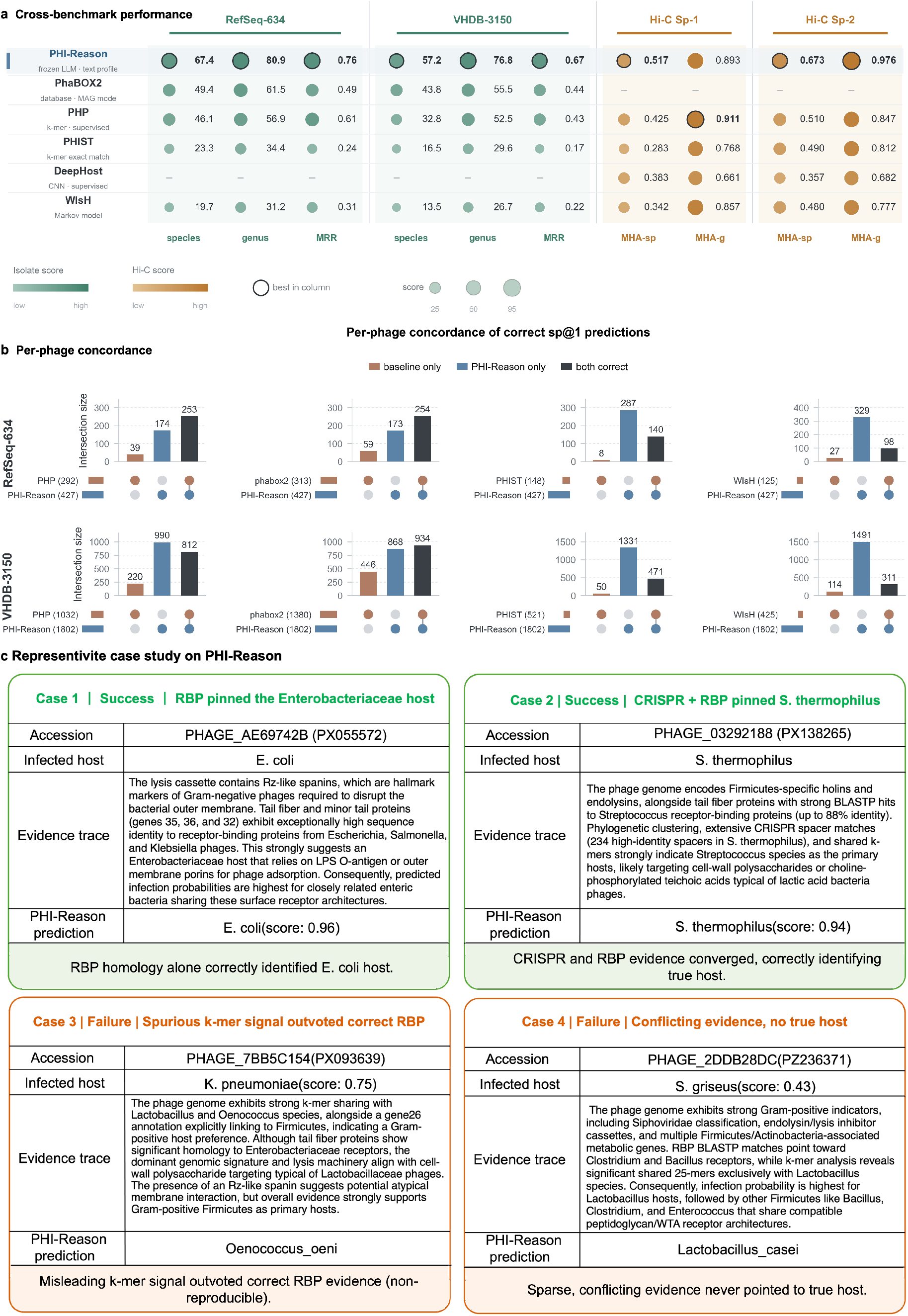
Structured profiles support portable host ranking across benchmarks. **a**, Cross-benchmark performance on RefSeq-634 (*n* = 634), VHDB-3150 (*n* = 3,150) and two Hi-C splits (Sp-1, *n* = 46; Sp-2, *n* = 82): sp@1, genus accuracy and MRR for the isolate benchmarks, and MHA-sp/MHA-g for Hi-C. Coverage in Supplementary Tables 1 and 19; 95% Wilson confidence intervals in Supplementary Table 1. **b**, Per-phage concordance with each baseline: PHI-Reason-only correct predictions exceed baseline-only in every comparison. **c**, Prospective case study (14 recently deposited genomes; Supplementary Note S12): two successes—*Escherichia* phage Henuyfy22N, where RBP homology alone identified *E. coli*, and *Streptococcus* phage P16, where concordant CRISPR and RBP evidence for *S. thermophilus* supported a correct species call—and two failures—*Klebsiella* phage Skif1059, where spurious alignment-free similarity to unrelated Firmicutes outweighed genuine Enterobacteriaceae RBP homology (not reproduced under repeated sampling), and the novel *Streptomyces* phage DeluluLabubu, where conflicting RBP and 25-mer signals produced a cross-genus error.

We included an embedding-only control to test whether the structured profiles could be used for host ranking without the language-model inference step. Using the same RefSeq-634 profiles encoded with qwen3-embedding:8B [17] and ranking candidates by cosine similarity, this training-free control reached sp@1 = 56.6%, compared with 67.4% for full profile listwise inference, a difference of 10.8 percentage points. The structured profiles therefore contained substantial host-discriminative information independently of the listwise inference procedure, whereas the evaluated global embedding-similarity formulation did not match candidate-conditioned language-model inference. This result does not establish language-model superiority over matched-access evidence-fusion or deterministic retrieval methods operating on the same records.

On the Hi-C cohorts, PHI-Reason achieved MHA-sp = 0.517 on Sp-1, compared with 0.425 for PHP, 0.383 for DeepHost, 0.342 for WIsH and 0.283 for PHIST. On Sp-2, PHI-Reason achieved MHA-sp = 0.673, compared with 0.510 for PHP, 0.490 for PHIST, 0.480 for WIsH and 0.357 for DeepHost (Fig. 4a; Supplementary Table 19). Given the modest cohort sizes and differences in the composition of the supplied evidence, these results should be interpreted cautiously rather than attributed to any single evidence source. Performance relative to the strongest baseline also varied across RefSeq-634 host families, with positive, small and negative margins depending on lineage. This variation is consistent with species-level performance depending on taxonomic coverage and evidence specificity rather than improving uniformly across host groups (Supplementary Table 11).

Per-phage concordance showed that PHI-Reason and the reference methods recovered overlapping but non-identical subsets of phages (Fig. 4b; Supplementary Table 20). On RefSeq-634, PHI-Reason correctly assigned 427 phages. The number recovered by PHI-Reason but missed by a given reference method exceeded the converse count for PhaBOX2 (173 versus 59), PHP (174 versus 39) and WIsH (329 versus 27). The same qualitative pattern was observed on VHDB-3150, although PhaBOX2 retained a subset of phages that was correctly assigned only by that method. These discordant prediction sets suggest potential complementarity between the methods, although the asymmetry in the counts is also partly attributable to the higher aggregate accuracy of PHI-Reason.

The cross-benchmark analyses indicate that structured PHI profiles carry species-level host information that can be used by frozen-language-model listwise inference across isolate-derived and metagenomic evidence regimes. The magnitude of the observed differences and the associated error patterns varied with prediction coverage, evidence availability and host-family composition and therefore do not imply uniform superiority across all retrieval or evidence-fusion settings. While these benchmarks established the framework’s performance across diverse phage–host evidence regimes, the true test of its generality lies in its ability to port to fundamentally different biological interaction tasks, which we examine in Section 2.8.

Finally, as a temporal evidence-attribution control, we applied the identical frozen-model interface to 14 recently deposited GenBank bacteriophage genomes whose hosts fall within the RefSeq-634 candidate catalogue (Fig. 4c, Supplementary Note S12, Supplementary Table 27). Because the clean genomes were only recently deposited, memorisation of the specific phage identity is less plausible, so this small set tests whether host calls follow the supplied evidence rather than memorised phage identities. Stripping all structured evidence (parametric-only) reduced genus-level top-1 from 11/14 to 2/14 and species-level from 6/14 to 1/14 (wide intervals at this sample size), with the single residual species-correct call confined to a textbook host-phage pairing (*Enterococcus faecalis*) recoverable from pretraining priors alone. Individual cases illustrate the range of behaviour. For *Streptococcus* phage P16, concordant evidence, including 234 CRISPR spacer matches at 100% identity and BLASTP receptor-binding-protein homology, both pointing to *S. thermophilus*, supported the correct species call under both the full profile and the profile with receptor-binding-protein evidence withheld, whereas the parametric-only control, lacking this named evidence, fell to an unrelated Gram-positive genus. This case illustrates evidence-supported correctness under the supplied structured evidence. A second success, the *Escherichia* phage Henuyfy22N, was identified from receptor-binding-protein homology alone, without a labelled BLASTN neighbour or CRISPR record. Failures took different forms. The novel jumbo *Streptomyces* phage DeluluLabubu combined conflicting field-level signals, with receptor-binding-protein homology pointing to *Clostridium* and *Bacillus* while alignment-free 25-mer sharing pointed exclusively to *Lactobacillus*, and the model followed the 25-mer signal to an incorrect cross-genus assignment under every evidence condition. The *Klebsiella* phage Skif1059 illustrated a distinct failure mode: despite tail-fibre homology to Enterobacteriaceae receptors consistent with the true host, the model over-weighted spurious alignment-free similarity to unrelated Firmicutes genera and misassigned even the host’s Gram status, remaining incorrect under every evidence condition. These prospective cases show that evidence-supported correctness and evidence-conditioned failure persist on recently deposited genomes, extending the dependence on named retrievable evidence documented on the benchmark. They remain illustrative rather than a powered accuracy estimate (full 14-genome results, provenance and deposition dates in Supplementary Note S12).

### 2.5 Post-cutoff evaluation on newly reported experimental phage–host associations

To evaluate PHI-Reason beyond the conventional benchmark setting, we assembled a post-cutoff test set of 207 experimentally supported phage–host associations reported in six studies published in 2025–2026 [18–23]. These associations were used exclusively for evaluation and were not included in the phage–host reference corpus. For this test, the reference corpus was constructed by integrating the RefSeq, VHDB and Hi-C datasets described above, while all auxiliary evidence resources, evidence-generation procedures, prompts and model settings were kept unchanged. Each query phage was processed through the same production pipeline and ranked against the fixed catalogue of 494 candidate host species. This setting therefore tests whether a frozen structured-evidence inference framework can use previously established biological knowledge to resolve newly reported phage–host associations.

PHI-Reason achieved species-level top-1, top-5 and top-10 accuracies of 52.2%, 82.1% and 91.8%, respectively, while genus-level top-1 accuracy reached 90.8% (Supplementary Note S13; Supplementary Tables 31–32). Thus, the large majority of post-cutoff phages were assigned to the correct host genus, whereas the remaining uncertainty was concentrated primarily in discriminating among closely related species within that genus. The substantial gap between genus- and species-level accuracy indicates that general host-lineage recovery transferred well to newly reported cases, while fine-grained species resolution remained more challenging.

We next used evidence perturbation to determine what limited performance in this post-cutoff setting. Keeping the model, prompt, candidate catalogue and reference corpus fixed, we removed individual evidence channels from the structured profiles. All perturbation conditions were scored over the complete set of 207 queries, with a query counted as incorrect whenever the confirmed host was not returned at rank 1, so that every condition is directly comparable to the full-profile accuracy reported above. Removing all specialized structured evidence reduced species-level top-1 accuracy from 52.2% to 18.8%. Removing BLASTN and CRISPR produced the largest decreases, reducing top-1 accuracy by 17.4 and 13.5 percentage points, respectively, followed by RBP removal (5.8 points), whereas removing the 25-mer channel increased accuracy by 1.9 points. These results show that performance on newly reported associations remained strongly dependent on retrievable biological evidence, while the informativeness of individual evidence channels differed substantially.

Additional label-masking and directed counterfactual tests showed that this dependence was not reducible to deterministic transfer of retrieved host identities. Masking BLASTN host labels was less disruptive than removing the complete BLASTN evidence block, accounting for 9.2 of the 17.4 percentage points lost on removal, and none of 185 evaluable predictions switched to an introduced same-genus or same-family decoy (Supplementary Note S13). Together, these results show that PHI-Reason can transfer a fixed structured-evidence inference framework to newly reported phage–host associations without incorporating those associations into its reference corpus. The principal residual difficulty lies at species-level resolution, where prediction increasingly depends on whether the available evidence is sufficiently specific to distinguish closely related candidate hosts.

### 2.6 Sufficiently capable backbones make effective use of structured profiles, while shared errors track evidence limitations

On RefSeq-634, we fixed the structured-profile schema, prompt, candidate-host catalogue and output requirements and varied only the language-model backbone and thinking mode. We compared prediction accuracy and concordance across configurations, quantified the evidence fields cited in generated rationales and examined the overlap between cross-backbone and cross-method errors. These analyses distinguish variation associated with model configuration from difficulty associated with the supplied evidence, but remain descriptive and do not isolate a single causal factor.

Host-prediction accuracy differed primarily between Qwen3-4B and the higher-capacity configurations evaluated here [17, 24–26]. Qwen3-4B reached sp@1 = 48.0% (95% confidence interval (CI), 44.1–51.8%; Wilson score interval) without thinking and 39.0% (35.2–42.8%) with thinking. GPT-oss-120B without thinking and GPT-oss-120B with thinking reached 63.3% and 65.8%, respectively (Fig. 5a; Supplementary Table 4). The deployed Qwen3.6:27b achieved the highest observed sp@1 among the tested configurations, reaching 67.4% (95% CI, 63.6–70.9%) without thinking and 67.7% with thinking, a difference of 0.3 pp despite a 5.3-fold increase in mean output length (Supplementary Table 4). Qwen3-Coder-Next reached 63.6% (59.7–67.2%), within the same higher-capacity tier as GPT-oss-120B but below Qwen3.6:27b, spanning 4.4 pp across these three higher-capacity backbones. Across the 634 phages, 81.4% were either correctly assigned by all five primary higher-capacity configurations— Qwen3.6:27b and GPT-oss-120B in both modes, plus Qwen3-Coder-Next—or missed by all five, with only small pairwise differences (McNemar tests; per-phage backbone predictions in Source Data). The observed separation therefore mainly distinguished Qwen3-4B from the higher-capacity configurations and does not define a universal or monotonic parameter-scale threshold; notably, the smallest of the higher-capacity backbones by parameter count, the dense Qwen3.6:27b, achieved the highest accuracy.

**Fig. 5.**
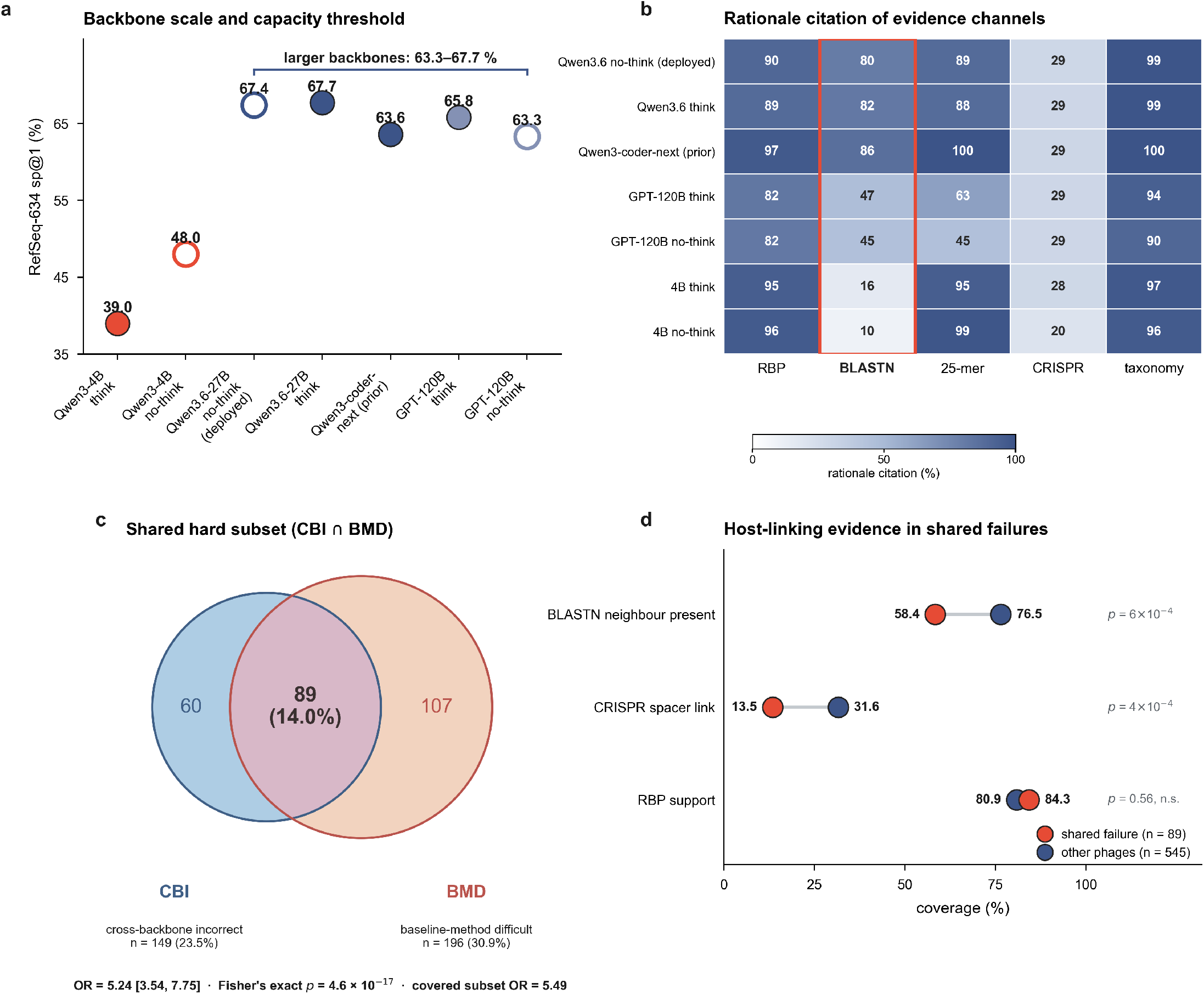
Model scale determines effective profile use; shared errors track evidence limitations. **a**, Backbone scale: the 4B configurations were markedly lower, while the larger backbones clustered at 63.3–67.7% sp@1 (Qwen3.6-27B highest at 67.4–67.7%, with no reliable gain from thinking). **b**, Rationale evidence-channel citation across seven backbone–mode configurations; the BLASTN column (red) separates backbones most strongly (86.4% for Qwen3-Coder-Next versus 9.5–15.5% for Qwen3-4B). Citations are descriptive, not causal. **c**, Shared hard subset: the cross-backbone-incorrect (CBI, *n* = 149) and baseline-method-difficult (BMD, *n* = 196) sets overlap in 89 phages (OR 5.24, 95% CI 3.54–7.75; two-sided Fisher’s exact *p* = 4.6 *×* 10*^−^*^17^). **d**, Host-linking evidence is depleted in shared failures (*n* = 89) versus the rest (*n* = 545): BLASTN neighbour 58.4% versus 76.5% (*p* = 6 *×* 10*^−^*^4^) and CRISPR 13.5% versus 31.6% (*p* = 4 *×* 10*^−^*^4^), both two-sided Fisher’s exact, whereas RBP does not differ.

The evidence fields cited in generated rationales varied more substantially across configurations than aggregate prediction accuracy. BLASTN showed the largest difference: Qwen3-Coder-Next cited BLASTN evidence in 86.4% of rationales and Qwen3.6-27B in 80.4%–82.0%, compared with 45.3%–47.0% for GPT-oss-120B and 9.5%–15.5% for Qwen3-4B (Supplementary Note S3; Fig. 5b). Despite these differences, the three primary higher-capacity configurations had similar aggregate accuracy and highly concordant predictions. Rationale citations therefore characterize the generated explanation and reporting style but do not establish that the cited evidence was causally responsible for the prediction.

Thinking mode increased output length without producing a consistent improvement in accuracy. For GPT-oss-120B, mean output length increased from 1,624 to 7,731 tokens, whereas sp@1 increased from 63.3% to 65.8% (+2.5 pp). For Qwen3-4B, output length increased from 1,012 to 3,497 tokens while sp@1 decreased from 48.0% to 39.0% (−9.0 pp). For Qwen3.6-27B, mean output length increased from 1,199 to 6,378 tokens (5.3-fold), with only a 0.3-pp change in sp@1 (67.4% to 67.7%; Supplementary Fig. 5 and Supplementary Note S4). Token count measures generated output length rather than total computational cost, but within these configurations longer generation was not monotonically associated with higher accuracy.

Backbone-invariant errors were enriched among phages that were also difficult for conventional PHI methods. The cross-backbone-incorrect set comprised 149 phages missed by all seven backbone–mode configurations, whereas the baseline-method-difficult set comprised 196 phages missed by PHP, PhaBOX2, PHIST and WIsH, with abstentions counted as errors. The two sets overlapped in 89 phages, corresponding to a 1.93-fold enrichment over the overlap expected under independence (odds ratio = 5.24; 95% CI, 3.54–7.75; two-sided Fisher’s exact test, *P* = 4.6 × 10*^−^*^17^; Fig. 5c). The enrichment persisted after restricting the analysis to the 532 phages for which all baseline methods returned valid predictions (odds ratio = 5.49; *P* = 2.3 × 10*^−^*^15^), indicating that baseline abstentions did not account for the overlap. This association does not, however, establish that the language-model and conventional methods failed through the same mechanism.

Shared failures were also associated with reduced availability of selected host-linking evidence. A labelled BLASTN neighbour was available for 58.4% of shared-failure phages, compared with 76.5% of the remaining phages (difference, −18.1 pp; 95% CI, −29.0 to −7.7 pp; *P* = 6 × 10*^−^*^4^). CRISPR-spacer links were available for 13.5% of shared-failure phages and 31.6% of the remaining phages (difference, −18.1 pp; 95% CI, −25.0 to −8.7 pp; *P* = 4 × 10*^−^*^4^). By contrast, RBP coverage did not differ detectably between the two groups (84.3% versus 80.9%; difference, +3.4 pp; 95% CI, −6.1 to 10.4 pp; *P* = 0.56). All comparisons used two-sided Fisher’s exact tests with Newcombe confidence intervals (Fig. 5d; Supplementary Table 8). These coverage measures quantify evidence availability rather than correctness or specificity, and the observed associations are not evidence of a causal effect. They are nevertheless consistent with the reference-coverage operating envelope observed across benchmarks in Section 2.4.

The model comparisons separate two sources of performance limitation. The smallest backbone converted the structured profiles into correct host assignments less effectively, whereas predictions from the larger-capacity configurations were highly concordant despite substantial differences in their generated citations and output lengths. Errors shared across model configurations and conventional methods were concentrated among phages with reduced BLASTN or CRISPR coverage and among profiles containing limited or conflicting support for the confirmed host. In a retrospective evidence-concordance stratification, profiles in which no evidence channel supported the confirmed genus remained below 15% genus-level accuracy across the evaluated configurations (*n* = 64), whereas profiles containing concordant support reached 98%–100% accuracy for the larger backbones (*n* = 198). Model capacity therefore affected how effectively structured profiles were converted into predictions, but evidence availability, correctness, concordance and specificity defined the remaining operating limits.

### 2.7 Pairwise scoring extends the structured-profile interface to nominated phage–host pairs

In addition to listwise ranking over a candidate catalogue, the structured-profile interface supports pairwise assessment of nominated phage–host hypotheses. The benchmark comprised 3,804 pairs from 634 phages, with one confirmed positive host and five fixed negative candidates for each phage, yielding 634 positive and 3,170 negative pairs and a positive prevalence of 0.167. The full profile retained BLASTN-

, CRISPR-, 25-mer- and RBP-derived host-associated evidence, whereas the binary pair label was used only to construct and evaluate the benchmark and was not supplied to the model. Only 1 of the 3,170 negative pairs belonged to the same genus as the confirmed host. The benchmark therefore primarily measures discrimination of a confirmed host from predominantly cross-genus candidates and does not provide a stringent test of within-genus, near-neighbour or fine species-level discrimination.

Pairwise scores strongly separated confirmed hosts from the negative candidates (Fig. 6a). The area under the precision–recall curve (PR-AUC) was 0.962, and the area under the receiver operating characteristic curve (ROC-AUC) was 0.992. Positive pairs received a mean support score of 0.847, compared with 0.026 for negative pairs (Cohen’s *d* = 6.10). When the six candidates associated with each phage were ranked by their pairwise scores, Hits@1 was 0.972, with 3.2% of phages producing a tie for the highest score.

**Fig. 6.**
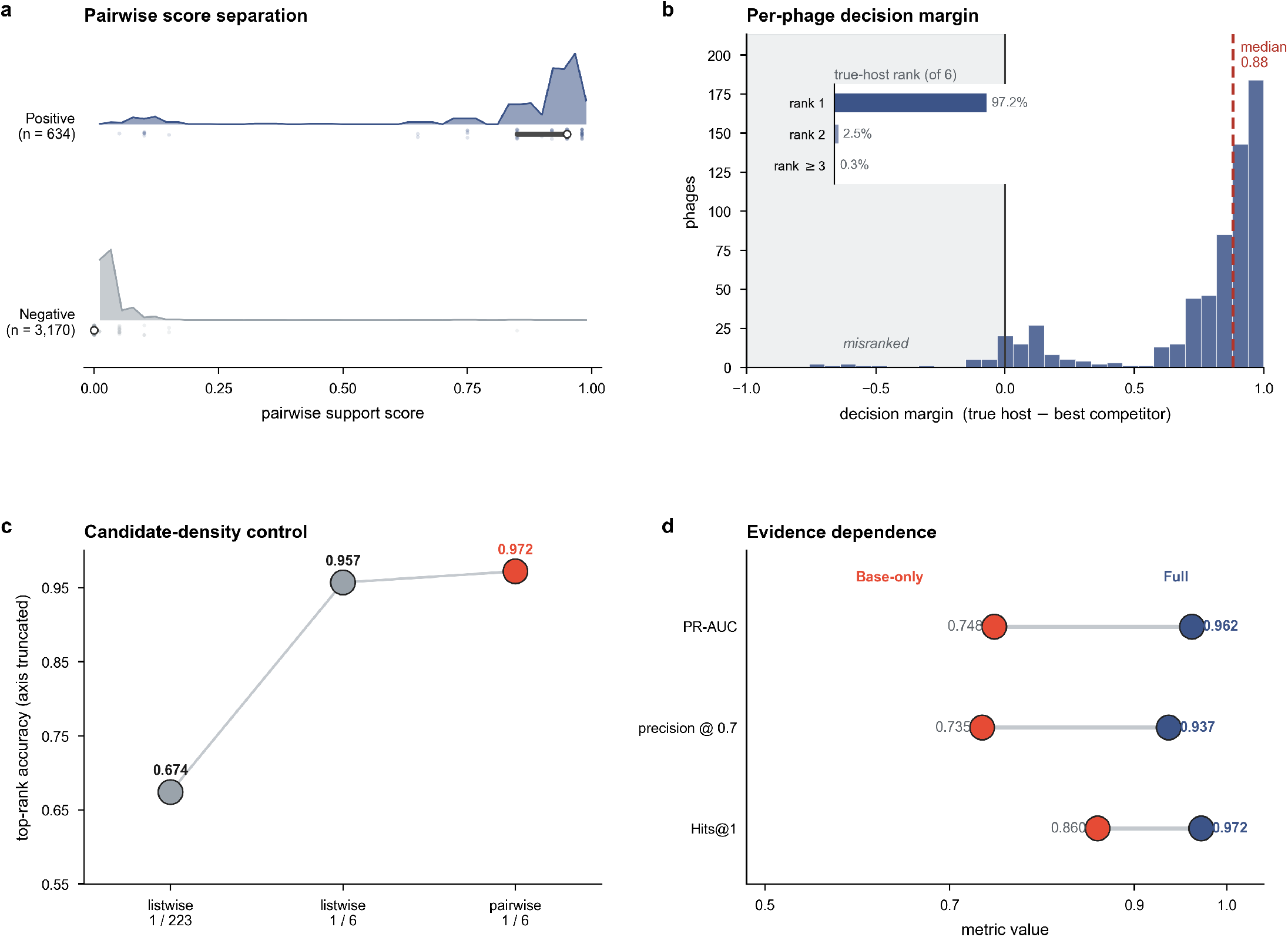
Pairwise scoring extends the structured-profile interface to nominated phage–host pairs. **a**, Support-score separation between 634 confirmed pairs and 3,170 genus-balanced negatives: PR-AUC 0.962, ROC-AUC 0.992, Cohen’s *d* 6.10. Negatives are operational alternatives, predominantly cross-genus. **b**, Per-phage margin (confirmed-host minus best negative score) positive for 94.0% of phages; the confirmed host ranked first or tied for first for 97.2% (strict first-only not recomputed for this backbone). **c**, Candidate-density control: restricting the list to the same six candidates raised sp@1 from 0.674 to 0.957, comparable to pairwise Hits@1 0.972. **d**, Evidence dependence: Base-only versus Full reduced PR-AUC (0.748 versus 0.962), precision at 0.7 (0.735 versus 0.937) and separation (Cohen’s *d* 2.70 versus 6.10).

The continuous score also supported prioritization within the nominated candidate pool (Fig. 6b). At a threshold of 0.7, the full profile reached a precision of 0.937, and empirical precision tended to increase at higher score thresholds (Supplementary Fig. 6). The score should not, however, be interpreted as a calibrated probability of phage–host interaction. Both PR-AUC and empirical precision depend on the constructed 1:5 positive-to-negative ratio, and their values therefore characterize prioritization within this benchmark rather than the precision expected under real-world PHI prevalence or a differently constructed candidate pool.

The high top-rank performance was largely attributable to the size and composition of the candidate sets rather than to an intrinsic advantage of pairwise inference. Restricting listwise inference from the complete 223-species catalogue to the same six candidates used in the pairwise benchmark increased species-level top-1 accuracy from 67.4% to 95.7%, close to the pairwise Hits@1 of 97.2% (Fig. 6c). Thus, the apparent improvement over full-catalogue ranking was explained primarily by the smaller and predominantly cross-genus candidate set, with no evidence that pairwise scoring outperformed listwise inference under matched candidate conditions. Pairwise performance consequently remained subject to the same candidate- and reference-conditioned operating limits observed for listwise inference in Section 2.4.

A Base-profile ablation showed that the specialized evidence fields contributed substantial discriminatory information beyond that retained in candidate-host names and genome annotations. Replacing the full profile with the Base profile reduced PRAUC from 0.962 to 0.748, precision at a score threshold of 0.7 from 0.937 to 0.735 and positive-negative separation from Cohen’s *d* = 6.10 to *d* = 2.70 (Fig. 6d). These results indicate that removing specialized evidence fields substantially weakens pairwise discrimination, while leaving a residual signal that may reflect information in the genome annotations, candidate-host identities or associations available from language-model pretraining. Pairwise assessment is therefore best viewed as a shortlist-ranking and candidate-triage mode for hypotheses nominated through upstream retrieval, experimental context or expert selection. It is not an open-world host-discovery procedure, a calibrated estimate of interaction probability or a validated test of discrimination among closely related host species.

### 2.8 Structured profiles transfer to eukaryotic virus–host prediction without architectural changes

To test whether the structured-evidence interface is restricted to phage–host prediction or is a portable contract for biological inference, we instantiated the same profile–prompt framework on the EvoMIL 36-host benchmark [27], which evaluates host assignment for eukaryotic viruses across 36 distinct host taxa. Because the biological mechanisms of eukaryotic virus entry, immune evasion and tropism differ fundamentally from phage–host interactions, the phage-specific evidence fields (CRISPR spacer matches, tail-fibre RBP homology, alignment-free 25-mer context) were replaced with domain-appropriate eukaryotic evidence: Baltimore classification, genome segmentation, taxonomic placement and BLASTP protein-homology neighbours (Supplementary Table 13). Critically, the profile–prompt contract, frozen model backbone, decoding settings and candidate-ranking procedure were held identical to the phage–host experiments; only the domain-specific evidence schema was adapted to the new biological domain.

Under this cross-domain transfer, PHI-Reason reached any-hit top-1 accuracies of 67.0% and 66.6% with gpt-oss-120B and Qwen3-Coder-Next, respectively, on the shared 3,186-virus evaluation set, exceeding EvoMIL (ESM-1b with multiple-instance learning, 60.4%), multi-neighbour homology-retrieval kNN (63.4%), and nucleotide-language-model and composition baselines (57.1% and 58.8%). Qwen3.6:27b reached 67.0% on the full 3,198-virus set; because this result uses a different denominator, it is reported separately rather than as a strictly matched comparison (Supplementary Note S10; Supplementary Tables 12–15). The matched-set advantage with gpt-oss-120B and Qwen3-Coder-Next indicates that the outcome was not specific to a single language model. Under the minimal-evidence floor condition, Qwen3.6:27b reached an any-hit top-1 accuracy of 19.2%, substantially below its performance with the full domain-appropriate evidence schema, further indicating that prediction performance depended strongly on the supplied structured evidence rather than on unaided model priors alone (Supplementary Note S10). As in the phage-host benchmarks, performance was strongest where informative domain evidence was available, while the marginal contribution and redundancy of individual fields shifted with the taxonomic coverage and mechanistic resolution of the eukaryotic resources.

These transfer results indicate that the predictive value of PHI-Reason does not depend on domain-specific parameter fitting or memorized phage sequences: with the inference layer frozen and only the evidence modules swapped, the framework integrates the available domain evidence and outperforms the task-trained baselines. The structured-profile contract is therefore portable across biological domains — the inference layer remains frozen and architecture-agnostic, while domain-specific biological knowledge is injected through swappable evidence modules. Because the framework is designed to integrate this evidence, its predictive performance in each domain reflects the coverage, correctness and resolution of the records made available to it, and it provides a unified, perturbation-ready platform for evaluating and combining these records across diverse interaction-prediction tasks.

### 2.9 Structured profiles transfer to a typed-decision inference engine for high-throughput inference

A central premise of the structured-evidence framework is that evidence construction is separated from inference, allowing the same structured profile to be paired with different inference engines. Having established portability across biological domains in Section 2.8, we next asked whether the same evidence interface could also transfer across inference architectures. We therefore replaced the Qwen3.6:27b language-model inference engine with Jev (TypeSafe AI) [28, 29], a commercial typed-decision inference system that returns structured probabilistic judgments over a candidate set rather than a generated natural-language rationale. The unmodified RefSeq-634, VHDB-3150 and HiC profiles were supplied to Jev, with candidate hosts specified through its choice primitive without changing the underlying evidence content, profile schema or candidate catalogue. This comparison therefore tests whether the predictive information exposed by the structured profiles can be used by a qualitatively different inference engine, rather than whether host ranking itself depends on generated reasoning.

Species-level predictive performance was broadly comparable between the typed-decision interface and Qwen3.6:27b (Supplementary Table 28). On RefSeq-634, Jev reached sp@1 = 68.3% (95% CI, 64.6–71.8%; Wilson score interval; *n* = 634), compared with 67.4% (63.6–70.9%) for Qwen3.6:27b. On VHDB-3150, Jev reached 54.8% (53.0–56.5%; *n* = 3,150), with seven provider-side token-limit errors scored as incorrect under the abstention-as-error policy, compared with 57.2% for Qwen3.6:27b. On the HiC cohorts, evaluated using the species-level multi-host accuracy adopted throughout this study, Jev reached MHA-sp = 0.483 on Sp-1 and 0.643 on Sp-2 under the noul primitive and 0.375 and 0.622 under choice, compared with 0.517 and 0.673 for Qwen3.6:27b; the corresponding any-hit sp@1 values were 50.0% and 68.3% versus 52.2% and 68.3% (Supplementary Table 29). Thus, across all three benchmarks, replacing the language-model inference engine with a typed-decision engine preserved broadly similar predictive performance despite the absence of generated rationales. This supports the interpretation that a substantial fraction of the host-discriminative information resides in the structured evidence interface itself and can be extracted by more than one class of inference architecture.

This architectural transfer also substantially reduced inference time. On RefSeq-634 and HiC, where the candidate catalogue fits within a single choice call, mean per-query inference time under the typed-decision engine was 0.69 s and 0.60 s, respectively, compared with 27.8 s and 19.7 s for Qwen3.6:27b, corresponding to approximately 40× and 33× shorter inference times. On VHDB-3150, Jev’s per-call candidate limit required a three-call elimination procedure for each query, whereas Qwen3.6:27b ranked all 488 candidates in a single call. Even under this less favourable setting, mean per-query time was 2.23 s (summed across the three calls), compared with 23.7 s for Qwen3.6:27b, corresponding to an approximately 11× reduction (Supplementary Table 30). All times are wall-clock means measured under matched serial, single-query execution and include network round-trip for the remote typed-decision service; full methodological detail is given in Supplementary Table 30.

These results extend the portability of the structured-evidence interface along a second axis. The eukaryotic virus–host analysis demonstrates transfer across biological domains by replacing domain-specific evidence fields while retaining the inference framework, whereas the Jev analysis demonstrates transfer across inference architectures by retaining the structured evidence while replacing the language-model inference engine with a typed-decision inference engine. The framework therefore supports complementary operating modes: language-model inference when natural-language rationales are required for evidence-grounding and error-partition analyses, and typed-decision inference when only ranked predictions are required and throughput is prioritized. The latter does not replace the former for analyses that depend on generated rationales, but shows that the structured profiles are not intrinsically coupled to any particular language model or inference architecture.

## 3 Discussion

PHI-Reason reframes biological interaction prediction as inference over an explicit, structured evidence interface. Rather than treating heterogeneous evidence only as input features to be compressed into a score or latent representation, the framework preserves genome annotations, reference-neighbour context, receptor-binding-protein homology, host surface features and defence-related information as named and independently manipulable records. A frozen general-purpose language model then integrates these records at inference time without domain-specific parameter fitting. The resulting system is therefore not merely a predictor: it is an experimental interface through which evidence contribution, redundancy, corruption, uncertainty and failure can be directly measured under a fixed model and decision setting.

This formulation marks a conceptual shift from recent language-model applications in biology. Protein and DNA foundation models primarily encode sequence regularities into numerical embeddings, while hybrid systems such as BioReason [11] and BioReason-Pro [12] connect learned biological representations to language-model reasoning. ProLLM similarly formulates protein–protein interaction prediction as a text-to-text task using pathway-structured prompts [30]. A critical limitation of these approaches is that the evidence is fused or encoded before the final prediction, making it difficult to isolate the contribution of individual biological records. In contrast to standard retrieval-augmented generation (RAG), which simply appends context snippets, our structured-profile paradigm treats the *identity and organization* of heterogeneous evidence as part of the inference problem itself. By rendering biologically distinct signals as explicit, named fields, the framework makes it possible to intervene on evidence–label relationships and test the model’s dependence on them, rather than interpreting model outputs only after evidence has been irreversibly fused.

The perturbation experiments reveal that the utility of a biological evidence source is conditional on its coverage, specificity, correctness and relationship to other fields. Nucleotide-level reference context provided the strongest individual signal, but its contribution depended on the availability and correctness of host-labelled neighbours. RBP homology supplied complementary adsorption-associated information, whereas CRISPR and alignment-free 25-mer evidence differed in coverage and marginal contribution. Crucially, a field could carry substantial signal in isolation yet add little once more specific evidence was already present, demonstrating that evidence utility cannot be inferred from single-field performance alone. Conversely, corrupting evidence–host labels could be more damaging than removing the corresponding field, showing that an informative but incorrectly associated record can actively redirect prediction. These findings shift the emphasis from asking which evidence type is universally strongest to asking under which reference, taxonomic and candidate conditions each evidence description becomes discriminative, redundant or misleading.

The same pattern links evidence perturbation to model-capacity and shared-failure analyses. Backbones with sufficient capacity used the structured profiles more effectively than the smallest evaluated model, but performance did not scale monotonically with parameter count among the larger configurations: the deployed dense Qwen3.6:27b matched or exceeded the substantially larger GPT-oss-120B. Extended thinking traces substantially increased output length and computational cost without producing a consistent accuracy gain. Thus, effective profile use appears to require a sufficiently capable backbone rather than simply a larger one, and not necessarily prolonged explicit reasoning. More importantly, errors shared across language-model backbones were enriched among queries also missed by conventional methods and among profiles with reduced or conflicting host-linking evidence. Once the backbone was sufficiently large, the dominant limitation shifted from model capacity to the coverage, correctness and concordance of the supplied evidence. A stronger language model could integrate available evidence more effectively, but it could not reliably recover a host that was not represented by decision-relevant evidence. The post-cutoff evaluation further supports this evidence-dependent interpretation: PHI-Reason resolved newly reported phage–host associations without task-specific retraining, while removing specialized evidence substantially reduced accuracy. The gap between genus- and species-level performance further indicates that generalization depends not only on evidence availability, but also on its taxonomic resolution.

The true test of this paradigm is its portability beyond the phage–host domain. To validate the domain-agnostic nature of the structured-profile contract, we instantiated the same interface on the EvoMIL eukaryotic virus–host benchmark, substituting phage-specific fields with domain-appropriate evidence. PHI-Reason reached 66.6– 67.0% any-hit top-1 accuracy, demonstrating that the structured-profile contract can transfer to a distinct virus–host setting while remaining strongly conditioned on homolog-neighbour support (Supplementary Note S10). The transferable component is therefore the organization of evidence into a common inference interface, whereas predictive performance remains determined by the quality and resolution of the evidence available in each biological domain. More generally, conditioning a frozen model on named, perturbable evidence records provides a portable contract for evidence-driven biological prediction: evidence modules can be inspected, perturbed and compared before model fitting or deployment, separating questions of evidence coverage from questions about the inference layer. This portability also extends to the inference architecture. Replacing the generative backbone with the Jev typed-decision engine preserved broadly comparable predictive performance while reducing inference time by 11–40-fold. Together, these experiments suggest that the structured evidence interface is transferable across both biological domains and inference engines, rather than being tied to a particular model architecture.

The evidence interface also clarifies the relationship among hallucination, prediction error and uncertainty. A rationale can remain faithful to the supplied profile while the final prediction is incorrect, because the profile itself may be incomplete, conflicting or non-decisive. Conversely, unsupported explanatory statements can accompany an otherwise correct prediction. PHI-Reason therefore separates claim-level grounding, rationale-linked hallucination and answer-level departure from the supplied evidence instead of treating all errors as hallucinations. Most incorrect predictions remained anchored to one or more supplied host-associated signals and arose when the confirmed host was absent from the profile or when several evidence-supported candidates remained plausible. The dominant failure mode was therefore evidence-bounded misranking rather than unconstrained host invention. This distinction has broader implications for scientific language-model evaluation: grounding measures whether an output stays within the supplied evidence, whereas correctness additionally depends on whether that evidence is sufficiently complete and discriminative to resolve the task.

A critical insight from our framework is that predictive performance is fundamentally bounded by the *resolution* of the available evidence. Our current implementation is scoped to species-level prediction because the reference signals (homology, composition, and defence records) are primarily organized at the species or genus level. Strain-level discrimination, by contrast, requires isolate-level facts—such as receptor-inactivating mutations, exact CRISPR spacer–protospacer matches, or specific capsule profiles [31–33]—that are frequently sparse or absent from current annotations. When such fine-grained evidence is missing, a model may fill the gap with plausible but unsupported mechanisms, producing the same evidence-conditioned hallucinations we observed at the species level. Reliable strain-resolved prediction would therefore require not a different model, but a deeper and more standardized evidence regime. PHI-Reason is thus best positioned as a species-resolution prioritisation and evidence-inspection framework, whose value for finer-grained work lies in making explicit which isolate-specific evidence a confident prediction would require.

In conclusion, PHI-Reason establishes a profile-based paradigm for biological interaction prediction in which heterogeneous evidence is retained as modular, human-readable and experimentally perturbable records. The framework itself is reference-agnostic: updated databases, catalogues and annotations can be incorporated as additional fields without retraining the frozen language model. Together with our embedding-only controls and perturbation analyses, these results show that predictive performance depends on the correctness and organization of evidence–label relationships, rather than on the mere presence of profile fields. Future improvements in biological prediction may therefore come not only from stronger models, but from evidence resources with broader coverage, finer taxonomic resolution and more reliable biological associations. More broadly, this study suggests a new practical role for general-purpose language models in scientific inference. Their value need not depend primarily on memorized domain knowledge or extended free-form reasoning. It can arise from their ability to compare heterogeneous records within an explicit and updateable evidence interface. In such systems, the fundamental unit of scientific inference is not only a sequence, feature vector or prompt, but an evidence record with a defined source, scope, resolution and relationship to competing hypotheses. By making these records explicit and experimentally manipulable, PHI-Reason converts the limits of prediction from hidden properties of a fitted model into measurable boundaries of the supplied evidence.

## 4 Methods

### 4.1 Profile generation

PHI-Reason represents phage genomes and candidate host genomes as structured textual profiles. Profiles were generated before benchmark inference using deterministic bioinformatics pipelines, curated lookup tables and predefined formatting rules. No language model was used during profile construction. Each phage profile contained gene-level functional annotations, sequence-derived protein descriptors, receptor-binding-protein evidence and optional nucleotide-neighbour (BLASTN), alignment-free 25-mer and CRISPR-spacer reference-context blocks. Each host profile contained surface-relevant proteins, anti-phage defence systems [34] and curated receptor, host-range and species-discriminating information. These fields were represented as named biological records so that they could be retained, removed, scrambled or recombined in downstream perturbation experiments.

#### 4.1.1 Phage genome profiles

For phage profiles, open reading frames were predicted from each genome using Prodigal v2.6.3 [35] and annotated with complementary phage and orthology resources, including PHROG v4 [36] searched with MMseqs2 13-45111+ds-2 [37], and eggNOG-mapper v2.1.13 [38] using the eggNOG v5.0.2 [39] database. PHROG product names were used as the primary standardized labels when available, with non-redundant eggNOG descriptions retained as qualifiers. For each annotated protein, sequence-derived descriptors including length, theoretical isoelectric point, molecular weight, hydrophobicity and an approximate transmembrane-helix count were computed and stored as profile fields. Annotated genes were grouped into functional sections relevant to host inference, including tail and host adsorption, lysis, head and DNA packaging, connector and head–tail junction, lysogeny, replication and metabolism, gene expression and regulation, and accessory host-derived factors. Genes without functional annotation were summarized as counts to retain information about hypothetical-gene burden without exceeding the inference context budget.

#### 4.1.2 Receptor-binding evidence

Receptor-binding evidence was generated by searching tail-associated or adsorption-related phage proteins against PHIStruct [40], a curated receptor-binding-protein corpus comprising 19,081 phage receptor-binding protein sequences across 238 host genera. The reference FASTA was indexed with DIAMOND v2.1.24 and queried using proteins annotated as tail-associated by PHROG or containing adsorption-related product keywords, including tail fibre, tail spike and receptor-binding protein. Candidate hits were generated with DIAMOND against a de-leaked receptor-binding-protein reference database at a minimum sequence identity of 40% (aligned with PHIStruct’s lowest similarity bin), a minimum query coverage of 5% and a maximum e-value of 10*^−^*^5^. During profile assembly, hits were aggregated by query gene and host genus, and up to five host genera per query gene were retained, ranked by a composite score pident^0.6^ qcov^0.4^.

For each query gene–host genus pair, the profile retained the best sequence identity, best query coverage and number of hits. The top host genera were then appended to the corresponding tail-gene entry in the phage profile. Species-level host ranking was based on the integration of this RBP-derived signal with other profile fields, including nucleotide-level (BLASTN) reference context, alignment-free 25-mer and CRISPR-spacer context, and candidate-host information. Detailed filtering thresholds, ranking scores and implementation details are provided in the released profile-generation scripts and source data.

#### 4.1.3 Reference-context blocks

Reference-context blocks were constructed by comparing query phages with a leave-test-out corpus of experimentally confirmed phage–host associations. Evaluation sequences were excluded from this corpus before profile generation; full dataset provenance, curation and leakage controls are detailed in Supplementary Note S9. Three host-labelled reference blocks were built. The nucleotide-neighbour (BLASTN) block was generated using NCBI BLAST+ v2.16.0 [41] and listed up to five qualifying BLASTN neighbours (percentage identity ≥ 70% and query coverage ≥ 3%), ranked by bitscore, together with their confirmed hosts. The alignment-free 25-mer block ranked reference phages by the number of shared canonical 25-mers (*k* = 25, PHIST-style) with the query genome and listed the top neighbours together with their confirmed hosts. The CRISPR block listed host CRISPR spacers matching the query phage genome at ≥ 90% identity over ≥ 90% of the spacer length with at most two mismatches, together with the corresponding host. Protein-level homology (DIAMOND v2.1.24 BLASTP [42]) was used only to identify receptor-binding-protein matches (above), not as a separate reference-context block. Reference phage accession identifiers were replaced by deterministic hashes before inference to reduce direct identifier use. Phages without qualifying neighbours in a given block received no corresponding reference-context block.

#### 4.1.4 Host genome profiles

Candidate host genomes were converted into structured profiles using three layers. First, predicted proteins were annotated and filtered for surface-associated functions using a curated lexicon covering outer-membrane proteins, porins, TonB-dependent receptors, lipopolysaccharide and O-antigen biosynthesis, capsular polysaccharide, peptidoglycan remodelling, flagella, type-IV pili, teichoic-acid biosynthesis, S-layer terms and related envelope features. Second, anti-phage defence systems were detected with DefenseFinder v2.0.1 [34] and summarized as a dedicated defence field. Third, each host was augmented with curated quick-profile fields describing taxonomy, Gram type, ecological context, primary receptors, known infecting phage families, species-discriminating features and host-range boundaries where available. These curated fields were compiled from primary literature and phage–host interaction resources before benchmark inference.

#### 4.1.5 Identifier masking

To reduce direct accession lookup and name-based shortcuts, query phage accessions, phage names, assembly identifiers and accession-like strings were replaced with deterministic opaque hash codes before inference. The same masking was applied to phage identifiers appearing in the BLASTN-, 25-mer- and CRISPR-derived reference-context blocks. Benchmark names, split labels and ground-truth host fields were not included in the inference prompts. Candidate host species names were retained because they define the prediction target, but no query-specific host label was provided outside the allowed reference-context evidence fields. As a sanity check against host-name-driven recall, the annotation-only Base condition retained the same candidate-host names but removed the specialised PHI evidence layers (RBP, BLASTN, CRISPR and 25-mer evidence); it achieved only 22.6% sp@1 on RefSeq-634, indicating that visible candidate-host names were insufficient to recover the correct assignments in the absence of specialised profile evidence.

#### 4.1.6 Host-list compression for inference

For host-list inference, candidate host profiles were compressed into a shared list placed in the same context window as the query phage profile. Each host was represented by a compact pipe-delimited line containing host name, habitat, receptor information and known infecting phage families, with missing fields marked as “-”. Hosts were grouped by broad envelope category (Gram-negative, Gram-positive or Archaea) and then by taxonomic family, preserving coarse cell-envelope and phylogenetic structure while allowing all candidate hosts to be ranked in a single language-model inference call.

### 4.2 Language-model inference and computational environment

PHI-Reason in species-level experiments used Qwen3.6:27b [26] served locally with Ollama v0.23.2 [43] in its no-thinking configuration, with internal reasoning disabled through the API-level think parameter rather than a text-prompt convention. Inference was performed with temperature 0.1, a 40,960-token context window and non-streaming generation. Unless otherwise stated, experiments used this no-thinking configuration; thinking-mode comparison runs (Section 2.6) enabled internal reasoning through the same API parameter and used a larger generation budget. Outputs were required to follow predefined JSON schemas and were retried up to twice if parsing failed. Successfully parsed phage-level or pair-level outputs were cached and reused in downstream analyses. Because the deployed predictor is a specific quantised build served at low but non-zero temperature, reported accuracies are specific to this model build and decoding configuration; the exact model file, quantisation and serving parameters are pinned in Supplementary Table 25 to support reproduction.

Two prompt formats were used. In the host-list setting, one inference call was made per query phage using the compressed 223-host catalogue, the query phage profile and fixed scoring instructions; the model returned the top 30 ranked candidate hosts. Hosts not returned in the top-30 output were assigned an implicit score of zero. In the pairwise setting, one inference call was made per nominated phage–host pair using one phage profile and one host profile; the model returned an infection-plausibility score and categorical decision for that pair independently of other hosts.

The main experiments were run on a local workstation with two NVIDIA RTX PRO 6000 Blackwell Max-Q GPUs and 96 GB GPU RAM for each GPU, using NVIDIA driver/CUDA versions 570.211.01/12.8. Flash attention was enabled, and model layers were offloaded to GPU memory through Ollama. GPU-hours were calculated as wall-clock runtime multiplied by the number of active GPUs, which was fixed at two for the main experiments. Full prompt templates, generation parameters and output schemas are provided in Supplementary Note S7.

#### Cross-backbone and reasoning-mode configurations

The cross-backbone comparison in Section 2.6 considered three additional open-weight language-model backbones served through the same local Ollama backend: Qwen3:4b-Q4 K M [17], gpt-oss:120b [25] and Qwen3-Coder-Next:Q4 K M [24]. Qwen3-4B, GPT-oss-120B and the deployed Qwen3.6-27B were each run in two configurations, with internal reasoning enabled and with internal reasoning suppressed; Qwen3-Coder-Next was run in its no-thinking configuration only. The same prompt formats, JSON schemas, host catalogue and phage profiles were used across backbone comparisons, so that the deliberate variables were model identity and reasoning-mode setting. The main-text comparison (Section 2.6) therefore spans seven backbone– mode configurations—Qwen3-4B and GPT-oss-120B with and without thinking, the deployed Qwen3.6-27B with and without thinking, plus Qwen3-Coder-Next (no-thinking only) (Supplementary Table 4).

Sampling temperature was fixed at 0.1 for all backbone comparisons, with a 40,960-token context window for no-thinking configurations and 49,152 tokens for thinking-mode configurations. To accommodate longer reasoning traces, the maximum generation budget was increased from 4,096 tokens (no-thinking) to up to 40,000 tokens for thinking-mode runs. Internal reasoning blocks were removed before JSON parsing. Mean generated-token counts for each backbone configuration are reported in Supplementary Table 4; hardware and the primary-run compute budget are given in Supplementary Table 25. All cross-backbone experiments used the same workstation and GPU-offload configuration as the primary PHI-Reason experiments.

#### Shared-failure and evidence-concordance sets

The cross-backbone-incorrect set (CBI) comprised phages predicted incorrectly by all seven backbone–mode configurations, and the baseline-method-difficult set (BMD) comprised phages predicted incorrectly by all four conventional baselines (PHP, PhaBOX2, PHIST and WIsH), with abstentions counted as errors; the enrichment of their overlap was tested against independence with a two-sided Fisher’s exact test over all 634 phages and repeated on the subset with a valid prediction from every baseline. For the evidence-concordance stratification, each profile was labelled retrospectively, using the confirmed host genus, according to whether its available evidence channels supported concordant or conflicting host genera and whether any channel supported the confirmed genus (definitions in Supplementary Note S5).

### 4.3 Evaluation benchmarks

#### Benchmark datasets

We evaluated PHI-Reason on three datasets that differ in curation procedure, host-label structure and biological domain: RefSeq-634, VHDB-3150 and a metagenomic HiC proximity-ligation benchmark.

##### RefSeq-634

RefSeq-634 is the held-out evaluation partition of the CHERRY-1940 dataset [7], which contains 1,940 phage–host pairs assembled from NCBI RefSeq [44] with species-level host assignments across 223 bacterial taxa. We partitioned this dataset by phage accession into a 1,306-phage reference partition and a 634-phage held-out evaluation partition. The 1,306-phage partition was used only as an explicit reference corpus for nucleotide-neighbour and alignment-free (25-mer) reference-context retrieval.

##### VHDB-3150

VHDB [15] is a larger RefSeq-derived benchmark comprising 4,698 phage–host pairs across 498 bacterial host species. To construct a benchmark consistent with the RefSeq-634 reference split, we used the 1,280 VHDB phages that overlapped with the CHERRY-1306 training partition as the training/reference partition. These phages were used as the labelled reference corpus for nucleotide-neighbour and alignment-free (25-mer) reference-context retrieval and for retraining applicable baseline methods. The remaining VHDB test phages were used for evaluation. Among these test phages, 268 did not have species-level ground truth suitable for species-level evaluation, leaving 3,150 phages with confirmed species-level host labels, ranked against a closed candidate universe of 488 host species; this subset is referred to as VHDB-3150.

##### HiC

The HiC benchmark is derived from MetaHiC proximity-ligation co-sequencing of human gut metagenomes. Viral contigs are linked to candidate host metagenome-assembled genomes (MAGs) through physical co-localisation evidence; MAG-level associations are then collapsed to species level using GTDB taxonomy, producing species-level phage–host associations across a 52-species host universe *H*. Unlike the isolate benchmarks, a single phage contig may be associated with multiple host species; therefore, the ground truth for each phage *q* is a set of host species *Y_q_* ⊆ *H* rather than a single label. We followed the evaluation protocol of [16], reporting results on two complementary splits: Sp-1 (*n* = 46 multi-host test phages) and Sp-2 (*n* = 82 host-number-stratified test phages). This benchmark provides a domain-shift evaluation because the candidate hosts are gut metagenomic species represented by selected MAG-derived reference genomes rather than RefSeq-derived isolate genomes.

#### Abstention-as-error policy

Unless otherwise stated, all evaluations used an abstention-as-error policy. If a method produced no prediction, returned a host outside the candidate space, or failed to assign a species-level host under the required output resolution, the case was counted as an incorrect prediction. This policy keeps denominators fixed across methods and prevents apparent performance gains from selective abstention on difficult cases.

### 4.4 Baseline methods

We compared PHI-Reason with five representative phage–host prediction methods: PHP, PhaBOX2, PHIST, WIsH and DeepHost. Together, these baselines cover composition-based probabilistic models, exact k-mer matching, hybrid multi-signal reference-based pipelines and supervised sequence models. Baseline methods were evaluated on a benchmark when their training requirements and output format were compatible with the corresponding evaluation task. For each benchmark, baseline predictions were mapped to the same benchmark-specific candidate host space used for PHI-Reason whenever possible. Predictions assigned to hosts outside the candidate set were counted as incorrect. Queries for which a method returned no host prediction were treated as abstentions and counted as errors.

#### PHP

is a host prediction method based on k-mer composition profiles and Gaussian mixture modelling. Query phages and candidate hosts were represented by k-mer frequency features, and candidate hosts were ranked according to the likelihood assigned by the fitted host models. PHP was rebuilt separately for each benchmark using the corresponding training partition and candidate host set. For RefSeq-634, PHP was trained using the CHERRY-1306 training partition. For VHDB-3150, PHP was trained using the VHDB training partition defined by overlap with the CHERRY training set. For the HiC benchmark, PHP was retrained using the corresponding training split and the 52 species-level candidate hosts, with each host species represented by one selected MAG-derived reference genome. Because PHP assigns scores to all candidate hosts, it produced a complete host ranking for every query phage.

#### PhaBOX2

is a hybrid multi-signal phage analysis framework that combines host prediction evidence from sequence-based comparisons, k-mer features and reference databases. We ran PhaBOX2 v2.2 in MAG mode using its complete default database for the RefSeq-634 and VHDB-3150 isolate-derived benchmarks. Reported host assignments were parsed, taxonomic prefixes were removed where present, and the resulting host names were normalized to the benchmark taxonomy before species-level evaluation. Predictions assigned to hosts outside the benchmark-specific candidate host space were counted as incorrect. Because PhaBOX2 returns sparse host assignments rather than a calibrated score or complete ranking over all candidate hosts, it was evaluated using its top-ranked species-level prediction. PhaBOX2 was not evaluated on the HiC benchmark because the HiC task required multi-host ranking over 52 species-level candidate hosts, each represented by one selected MAG-derived reference genome, whereas PhaBOX2 returns conservative sparse host assignments rather than complete candidate-host rankings.

#### PHIST

is an exact k-mer matching method that identifies statistically significant sequence similarity between query phage genomes and candidate host genomes. We ran PHIST with default parameters against the benchmark-specific host genome sets. Candidate hosts were ranked according to the reported significance scores, with more significant matches placed higher. Queries for which PHIST returned no significant host prediction were treated as abstentions and counted as errors in all reported metrics.

#### WIsH

is a host prediction method based on homogeneous Markov models of host nucleotide composition. For each benchmark, host-specific Markov models were built from the corresponding candidate host genomes using the WIsH build procedure. Query phage genomes were then scored against all candidate host models, and hosts were ranked by log-likelihood. WIsH was evaluated on RefSeq-634, VHDB-3150 and the HiC benchmark using the corresponding benchmark-specific candidate host sets. Because WIsH assigns a likelihood to each candidate host, it produced a complete ranking for every query phage.

#### DeepHost

is a supervised deep learning method that predicts host labels from one-hot encoded phage genome sequences using a convolutional neural network. As a closed-set classifier, DeepHost can only predict host classes observed during training. We therefore evaluated DeepHost only in settings where a compatible benchmark-specific host vocabulary could be defined for both training and testing. DeepHost was not evaluated on the RefSeq-derived isolate benchmarks because the host classes in the training and test partitions were not identical, leaving some test hosts outside the model’s prediction space. For the HiC benchmark, DeepHost was retrained on the training split using the 52 species-level host labels and evaluated on the corresponding held-out test split.

#### Profile-embedding retrieval control

To test whether the structured profiles could be exploited by a purely numerical procedure without the language-model ranking step, we added a training-free embedding-only control on RefSeq-634. Using the same phage and host profiles supplied to PHI-Reason, each profile was encoded with the text-embedding model qwen3-embedding:8B [17]. The phage and host embedding matrices were mean-centred separately, the vectors were L2-normalised, and candidate hosts were ranked for each query phage by cosine similarity over the 223-host catalogue. No labels, model training or language-model inference were used; species-level top-1 accuracy (sp@1) was the primary metric, providing a lower bound on the host-discriminative information carried by the profiles themselves.

### 4.5 Evidence perturbation experiments

Because PHI-Reason conditions the frozen model on named evidence records rather than latent feature vectors, the input evidence itself can be treated as a controlled experimental object: named fields and their host labels were added, withheld, degraded or relabelled while the language-model backbone, prompt format, candidate host list and evaluation metric were held fixed, isolating the contribution, redundancy and correctness-dependence of each source. These are descriptive input-level interventions in a closed-set, candidate-conditioned setting and are not causal manipulations of the model or of biology. All perturbation experiments were performed by modifying named fields in the input profiles before inference. No model parameters were updated, and each perturbation variant was evaluated with the same inference backend and decoding settings as the corresponding full profile condition.

For species-level host-list experiments, the Base condition was defined as an annotation-only phage profile. It retained query-genome-derived functional annotations, gene-level product labels, protein descriptors and taxonomy-derived annotation signals, but removed all specialised evidence layers. The four specialised layers were receptor-binding-protein (RBP) matches, BLASTN-derived whole-genome nucleotide-reference context, host CRISPR-spacer records and alignment-free 25-mer genome-composition similarity. Single-field conditions were generated by adding one specialised layer back to the Base profile; the full condition contained all four layers. Leave-one-field conditions removed a single layer from the full profile. Independent field gains were measured relative to the Base condition, and redundancy was quantified by comparing the observed full profile gain with the sum of the single-field gains and with the leave-one-field reductions. Gram-type and BLASTN-neighbour-availability stratifications partitioned RefSeq-634 by host Gram classification and by the presence of at least one labelled BLASTN neighbour in the supplied profile (the reference-context threshold of ≥ 70% identity and ≥ 3% query coverage), respectively; the shared-failure coverage analysis (Supplementary Table 8) used the same labelled-BLASTN-neighbour-presence flag on the shared-failure subset.

To test whether reference-context improvements reflected use of explicit host labels rather than similarity scores alone, we generated label-perturbation controls for the three host-labelled reference blocks (BLASTN, 25-mer and CRISPR), while retaining the RBP field and all non-label content. In the label-scrambled conditions, host labels in the corresponding block were permuted while preserving the retrieved reference entries, similarity scores, hit ordering and all non-label annotations; BLASTN-only, 25-mer-only, CRISPR-only and all-block scrambling were evaluated separately. In the label-free control, host labels were removed from all three host-labelled blocks while retaining the reference entries and similarity information. These controls distinguished the contribution of labelled reference associations from label-independent similarity information.

The same perturbation logic was applied to the pairwise evaluation experiments. Each phage–host pair was evaluated under the full profile and under matched perturbation variants in which specialised evidence layers were removed, isolated or label-scrambled. Pairwise perturbation results were assessed using PR-AUC, ROC-AUC, F1, MCC, per-phage Hits@1, log-loss and high-confidence precision. Because pairwise scoring evaluates individual phage–host hypotheses rather than global candidate ranking, these analyses were used to test whether the evidence layers identified in host-list ranking also supported reliable pair-level scores.

All perturbation outputs were parsed with the same JSON validation and retry rules as the primary experiments. Cached outputs were reused only when the input profile, prompt template, model setting and perturbation condition were identical. Numerical values for host-list perturbations are reported in Supplementary Tables 2 and 6, and pairwise perturbation results are reported in Supplementary Table 9.

### 4.6 Pairwise evaluation construction and host-list comparison

Pairwise evaluation experiments were constructed from RefSeq-634. For each phage, the experimentally confirmed host species was used as the positive pair, and five negative host species were sampled from the benchmark candidate-host space while excluding the confirmed host genus where possible. This yielded one positive and five genus-balanced negative pairs per phage, or 3,804 phage–host pairs per evidence condition. The same pair set was used across all perturbation variants.

Each pair was evaluated with the pairwise prompt using the same Qwen3.6:27b backbone, phage profiles, host profiles and decoding settings as the host-list experiments. The pairwise prompt followed the input and output schema described above; prompt provenance and the limits of cross-configuration comparisons are documented in Supplementary Note S6. The model received one phage profile and one candidate host profile, and returned an infection-plausibility score in [0, 1] together with a categorical decision. Pairwise Hits@1 was computed per phage by comparing the confirmed-host score with the five negative-host scores.

To compare pairwise evaluation with host-list ranking under matched candidate density, we performed a fresh Listwise-6 inference experiment. For each phage, the host-list prompt was rebuilt to contain only the six hosts used in the pairwise evaluation: the experimentally confirmed host and the same five genus-balanced negative hosts. The query-phage profile, compact host metadata format, scoring guide, output schema, Qwen3.6:27b backbone and decoding settings were kept unchanged from the original Listwise-223 host-list experiment, except that the candidate catalogue and the required number of returned entries were reduced from 223 and 30 to 6 and 6, respectively.

Listwise-6 Hits@1 was computed directly from these fresh six-candidate host-list outputs. It was compared with pairwise Hits@1 on the same phage-level candidate sets. Argmax agreement measured whether the top-scoring host selected by fresh Listwise-6 ranking was identical to the top-scoring host selected by pairwise scoring among the same six hosts. A post-hoc restriction of the original Listwise-223 output to the same six hosts was used only as an auxiliary diagnostic and was not used for the main Listwise-6 result reported in Fig. 6 and Supplementary Table 10.

Discordance categories were defined by crossing full-list correctness with pairwise support for the confirmed host. A confirmed-host pairwise score ≥ 0.7 was treated as high-confidence positive support. Phages that were listwise incorrect but had high pairwise positive scores were interpreted as candidate-density displacement cases, whereas phages that were incorrect in both formulations and had low pairwise positive scores were treated as paradigm-independent hard cases. Genus-level divergence was computed by comparing full-list sp@1 and pairwise Hits@1 within each confirmed host genus. Numerical values are reported in Supplementary Table 10 and Supplementary Note S6.

### 4.7 Quantifying evidence-interface hallucination risk

#### Scope and definitions

We evaluated the frozen predictor (Qwen3.6:27b, no-thinking) in its full-evidence configuration on RefSeq-634, analysing for each phage the cached full-evidence inference output: the generated rationale, the ranked host output and the verbatim input profile. Because hallucination has no universal operational definition and prediction error alone is insufficient to identify it, we operationalised three related evidence-interface quantities, evaluating only the evidence available to the model rather than the external biological literature. *Profile faithfulness* is a rationale-level quantity: the fraction of rationale claims entailed by the input profile, with a claim considered unsupported when no profile field entails it. *Rationale-linked hallucination* is an incorrect species-level prediction whose rationale contains at least one claim not grounded in the supplied profile; this was operationalised as an unsupported field-referenced claim for the programmatic verifier, and as a contradicted or unverifiable semantic claim for the blind judge. *Answer-level hallucination* is the stricter case of an incorrect prediction whose predicted host genus appears nowhere in the host-supporting evidence fields of the supplied profile. Because all candidate hosts are visible in the closed-set ranking task, catalogue membership alone was not counted as supporting evidence. Prediction correctness was scored at the host-species level after whitespace/underscore normalisation, whereas answer-level hallucination and the error-source decomposition were evaluated at the host-genus level, the resolution at which the BLASTN-neighbour, 25-mer, CRISPR and RBP host associations are most consistently reported. Blind atomic-claim judging was applied to all 634 phages; resampling and reliability bench-marking used a shared correctness-balanced 200-phage sample (Balanced-200; 100 correct and 100 incorrect predictions drawn from the cached full-evidence outputs), and the held-out natural subset (*n* = 434) was its complement within RefSeq-634.

#### Rationale-grounding checks

The raw profile was parsed deterministically into a compact evidence digest containing per-gene annotations, BLASTN-neighbour hosts with identities and coverages, 25-mer and CRISPR-spacer host associations, explicit RBP matches, the Gram hint and the phage family; the same digest was supplied to the judges. Rationale faithfulness was assessed two ways. Following source-grounded claim verification, a deterministic verifier extracted six claim types—gene reference, gene function, RBP match, neighbour-host signal, Gram type and phage family—and checked each against the parsed profile under a strict literal setting and a predefined lenient setting that corrected two known verifier artefacts (RBP near-binding within a ±2% identity tolerance and a parent-lineage map) [45, 46]. Following LLM-as-judge and retrieval-augmented faithfulness protocols, a blind judge decomposed each trace into atomic claims, labelled each supported/contradicted/unverifiable against the digest, flagged decision-relevant claims and assigned a per-phage severity (none/minor/major) [47–50]. The blind semantic judge (Claude Opus 4.8, model identifier claude-opus-4-8, accessed via the Anthropic API) saw neither the true host nor the correctness label and was applied to all 634 phages; rationale-linked semantic hallucination was counted for an incorrect prediction with at least one decision-relevant claim labelled contradicted or unverifiable. The digest, verifier rules, judge prompt and output schema are given in Supplementary Note S8.1–S8.4.

#### Answer stability and inference-time reliability

To measure answer stability we resampled the predictor on the balanced 200-phage sample (*N* = 10, *T* = 0.7, identical saved prompt); answer-level hallucinations that fell outside this sample were additionally resampled so that this subset had complete coverage. Resampled top-1 predictions were clustered by host genus, from which we computed genus/species self-consistency, modal consistency, semantic entropy and the number of distinct genera. To benchmark auxiliary reliability signals, we computed AUROC for predicting top-1 correctness from the top-1–top-2 score margin, the top-1 score, homology-derived features, resampling self-consistency and semantic entropy. The score margin was interpreted as ranking-level decisiveness and self-consistency as whether a host call was stably reproducible from the supplied evidence. For auxiliary selective-prediction analysis, score-margin thresholds were fixed on the balanced validation split and applied unchanged to the held-out natural subset (*n* = 434), without test-set tuning. Details are given in Supplementary Note S8.5–S8.6.

#### Error-source decomposition and statistics

Each wrong RefSeq-634 prediction was characterised by two facts from the parsed host-supporting evidence fields, defined as the union of host genera named by the BLASTN-neighbour, 25-mer, CRISPR and RBP evidence blocks: whether the true host genus was recoverable, and whether the predicted genus was itself an evidence candidate. A prediction was *off-evidence* when its predicted genus was absent from these host-supporting fields, a flag computable without the ground-truth host. An off-evidence prediction was counted as an answer-level hallucination only when it was also wrong at the species level. For mechanism analysis, errors were partitioned in priority order into *incomplete evidence* (true genus absent), *non-decisive evidence* (true and predicted genera both present, a competitor ranked first) and *recoverable-profile hallucination* (true genus present but predicted genus absent). Evidence-incomplete hallucinations, in which both the true and predicted genera were absent, fall within the incomplete-evidence class; when answer-level hallucinations were reported separately, the incomplete-evidence class was split into evidence-incomplete answer-level hallucinations and evidence-anchored but wrong incomplete-evidence cases. For each class we summarised the score margin and resampling self-consistency, reporting bucket-level AUROC for the larger classes and descriptive statistics for the small hallucination subtypes, whose sample sizes make bucket-level AUROC unstable. All headline effect sizes were reported with bootstrap 95% confidence intervals; categorical enrichments used Fisher’s exact test with odds ratios, correlations used point-biserial *r*, self-consistency and entropy differences additionally used permutation tests, and AUROC values were bootstrapped over phages (Supplementary Note S8.7–S8.8).

### 4.8 Layerwise emergence analysis (target-conditioned local Jacobian readout)

To characterize how intermediate decoder activations contribute, to first order, to the confirmed host-genus logit, we applied a per-input, target-conditioned local Jacobian attribution to each RefSeq-634 phage under the full profile. The model received the fixed evidence profile followed by the answer prefix, and the readout was evaluated at the single query position immediately preceding host-name generation (the token after `“host”`: ”). The confirmed host genus was not inserted as a ground-truth answer in the output prefix and served only as the differentiation target, although its lexical form was present in the candidate-host catalogue and could additionally occur in host-linking evidence fields. Adapting the Jacobian-lens idea [14] to a single input, we read the residual stream through the first-order sensitivity of the confirmed host-genus logit to each decoder-block activation, *r_ℓ_* = ⟨*∂z*_host_*/∂h_ℓ_, h_ℓ_*⟩, computed in reverse mode with model parameters frozen (strided every four of the 64 text-decoder blocks). Qwen3.6:27b is a multimodal conditional-generation model; activations were read from the text-decoder submodule only (the vision tower was not exercised); three of every four decoder blocks use linear attention and the remaining block uses full attention. This is a first-order (input×gradient) attribution of one target logit at one position; unlike the original context-averaged Jacobian lens it involves no averaging over contexts, produces no vocabulary ranking, and uses the label token to define the read direction, so it is a target-conditioned attribution rather than an independent decoder or a causal readout. Activations were the post-block residual-stream states and gradients were obtained by reverse-mode automatic differentiation on a matched bf16 checkpoint, because gradients are not available through the quantised inference stack used for the deployed predictions; the readout therefore characterises a closely matched full-precision model rather than the exact quantised inference-time weights, and correct and incorrect groups were defined by the deployed quantised model’s species-level top-1 predictions (427 and 207 phages). We defined the emergence layer as the first sampled block at which the positive, peak-normalised *r_ℓ_* reached 20% of that phage’s positive peak, verified that this crossing was stable for thresholds from 10% to 50%, and compared the median signed *r_ℓ_* trajectory between correct and incorrect predictions; magnitude differences (raw positive-attribution peak and mean over the mid-to-late relative-depth window, layers 32–53 of the 64-block Qwen3.6:27b model) were assessed with phage-level bootstrap 95% confidence intervals. All 634 phages were eligible (single-token genus target); one phage’s Jacobian curve did not reach the emergence threshold at any sampled layer and was excluded from emergence-layer statistics only. Full definitions, the threshold-sensitivity and magnitude results, and the final-block behaviour are given in Supplementary Note S11.

### 4.9 Prospective post-cutoff phage case study

As a temporal evidence-attribution control, we applied the unchanged RefSeq-634 full-profile interface to 14 GenBank bacteriophages whose confirmed host species fell within the 223-species RefSeq-634 candidate catalogue. Profiles were built with the identical deterministic pipeline (genome annotation, RBP homology, BLASTN nucleotide-neighbour, alignment-free 25-mer and CRISPR-spacer blocks), query identifiers were masked, and predictions used the same frozen Qwen3.6:27b backbone, prompt, decoding settings and closed candidate catalogue as the primary experiments, together with the parametric-only and no-RBP ablation variants. Full provenance, deposition dates, leakage flags, per-genome evidence-ablation results and the illustrative vignettes are provided in Supplementary Note S12, Supplementary Table 27 and Fig. 4c.

### 4.10 Eukaryotic virus–host portability case study

As a supplementary portability probe, we applied the unchanged PHI-Reason profile–prompt interface to the EvoMIL 36-host eukaryotic virus–host benchmark, using a shared 3,186-virus set for the gpt-oss-120B, Qwen3-Coder-Next and baseline comparisons, and reporting Qwen3.6:27b separately on the full 3,198-virus set. Within each reported evaluation set, abstentions, unparsable outputs and out-of-catalogue answers were counted as errors. The Qwen3.6:27b result was not a strictly matched-denominator comparison (Supplementary Table 14). The case study used fold-aware five-fold cross-validation, query and homolog-neighbour identifier masking, and a eukaryotic evidence schema substituted for the phage-specific receptor-binding-protein, 25-mer and CRISPR modules. Full protocol details, leakage controls, baselines, metrics and homolog-neighbour audits are provided in Supplementary Note S10, Supplementary Tables 12–15 and Supplementary Figure 7 (per-virus values in Source Data 3).

### 4.11 Typed-decision inference-interface transfer (Jev)

As an architecture-level portability probe, we applied the unmodified RefSeq-634, VHDB-3150 and HiC full-evidence profiles to Jev (TypeSafe AI) [28, 29], a commercial typed-decision inference system that returns structured probabilistic judgments over an explicit candidate set rather than free-form generated text. Candidate hosts were supplied through Jev’s choice primitive, with no change to the profile content, evidence schema or candidate catalogue used for the language-model backbones. Query construction, VHDB-3150’s candidate-batching procedure and error handling are detailed in Supplementary Table 28.

### 4.12 Statistical analysis

All statistical analyses were performed on fixed model outputs without model retraining or parameter updating. Unless otherwise stated, denominators were kept fixed under the abstention-as-error policy. Confidence intervals were matched to the estimand. Single-binomial proportions and accuracies with one binary outcome per item, including sp@1, genus-level accuracy, pairwise Hits@1, any-hit top-1, strict top-1, prediction coverage and off-evidence or hallucination rates, were reported with two-sided Wilson score 95% confidence intervals. Absolute differences between independent subgroups, including Gram-stratified, BLASTN-neighbour-availability and shared-failure evidence-coverage contrasts where the compared phages were disjoint, were reported with Newcombe hybrid-score 95% confidence intervals constructed from the two component Wilson intervals. Rank-sensitive or continuous metrics, including MRR, MHA-sp, MHA-gen, top-1–top-2 score margins, self-consistency, AUROC, PR-AUC and log-loss, were reported with non-parametric percentile bootstrap 95% confidence intervals. Host-list bootstrap intervals used 1,000 phage-level resamples with random seed 42; pairwise AUROC, PR-AUC and log-loss intervals used phage-cluster bootstrap resampling that retained all evaluated pairs for each resampled phage rather than independent pair-level resampling.

Between-method comparisons on shared phage sets were assessed using paired statistical tests. Paired per-phage success differences were tested with the two-sided Wilcoxon signed-rank test, and binary top-1 discordance tables were additionally evaluated with McNemar’s [51] test with Yates’ correction. Categorical enrichment analyses used Fisher’s exact test; related tests were adjusted using the Benjamini– Hochberg false-discovery-rate procedure. Shared-failure overlap was assessed with a two-sided Fisher’s exact test and a 10,000-resample phage-level overlap permutation null. The three shared-failure evidence-coverage comparisons (BLASTN, CRISPR and RBP) are reported as unadjusted two-sided Fisher *p* values.

For pairwise evaluation, F1 and MCC were reported at the threshold that maximised F1 on the pooled evaluated pair set; this threshold was used only for post hoc score evaluation and not for model selection. Log-loss was computed after clipping predicted probabilities to [10*^−^*^7^, 1 − 10*^−^*^7^]. Correlations between host-list and pair-wise scores were computed using Spearman rank correlation, Pearson correlation and point-biserial correlation where appropriate. The relationship between reasoning-trace breadth and correctness was assessed using the Cochran–Armitage trend test and logistic regression. Analyses were implemented in Python using numpy [52], scipy [53], scikit-learn [54] and statsmodels [55]; detailed scripts and package requirements are provided with the released code.

### 4.13 Performance metrics

#### Top-k species accuracy (sp@k)

For single-host benchmarks, each test phage *q* has one ground-truth host species *y_q_*, and each method returns a ranked list *π*^*_q_* over the benchmark-specific candidate host space:

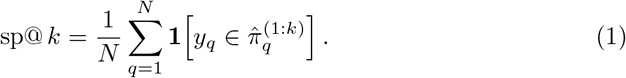

For PHI-Reason listwise inference, the model returned the top 30 ranked hosts; candidates absent from this output were treated as unranked. sp@1 was the primary metric on RefSeq-634 and VHDB-3150, with sp@5 and sp@10 reported as broader ranking diagnostics. Genus-level accuracy (g@*k*) was computed analogously after mapping predicted and true hosts to genus.

#### Mean reciprocal rank (MRR)

MRR was used as a rank-sensitive complement to top-*k* accuracy:

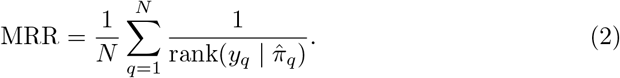

If the true host was outside the candidate space or absent from the ranked output, its contribution was set to zero.

#### Multi-host accuracy (MHA-sp and MHA-gen)

For the HiC benchmark, each phage can have multiple confirmed host species *Y_q_*. We followed the MHA protocol of [16], comparing the first |*Y_q_*| predictions with the known host set:

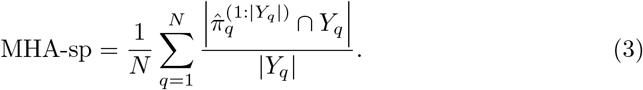

MHA-sp reports the mean fraction of confirmed host species recovered when each method returns the same number of predictions as known hosts for that phage. MHA-gen was computed after mapping host labels to genus. MHA-sp was the primary HiC metric, with MHA-gen, single-top species accuracy and MRR reported as secondary diagnostics where ranked outputs were available.

#### Pairwise evaluation metrics

For pairwise PHI assessment, each phage was paired with one confirmed host and five genus-balanced negative hosts. Each pair received an infection-plausibility score *s_i_* ∈ [0, 1]. Because this task is class-imbalanced and intended for prioritisation, PR-AUC was the primary score-based metric and ROC-AUC was retained as a secondary rank-discrimination metric. F1 and MCC were reported at the threshold that maximised F1. Log-loss was used as a calibration-sensitive metric:

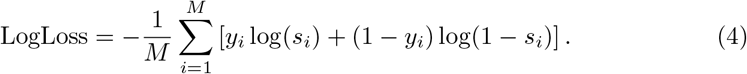

High-confidence precision was computed among pairs with *s_i_* ≥ 0.7:

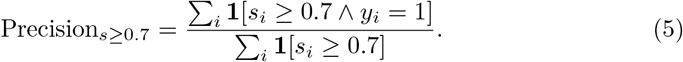

For each phage, pairwise Hits@1 compared the confirmed-host score with the five negative-host scores. Unless otherwise stated, we used a lenient definition in which ties were counted as correct:

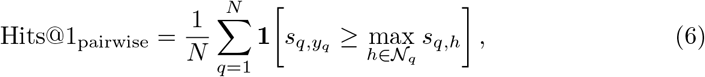

where N*_q_* denotes the five negative hosts. Strict Hits@1, in which ties were counted as misses, was used only as a diagnostic. Positive and negative score separation was summarized by Cohen’s *d*:

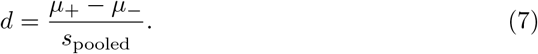

#### Host-list versus pairwise comparison metrics

Candidate-density effects were assessed using a fresh Listwise-6 control. For each phage, the host-list prompt was rebuilt to contain the same six hosts used in pairwise evaluation: the confirmed host and five genus-balanced negatives. The phage profile, host metadata format, scoring guide, output schema, model backbone and decoding settings were unchanged from the full Listwise-223 experiment, except that the candidate catalogue and required output length were reduced from 223 and 30 to 6 and 6.

Listwise-6 Hits@1 was compared with pairwise Hits@1 on the same six-host candidate sets. The candidate-density component was defined as

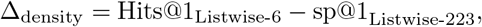

and the residual formulation difference as

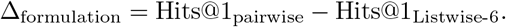

Argmax agreement measured the fraction of phages for which pairwise scoring and fresh Listwise-6 selected the same top-scoring host among the six candidates.

## Data availability

Input datasets used in this study are publicly available. The CHERRY-1940 corpus from which the RefSeq-634 evaluation partition was drawn is accessible through the NCBI RefSeq database through the original release at github.com/KennthShang/CHERRY. The VHDB-3150 benchmark and its associated phage–host pair table are provided by the VHDB authors at github.com/KennthShang/HostPredictionReview. The HiC proximity-ligation dataset, including the 406 viral contigs and 52 metagenome-assembled host genomes, is available from the source study zenodo.org/records/14851637. The PHIStruct receptor-binding-protein corpus used as the RBP reference database (19,081 sequences across 238 host genera) is available from zenodo.org/records/11202338. Functional-annotation reference databases were obtained from PHROG v4 (phrogs.lmge.uca.fr) and eggNOG 5 via eggnog-mapper v2. Defence-system reference profiles were obtained from DefenseFinder v2.0.1 (github.com/mdmparis/defense-finder). The 14 recently deposited phage genomes analysed in the prospective case study (ten cleanly post-cutoff and four temporally flagged; Supplementary Note S12, Supplementary Table 27) were retrieved from NCBI GenBank under accessions PX563682, PV491271, PQ473534, PV872382, PZ236371, PZ319873, PV815418, PX609837, PP895303, PX055572, PX093639, PQ074102, PX138265 and PZ133968; their per-genome host assignments, deposition and release dates, and reference-neighbour status are listed in Supplementary Table 27.

## Ethics approval and consent to participate

Not applicable.

## Consent for publication

Not applicable.

## Code availability

Code required to reproduce this study, including the structured-profile construction pipeline and the host-list and pairwise inference drivers, is publicly available at github.com/yaozhong/PHI-Reason.

## Authors’ contributions

Y.Z. conceived and designed the study. L.X. and Y.Z. developed the computational methods, conducted the experiments, and performed the data analysis. L.X., Y.Z. and S.I. wrote the manuscript. Y.Z. and S.I. supervised the project.

## Acknowledgments

Computing resources were provided by the Human Genome Center, Institute of Medical Science, University of Tokyo. This work was supported by Grant-in-Aid for Scientific Research [JSPS KAKENHI grant number JP26K15038]. L.X. acknowledges the financial support from the China Scholarship Council (CSC) under grant No. 202506120217.

Generative AI tools were used as components of the research for structured-evidence inference and rationale evaluation. The tools, model versions, prompts, and inference settings are documented in the Methods and Supplementary Materials. AI-assisted tools were also used for language polishing. All text was reviewed and verified by the authors.

