## Supplementary material for "Structured biological evidence enables general-purpose language models to predict bacteriophage–host interactions": Supplmentary

### Supplementary Tables

*Organisation. Supplementary Tables 1–32 support the main-text claims (cross-benchmark performance, evidence ablation, host strata, hallucination and reliability audit, eukaryotic transfer, prompts, construction parameters, dataset provenance, compute and the prospective post-cutoff case study; Tables 31–32 cover the external 207-pair wet-lab-supported validation). The cross-benchmark accuracy, per-phage concordance and per-host-family advantage are visualised in Supplementary Figure 3. Per-item, per-host and per-virus audits are additionally supplied as machine-readable Source Data 1–3 (Data Availability). All accuracies are argmax top-1 with a full-set denominator (abstention counted as error) unless noted.*

**Supplementary Table 1.** Cross-benchmark performance of all evaluated methods (species- and genus-level accuracy with 95% Wilson CIs; prediction coverage).

| | RefSeq-634 | | | | VHDB-3150 | | Coverage ($n$) | |
|---|---|---|---|---|---|---|---|---|
| Method | sp@1 | sp@1 CI | g@1 | g@1 CI | sp@1 | g@1 | RefSeq | VHDB |
| PHI-Reason | 67.4 | 63.6–70.9 | 80.9 | 77.7–83.8 | 57.2 | 76.8 | 634 | 3150 |
| PhaBOX2 (MAG, full DB) | 49.4 | 45.5–53.3 | 61.5 | 57.7–65.2 | 43.8 | 55.5 | 631 | 2981 |
| PHP (retrain) | 46.1 | 42.2–49.9 | 56.9 | 53.1–60.7 | 32.8 | 52.5 | 634 | 3150 |
| PHIST | 23.3 | 20.2–26.8 | 34.4 | 30.8–38.2 | 16.5 | 29.6 | 534 | 2649 |
| WIsH | 19.7 | 16.8–23.0 | 31.2 | 27.7–34.9 | 13.5 | 26.7 | 634 | 2982 |
| Embedding-only control | 56.6 | 52.7–60.4 | 69.2 | 65.5–72.6 | n/r | n/r | 634 | n/r |

**Supplementary Table 2.** Structured-profile evidence ablation on RefSeq-634: additive ladder, single-field additions and leave-one-field removals, with genus-level accuracy and 95% CIs.

| Configuration | sp@1 | sp@1 CI | g@1 | g@1 CI | Δ (95% CI) |
|---|---|---|---|---|---|
| Base (annotation only) | 22.6 | 19.5–26.0 | 38.2 | 34.5–42.0 | 0 (reference) |
| Base + RBP | 41.0 | 37.2–44.9 | 63.4 | 59.6–67.1 | +18.5 (+13.4, +23.4) |
| Base + RBP + BLASTN | 64.0 | 60.2–67.7 | 79.8 | 76.5–82.8 | +41.5 (+36.4, +46.3) |
| + 25-mer (=Full−CRISPR) | 66.4 | 62.6–70.0 | 80.0 | 76.7–82.9 | +43.8 (+38.8, +48.6) |
| Full (all four) | 67.4 | 63.6–70.9 | 80.9 | 77.7–83.8 | +44.8 (+39.7, +49.5) |
| *Single field added to Base (Δ vs Base):* | | | | | |
| Base + BLASTN | 56.9 | 53.1–60.7 | 69.7 | 66.0–73.2 | +34.4 (+29.2, +39.3) |
| Base + RBP | 41.0 | 37.2–44.9 | 63.4 | 59.6–67.1 | +18.5 (+13.4, +23.4) |
| Base + CRISPR | 36.9 | 33.2–40.7 | 50.3 | 46.4–54.2 | +14.3 (+9.3, +19.3) |
| Base + 25-mer | 29.7 | 26.2–33.3 | 44.3 | 40.5–48.2 | +7.1 (+2.3, +11.9) |
| *Leave-one field from Full (Δ vs Full):* | | | | | |
| Full − BLASTN | 49.2 | 45.3–53.1 | 68.8 | 65.1–72.3 | −18.1 (−23.4, −12.7) |
| Full − RBP | 63.1 | 59.3–66.8 | 74.8 | 71.2–78.0 | −4.3 (−9.5, +1.0) |
| Full − CRISPR | 66.4 | 62.6–70.0 | 80.0 | 76.7–82.9 | −0.9 (−6.1, +4.2) |
| Full − 25-mer | 68.0 | 64.3–71.5 | 81.5 | 78.3–84.4 | +0.6 (−4.5, +5.8) |

**Supplementary Table 3.** Evidence-field coverage and specificity across RefSeq-634.

| Evidence field | coverage $n/634$ | coverage % | genus hit-rate when present |
|---|---|---|---|
| 25-mer shared count | 534 | 84.2 | top-5 contains true genus 55.8% (298/534) |
| RBP homology (BLASTP) | 516 | 81.4 | top-5 contains true genus 72.7% (375/516) |
| BLASTN genomic neighbour | 469 | 74.0 | neighbour host = true genus 88.7% (416/469) |
| CRISPR spacer link | 184 | 29.0 | nominated set contains true genus 86.4% (159/184) |

**Supplementary Table 4.** Backbone–mode generalization on RefSeq-634 (same full evidence profiles, prompt, catalogue and decoding; only the frozen backbone and thinking mode change). Qwen3.6-27B is the deployed backbone for all other analyses in this paper; among the seven backbone–mode configurations tested here it attained the highest observed species-level top-1 accuracy (67.4% without thinking, 67.7% with thinking), exceeding Qwen3-Coder-Next while generating fewer tokens in its no-thinking configuration despite having the fewest parameters (27.8B dense) of the three higher-capacity backbones evaluated. Enabling thinking on Qwen3.6-27B did not improve species-level accuracy (67.7% versus 67.4%, overlapping confidence intervals) while generating 5.3× more tokens (6,378 versus 1,199), consistent with the other backbones in showing no reliable accuracy gain from extended reasoning traces.

| Backbone × mode | sp@1 | sp@1 CI | sp@5 | g@1 | MRR | mean gen. tokens |
|---|---|---|---|---|---|---|
| Qwen3-4B, thinking | 39.0 | 35.2–42.8 | 61.4 | 53.9 | 0.503 | 3497 |
| Qwen3-4B, no-thinking | 48.0 | 44.1–51.8 | 73.0 | 64.4 | 0.589 | 1012 |
| GPT-oss-120B, thinking | 65.8 | 62.0–69.4 | 86.4 | 79.8 | 0.745 | 7731 |
| GPT-oss-120B, no-thinking | 63.3 | 59.4–66.9 | 83.6 | 79.2 | 0.717 | 1624 |
| Qwen3.6-27B, no-thinking (deployed) | 67.4 | 63.6–70.9 | 87.9 | 80.9 | 0.760 | 1199 |
| Qwen3.6-27B, thinking | 67.7 | 63.9–71.2 | 88.0 | 80.3 | 0.758 | 6378 |
| Qwen3-Coder-Next 79B | 63.6 | 59.7–67.2 | 86.0 | 78.4 | 0.732 | 1296 |

**Supplementary Table 5.** Receptor-binding-protein reference corpus and filtering.

| Item | Value |
|---|---|
| Reference corpus | PHIStruct RBP database |
| Sequences / host genera | 19,081 RBP proteins across 238 host genera |
| Aligner | DIAMOND v2.1.24 (`blastp`) |
| Filters | identity $\geq$ 40%, query coverage $\geq$ 5%, e-value $\leq 10^{-5}$ |
| Aggregation | top-5 host genera per query RBP, ranked by pident$^{0.6}$qcov$^{0.4}$ |
| De-leaking | reference entries at $\geq$ 95% identity and $\geq$ 85% coverage to evaluation proteins removed |

**Supplementary Table 6.** Reference-label perturbation controls on RefSeq-634 (host-label withholding and per-block scrambling).

| Condition | Perturbed block | sp@1 | sp@1 CI | g@1 | Δ sp@1 vs Full (95% CI) |
|---|---|---|---|---|---|
| Full | none | 67.4 | 63.6–70.9 | 80.9 | 0 |
| Label-free (labels withheld) | BLASTN/25-mer/CRISPR | 41.3 | 37.6–45.2 | 63.4 | −26.0 (−31.2, −20.6) |
| BLASTN scrambled | BLASTN | 44.0 | 40.2–47.9 | 63.9 | −23.3 (−28.5, −17.9) |
| 25-mer scrambled | 25-mer | 68.6 | 64.9–72.1 | 81.5 | +1.3 (−3.9, +6.4) |
| CRISPR scrambled | CRISPR | 65.6 | 61.8–69.2 | 79.8 | −1.7 (−6.9, +3.5) |
| All labels scrambled | all three | 33.1 | 29.6–36.9 | 52.8 | −34.2 (−39.2, −28.9) |

**Supplementary Table 7.** Gram-type and reference-coverage stratification of the full profile (Supplementary Note S1).

*Panel a — taxonomic strata*

| Partition | $n$ | sp@1 | sp@1 CI | g@1 | g@1 CI |
|---|---|---|---|---|---|
| Gram-positive | 353 | 71.4 | 66.5–75.9 | 90.4 | 86.8–93.0 |
| Gram-negative | 270 | 63.3 | 57.4–68.9 | 68.1 | 62.4–73.4 |
| Archaea | 11 | 36.4 | 15.2–64.6 | 90.9 | 62.3–98.4 |
| All | 634 | 67.4 | 63.6–70.9 | 80.9 | 77.7–83.8 |

*Panel b — per-field Gram contrast (Base→Base+field, sp@1; ablation-suite profile build, see Supplementary Table 2 footnote)*

| Field added | Gram+ sp@1 | Gram− sp@1 | Gram+ − Gram− (pp) |
|---|---|---|---|
| base | 22.1 | 24.1 | −2.0 |
| + RBP | 31.4 | 54.4 | −23.0 |
| + BLASTN | 61.2 | 50.4 | +10.8 |
| + 25-mer | 34.0 | 24.8 | +9.2 |
| + CRISPR | 40.8 | 32.2 | +8.6 |

*Panel c — reference-coverage strata*

| Group | $n$ | sp@1 | sp@1 CI | g@1 | g@1 CI |
|---|---|---|---|---|---|
| Has BLASTN neighbour | 469 | 72.7 | 68.5–76.5 | 84.2 | 80.6–87.2 |
| No BLASTN neighbour | 165 | 52.1 | 44.5–59.6 | 71.5 | 64.2–77.9 |

**Supplementary Table 8.** Shared-failure analysis across backbones and baselines (CBI/BMD).

*Panel a — enrichment*

| Denominator | CBI, BMD, overlap | Odds ratio | 95% CI | Fisher $p$ |
|---|---|---|---|---|
| All 634 phages | 149, 196, 89 | 5.24 | 3.54–7.75 | $4.6 \times 10^{-17}$ |
| 532 valid-prediction subset | 127, 169, 78 | 5.49 | 3.58–8.42 | $2.3 \times 10^{-15}$ |

*Panel b — host-linking evidence coverage (shared-failure $n = 89$ vs remainder $n = 545$)*

| Channel | Shared-failure (%) | Remainder (%) | Difference (95% CI) | Fisher $p$ |
|---|---|---|---|---|
| BLASTN neighbour present | 58.4 | 76.5 | −18.1 (−29.0, −7.7) | $6 \times 10^{-4}$ |
| CRISPR-spacer link | 13.5 | 31.6 | −18.1 (−25.0, −8.7) | $4 \times 10^{-4}$ |
| RBP support | 84.3 | 80.9 | +3.4 (−6.1, +10.4) | 0.56 |

**Supplementary Table 9.** Pairwise discrimination across evidence configurations (634 positive and 3,170 genus-balanced negative pairs).

| Configuration | pos | neg | PR-AUC (95% CI) | ROC-AUC (95% CI) | Cohen $d$ | prec.@0.7 | Hits@1 (len./strict) | tie % |
|---|---|---|---|---|---|---|---|---|
| Full profile | 634 | 3170 | 0.962 | 0.992 | 6.10 | 0.937 | 0.972 / — | 3.2 |
| Base-only | 634 | 3170 | 0.748 | 0.949 | 2.70 | 0.735 | 0.860 / — | 12.2 |

**Supplementary Table 10.** Host-list versus pairwise candidate-density control.

| Setting | candidates | metric | value (95% CI) | McNemar $p$ vs pairwise |
|---|---|---|---|---|
| Listwise | 223 | sp@1 | 0.674 (0.636–0.709) | — |
| Listwise | 6 | sp@1 | 0.957 (0.939–0.971) | — |
| Pairwise | 6 | Hits@1 (lenient) | 0.972 (0.956–0.982) | — |

**Supplementary Table 11.** Per-host-family species accuracy on RefSeq-634 with 95% Wilson CIs and paired margin over the strongest baseline.

| Family | $n$ | PHI sp@1 | PHI CI | best baseline | baseline CI | Δ (95% CI) |
|---|---|---|---|---|---|---|
| Enterobacteriaceae | 135 | 56.3 | 47.9–64.4 | 22.2 | 16.0–29.9 | +34.1 (+22.6, +44.3) |
| Staphylococcaceae | 15 | 80.0 | 54.8–93.0 | 53.3 | 30.1–75.2 | +26.7 (−6.7, +53.3) |
| Streptococcaceae | 34 | 76.5 | 60.0–87.6 | 64.7 | 47.9–78.5 | +11.8 (−9.7, +31.9) |
| Mycobacteriaceae | 98 | 100.0 | 96.2–100.0 | 92.9 | 86.0–96.5 | +7.1 (+1.9, +14.0) |
| Bacillaceae | 42 | 45.2 | 31.2–60.1 | 50.0 | 35.5–64.5 | −4.8 (−24.9, +15.9) |
| Gordoniaceae | 60 | 25.0 | 15.8–37.2 | 55.0 | 42.5–66.9 | −30.0 (−45.1, −12.5) |

**Supplementary Table 12.** Eukaryotic virus–host benchmark protocol (EvoMIL 36-host closed set).

| Item | Setting |
|---|---|
| Candidate catalogue | 36 eukaryotic host species (closed set) |
| Viruses (full / shared) | 3,198 / 3,186 |
| Cross-validation | fold-aware five-fold; same species confined to one fold |
| Retrieval exclusions | query, same-species, BLASTN $\geq$ 95% neighbours |
| Identifier masking | query and homolog-neighbour IDs hashed |
| Error policy | abstentions / unparsable / out-of-catalogue = errors |
| Primary metric | any-hit top-1 (secondary: strict top-1, macro-F1) |

**Supplementary Table 13.** Eukaryotic versus phage profile schema.

| Block | Phage profile | Eukaryotic profile |
|---|---|---|
| Base genome architecture | yes | yes |
| Gene/protein functions | yes | yes |
| Taxonomy | yes | yes (added emphasis) |
| Baltimore class / segmentation | — | added |
| BLASTP homolog-neighbour context | RBP matches only | added (whole-proteome) |
| RBP / 25-mer / CRISPR | yes | removed |
| Floor profile P0 | — | architecture and gene function only |

**Supplementary Table 14.** Eukaryotic case-study results and baselines (any-hit and strict top-1 with 95% Wilson CIs; macro-F1; floor-profile P0 any-hit).

*Panel a — full-catalogue comparison (36 hosts)*

| Method / configuration | $N$ | any-hit | any-hit CI | strict | macro-F1 | P0 any-hit |
|---|---|---|---|---|---|---|
| PHI-Reason (Qwen3.6:27b, full)$^{\dagger}$ | 3198 | 67.0 | 65.3–68.6 | 61.9 | 0.401 | 19.2 |
| PHI-Reason (gpt-oss:120b, full) | 3186 | 67.0 | 65.3–68.6 | 61.7 | 0.396 | 35.1 |
| PHI-Reason (Qwen3-Coder-Next, full) | 3186 | 66.6 | 64.9–68.2 | 61.5 | 0.394 | 23.7 |
| kNN homology retrieval | 3186 | 63.4 | 61.7–65.1 | 57.6 | 0.312 | — |
| EvoMIL (ESM-1b + MIL) | 3186 | 60.4 | 58.7–62.1 | 56.9 | 0.359 | — |
| Composition $k$-mer RF | 3186 | 58.8 | 57.1–60.5 | 54.1 | 0.282 | — |
| Nucleotide LM + MIL | 3186 | 57.1 | 55.4–58.8 | 53.5 | 0.312 | — |

*Panel b — decision-rule sensitivity on the homolog-available subset ($n = 3162$)*

| Rule / method | any-hit | any-hit CI | note |
|---|---|---|---|
| PHI-Reason (Qwen3-Coder-Next) | 67.0 | 65.3–68.6 | integrated evidence |
| kNN homology retrieval | 63.8 | 62.2–65.5 | $k$-neighbour vote |
| Majority-of-homologs rule | 53.1 | 51.4–54.9 | plurality host among hits |

**Supplementary Table 15.** BLASTN nucleotide-identity stratification (eukaryotic; any-hit top-1, PHI-Reason Qwen3-Coder-Next vs kNN homology). Strata are defined by whole-genome BLASTN identity to the reference set and cover the full eukaryotic set ($n = 3{,}198$). The no-BLASTN-neighbour stratum ($n = 236$) is a nucleotide-similarity category distinct from the BLASTP protein-homolog-available subset ($n = 3{,}162$) used in Supplementary Table 14b; because BLASTP is more sensitive than BLASTN, more viruses have a protein homolog than a nucleotide neighbour. The 3,186 shared denominator in Supplementary Table 14a applies only to the cross-method comparison.

| BLASTN identity stratum | $n$ | PHI-Reason | kNN | $\Delta$ |
|---|---|---|---|---|
| Strong ($\geq$ 70% BLASTN identity) | 2937 | 66.7 | 63.2 | +3.5 |
| Weak (< 70% BLASTN identity) | 25 | 24.0 | 20.0 | +4.0 |
| No BLASTN neighbour | 236 | 67.4 | 66.5 | +0.9 |
| All (full set) | 3198 | 66.4 | 63.1 | +3.3 |

**Supplementary Table 16.** Programmatic hallucination verifier: claim-level support by claim type (strict field-specific and lenient entity-level).

| Claim type | $n$ claims | strict support (%) | lenient support (%) |
|---|---|---|---|
| Gene reference | 1458 | 100 | 100 |
| RBP match | 927 | 94 | 94 |
| 25-mer host | 619 | 97 | 97 |
| CRISPR host | 119 | 94 | 94 |
| BLASTN-neighbour host | 173 | 98 | 98 |
| Gene-function attribution | 69 | 49 | 49 |
| Aggregate | 3,365 | 96.4 | 96.4 |

**Supplementary Table 17.** Genus-level answer-anchor and error partition (207 errors of 634 predictions).

| Partition | $n$ | % of 634 | % of 207 errors |
|---|---|---|---|
| Correct (anchored) | 427 | 67.4 | — |
| Non-decisive error | 157 | 24.8 | 75.8 |
| Evidence-incomplete, anchored-incorrect | 31 | 4.9 | 15.0 |
| Evidence-incomplete, off-evidence | 17 | 2.7 | 8.2 |
| Recoverable-profile off-evidence | 2 | 0.3 | 1.0 |
| All off-evidence | 19 | 3.0 | 9.2 |

**Supplementary Table 18.** Reliability signals and selective prediction.

| Signal | group | mean | AUROC (95% CI) |
|---|---|---|---|
| Top-1–top-2 margin | correct | 0.146 | 0.854 (0.824–0.883) |
| Top-1–top-2 margin | incorrect | 0.036 | — |
| Top-1–top-2 margin | answer-level hallucination ($n = 19$) | 0.030 | — |
| Top-1–top-2 margin | Balanced-200 | — | 0.823 (0.761–0.877) |
| Self-consistency (genus) | correct | 0.981 | 0.672 |
| Self-consistency (genus) | wrong | 0.835 | — |
| Self-consistency (genus) | answer-level hallucination ($n = 19$, off-evidence subtypes pooled) | 0.589 | — |

**Supplementary Table 19.** Hi-C multi-host accuracy (Sp-1, Sp-2; PhaBOX2 excluded).

| Method | Sp-1 MHA-sp | Sp-1 MHA-g | Sp-2 MHA-sp | Sp-2 MHA-g |
|---|---|---|---|---|
| PHI-Reason | 0.517 | 0.893 | 0.673 | 0.976 |
| PHP (retrain) | 0.425 | 0.911 | 0.510 | 0.847 |
| WIsH | 0.342 | 0.857 | 0.480 | 0.777 |
| PHIST | 0.283 | 0.768 | 0.490 | 0.812 |
| DeepHost (retrain) | 0.383 | 0.661 | 0.357 | 0.682 |

**Supplementary Table 20.** Per-phage concordance with baselines (method-unique correct predictions).

| Comparison (RefSeq-634) | PHI-only | baseline-only | both |
|---|---|---|---|
| vs PhaBOX2 | 173 | 59 | 254 |
| vs PHP | 174 | 39 | 253 |
| vs PHIST | 287 | 8 | 140 |
| vs WIsH | 329 | 27 | 98 |

**Supplementary Table 21.** Construction parameters for structured profiles.

| Component | Tool / database | identity | coverage | e-value | $k$ / mismatch | top-$k$ |
|---|---|---|---|---|---|---|
| RBP match | DIAMOND v2.1.24 / PHIStruct | $\geq$ 40% | $\geq$ 5% | $\leq 10^{-5}$ | — | top-5 genera |
| BLASTN neighbour | NCBI BLAST+ v2.16.0 | $\geq$ 70% | $\geq$ 3% | — | — | top-5 (bitscore) |
| 25-mer | canonical $k$-mer index | — | — | — | $k = 25$ | shared-count rank |
| CRISPR spacer | spacer–genome match | $\geq$ 90% | $\geq$ 90% len | — | $\leq$ 2 mismatch | — |
| Eukaryotic BLASTP | DIAMOND v2.1.24 | — | — | $\leq 10^{-5}$ | — | top-50 |

**Supplementary Table 22.** Schematic prompt, output schema, decoding and parsing (host-list task, primary backbone); the complete unabridged prompt and parser are in the code repository (Code availability).

System prompt

```
/no_think
You are a bacteriophage-host infection prediction system.
```

User message (structure)

```
INSTRUCTIONS:
  Below is a phage genome profile, followed by a list of candidate host species.
  Select the 30 hosts most likely to be infected by this phage and assign each an
  infection probability score (0.0-1.0).  ...  Consider all available evidence:  tail/
  receptor-binding proteins and their BLASTP host matches, lysis-cassette genes, genomic
  neighbor context, genome-wide shared-25-mer homology (PHIST), and taxonomic signals
  from the gene annotations.  Weigh the evidence yourself; the profile does not pre-label
  which genes or sections matter most.
Output ONLY valid JSON (no text before or after):
{ "reasoning":  "<brief justification based on the evidence above>",
  "predictions":  [ {"rank":  1, "host":  "<exact name from candidate host list>", "score":  <0.0-1.0>}, ...
]  }
Rules:
- Host names must match exactly as shown in the candidate host list below.
- Include exactly 30 entries, ordered by score descending.
- After the closing } stop immediately.
<PHAGE GENOME PROFILE>    # hashed ID; RBP/BLASTN/25-mer/CRISPR/annotation fields
Top 5 hosts by shared-25-mer count:  ...
CANDIDATE HOST LIST:   1.  <species> ...  223.  <species>
```

| Setting | Value |
|---|---|
| Endpoint | Ollama /api/generate, stream=false, keep_alive=120m |
| Decoding | temperature=0.1, num_predict=4096, num_ctx=40960, num_gpu=99 |
| Stop tokens | <\|endoftext\|>, <\|im_start\|>, <\|im_end\|> |
| Thinking | disabled via the API-level think=false request parameter (the /no_think text prefix in the system prompt has no effect on Qwen3.6:27b); any <think>...</think> stripped post-hoc |
| Output | JSON: reasoning (free text) + predictions (30 × {rank, host, score}) |
| Scoring | score of true host → rank; unlisted host → 0; argmax = top-1 |
| Parsing | tolerant JSON raw-decode; up to 3 attempts; on final failure all-zero scores |
| Error policy | parse-failure / abstention / out-of-catalogue → counted as error |

**Supplementary Table 23.** Statistical analysis methods.

| Quantity | CI / test | resampling unit |
|---|---|---|
| Single proportion / accuracy | Wilson score 95% CI | — |
| Independent-subgroup difference | Newcombe hybrid-score 95% CI | — |
| MRR / MHA / AUROC / PR-AUC / log-loss | percentile bootstrap 95% CI | phage (1,000 resamples, seed 42) |
| Pairwise AUCs | phage-cluster bootstrap | phage (all pairs retained) |
| Paired correctness difference | McNemar (Yates) / Wilcoxon signed-rank | phage |
| Categorical enrichment | Fisher's exact (two-sided) | — |
| Multiple testing | Benjamini–Hochberg where applicable | — |

**Supplementary Table 24.** Dataset construction, source databases and curation for all four benchmarks (Supplementary Note S9).

| Item | RefSeq-634 | VHDB-3150 | Hi-C | EvoMIL (eukaryotic) |
|---|---|---|---|---|
| Source | CHERRY-1940 | Virus–Host DB | MetaHiC human gut | EvoMIL benchmark |
| Origin database | NCBI RefSeq | RefSeq-derived VHDB | proximity-ligation | VHDB (eukaryotic) |
| Reference | Shang 2022 | Shang 2025 | this work | EvoMIL |
| Starting pairs | 1,940 pairs | 4,698 pairs / 498 sp. | 406 phages | 3,198 viruses |
| Evaluation $n$ | 634 phages | 3,150 phages | 406 (Sp-1 46 / Sp-2 82) | 3,198 (shared 3,186) |
| Candidate universe | 223 host species | 488 host species | 52 host species | 36 host species (closed) |
| Partition | held-out eval split | RefSeq-derived eval | Sp-1 / Sp-2 tiers | fold-aware 5-fold CV |
| Same-species control | held out from train | train/test by overlap | — | confined to one fold |
| Reference de-leaking | leave-test-out corpora | leave-test-out corpora | leave-test-out | query + same-species + $\geq$ 95% nbr excluded |
| Identifier masking | hashed phage IDs | hashed phage IDs | hashed IDs | query + homolog IDs hashed |
| Taxonomy normalisation | 17-name normalisation | benchmark taxonomy | MAG-level labels | host-species labels |
| Primary metric | sp@1 (argmax top-1) | sp@1 | MHA-sp/g (Sp-1/Sp-2) | any-hit top-1 |

**Supplementary Table 25.** Compute, runtime and reproducibility of the primary inference pipeline.

| Item | Value |
|---|---|
| Backbone (frozen) | `qwen3.6:27b` (27.8B dense, 4-bit K_M quantisation), no-thinking configuration |
| Serving | Ollama local backend, two shards (ports 11435/11436), `num_gpu=99` |
| Hardware | 2× NVIDIA RTX PRO 6000 Blackwell Max-Q (95.6 GiB VRAM each) |
| Decoding | `temperature=0.1`, `num_ctx=40960`, `num_predict=4096` |
| Concurrency | 6 requests per shard |
| Generated tokens | mean 1,199, max 1,940 per phage |
| Context truncation | 0/634 cutoff (generation and profiles fit within `num_ctx`) |
| Primary run (RefSeq-634) | 634 phages, single pass, completed within one day on the 2-GPU node |
| Determinism | $T = 0.1$ near-deterministic; two independent full-profile runs over byte-identical inputs (configB and the evidence-ablation ladder's own Full condition) agreed on 620/634 phages (97.8%) and gave aggregate sp@1 of 67.4% and 68.3% respectively, a ±0.9-pp run-to-run swing with no fixed seed (Supplementary Table 2 footnote) |
| Fine-tuning | none (training-free; no gradient updates to the backbone) |
| Backbone comparison | Supplementary Table 4 (num_ctx up to 49152 / num_predict up to 40000 for thinking modes) |
| Artifacts | prompts/schema (Table 22), construction parameters (Table 21), Source Data 1–3 |

**Supplementary Table 26.** Genus-level comparison against iPHoP on the isolate benchmarks (iPHoP is GTDB-native with no species-level output; GTDB↔NCBI-harmonised genus top-1, abstention = error; deployed Qwen3.6:27b backbone).

| Dataset | $n$ | iPHoP g@1 (95% CI) | iPHoP coverage | PHI-Reason g@1 (95% CI) | $\Delta$ (PHI − iPHoP) |
|---|---|---|---|---|---|
| RefSeq-634 | 634 | 75.2 (71.7–78.4) | 84.1% | 83.6 (80.5–86.3) | +8.4 |
| VHDB-3150 | 3150 | 50.6 (48.9–52.3) | 80.0% | 78.5 (77.0–79.9) | +27.9 |

**Supplementary Table 27. Prospective leak-controlled case study (14 recently deposited phage genomes; 10 clean post-cutoff, 4 temporally flagged), deployed Qwen3.6:27b backbone.** Each genome was run through the identical frozen-model full-profile interface under full-profile and parametric-only (evidence-stripped) conditions. ⋆ marks the four cases discussed in the main text. Aggregate: full genus 11/14, species 6/14; parametric-only genus 2/14, species 1/14.

| | Phage | Accession | True host | BN | Released | Leak | Full top-1 | sp/gn | Parametric top-1 | sp/gn |
|---|---|---|---|---|---|---|---|---|---|---|
| | Va260-JW1 | PX609837 | *Vibrio_alginolyticus* | - | 2025-12-07 | clean | *Vibrio_cholerae* | ×/✓ | *Vibrio_cholerae* | ×/✓ |
| ⋆ | Skif1059 | PX093639 | *Klebsiella_pneumoniae* | - | 2025-10-08 | clean | *Oenococcus_oeni* | ×/× | *Bacillus_subtilis* | ×/× |
| ⋆ | P16 | PX138265 | *Streptococcus_thermophilus* | + | 2026-01-28 | clean | *Streptococcus_thermophilus* | ✓/✓ | *Bacillus_subtilis* | ×/× |
| ⋆ | DeluluLabubu | PZ236371 | *Streptomyces_griseus* | - | 2026-07-06 | clean | *Lactobacillus_casei* | ×/× | *Mycolicibacterium_smegmatis* | ×/× |
| | JC53 | PX563682 | *Yersinia_pestis* | - | 2026-01-13 | clean | *Providencia_stuartii* | ×/× | *Escherichia_coli* | ×/× |
| ⋆ | Henuyfy22N | PX055572 | *Escherichia_coli* | - | 2025-10-06 | clean | *Escherichia_coli* | ✓/✓ | *Lactococcus_lactis* | ×/× |
| | vB_EfaS_VL6 | PZ133968 | *Enterococcus_faecalis* | + | 2026-04-15 | clean | *Enterococcus_faecalis* | ✓/✓ | *Enterococcus_faecalis* | ✓/✓ |
| | FUzhou-2025 | PV815418 | *Vibrio_harveyi* | - | 2025-10-01 | clean | *Vibrio_cholerae* | ×/✓ | *Synechococcus_sp._WH_8102* | ×/× |
| | S-RIP4a | PV872382 | *Synechococcus_sp._WH_7803* | - | 2026-06-20 | clean | *Synechococcus_sp._WH_8102* | ×/✓ | *Escherichia_coli* | ×/× |
| | Wigberry | PZ319873 | *Streptomyces_lividans* | - | 2026-05-03 | clean | *Streptomyces_venezuelae* | ×/✓ | *Clostridioides_difficile* | ×/× |
| | XWef1 | PV491271 | *Enterococcus_faecalis* | - | 2025-09-20 | **pre-cut** | *Enterococcus_faecium* | ×/✓ | *Bacillus_subtilis* | ×/× |
| | JB8 | PQ473534 | *Pseudomonas_aeruginosa* | - | 2024-11-14 | **pre-cut** | *Pseudomonas_aeruginosa* | ✓/✓ | *Escherichia_coli* | ×/× |
| | LPPA15 | PP895303 | *Pseudomonas_aeruginosa* | - | 2026-06-06 | embargo* | *Pseudomonas_aeruginosa* | ✓/✓ | *Escherichia_coli* | ×/× |
| | SJW01A | PQ074102 | *Salmonella_enterica* | + | 2026-06-17 | embargo* | *Salmonella_enterica* | ✓/✓ | *Escherichia_coli* | ×/× |

**Supplementary Table 28. Typed-decision inference-interface transfer (Jev, TypeSafe AI).** Species-level top-1 accuracy (sp@1) under Jev's `choice` primitive on the unmodified RefSeq-634, VHDB-3150 and HiC full-evidence profiles, compared with the deployed Qwen3.6:27b backbone (Supplementary Table 1). No profile content, evidence schema or candidate catalogue was changed between the two inference interfaces (Methods); full HiC multi-host accuracy is given in Supplementary Table 32.

| Dataset | $n$ | Jev sp@1 (95% CI) | Qwen3.6:27b sp@1 |
|---|---|---|---|
| RefSeq-634 | 634 | 68.3 (64.6–71.8) | 67.4 (63.6–70.9) |
| VHDB-3150 | 3,150 | 54.8 (53.0–56.5) | 57.2 |
| HiC Sp-1 | 46 | 50.0 | 52.2 |
| HiC Sp-2 | 82 | 68.3 | 68.3 |

**Supplementary Table 29. Typed-decision inference-interface transfer (Jev) — HiC multi-host accuracy.** Full species- and genus-level ranking metrics for Jev's `choice` and `noul` primitives on the unmodified HiC full-evidence profiles, with PHI-Reason (deployed Qwen3.6:27b) MHA-sp/MHA-g reproduced from Supplementary Table 19 for reference.

| Split | Method | $n$ | sp@1 | sp@5 | g@1 | g@5 | MRR | MHA-sp | MHA-g |
|---|---|---|---|---|---|---|---|---|---|
| Sp-1 | Jev (`choice`) | 46 | 0.500 | 0.826 | 0.935 | 0.978 | 0.648 | 0.375 | 0.857 |
| Sp-1 | Jev (`noul`) | 46 | 0.522 | 0.891 | 0.935 | 1.000 | 0.691 | 0.483 | 0.911 |
| Sp-1 | PHI-Reason (Qwen3.6:27b) | 46 | — | — | — | — | — | 0.517 | 0.893 |
| Sp-2 | Jev (`choice`) | 82 | 0.683 | 0.866 | 0.951 | 0.988 | 0.769 | 0.622 | 0.929 |
| Sp-2 | Jev (`noul`) | 82 | 0.659 | 0.939 | 0.902 | 1.000 | 0.772 | 0.643 | 0.906 |
| Sp-2 | PHI-Reason (Qwen3.6:27b) | 82 | — | — | — | — | — | 0.673 | 0.976 |

**Supplementary Table 30. Typed-decision vs. reasoning-engine inference latency.** Wall-clock, per-query inference time for Jev's `choice` primitive versus the deployed Qwen3.6:27b reasoning engine, under matched serial, single-query execution, on the unmodified RefSeq-634, HiC and VHDB-3150 full-evidence profiles (Methods).

| Dataset | Jev latency (s) | Qwen3.6:27b latency (s) | Approx. speedup |
|---|---|---|---|
| RefSeq-634 | 0.69 | 27.8 | ∼40× |
| HiC (Sp-1+Sp-2) | 0.60 | 19.7 | ∼33× |
| VHDB-3150 | 2.23 | 23.7 | ∼11× |

**Supplementary Table 31.** **External 207-pair wet-lab-supported validation.** **a**, Overall accuracy. **b**, Accuracy by confirmed host species (12 most frequent). **c**, Evidence availability across the cohort.

**a**

| | $k = 1$ | $k = 3$ | $k = 5$ | $k = 10$ | $k = 30$ |
|---|---|---|---|---|---|
| Species top-$k$ (%) | 52.2 | 68.6 | 82.1 | 91.8 | 97.6 |
| Genus top-$k$ (%) | 90.8 | 96.1 | 97.6 | 98.1 | 100.0 |

**b**

| Confirmed host | $n$ | sp@1 | gn@1 | Confirmed host | $n$ | sp@1 | gn@1 |
|---|---|---|---|---|---|---|---|
| *Klebsiella pneumoniae* | 52 | 90.4 | 96.2 | *Phocaeicola dorei* | 5 | 0.0 | 0.0 |
| *Bacteroides caccae* | 33 | 27.3 | 100.0 | *Proteus mirabilis* | 5 | 100.0 | 100.0 |
| *Escherichia coli* | 22 | 45.5 | 72.7 | *Bifidobacterium adolescentis* | 4 | 100.0 | 100.0 |
| *Bacteroides thetaiotaomicron* | 8 | 0.0 | 100.0 | *Phocaeicola plebeius* | 4 | 50.0 | 50.0 |
| *Bacteroides faecis* | 8 | 37.5 | 100.0 | *Enterococcus faecium* | 4 | 75.0 | 100.0 |
| *Clostridium perfringens* | 5 | 100.0 | 100.0 | *Bacteroides ovatus* | 4 | 25.0 | 100.0 |

**c**

| Evidence | $n/207$ | % |
|---|---|---|
| CRISPR record present | 138 | 66.7 |
| BLASTN neighbour matching confirmed host | 153 | 73.9 |
| BLASTN host signal STRONG | 164 | 79.2 |
| Median identity / coverage of confirmed-host BLASTN hits | 96.8% / 31% | |

All 207 queries were resubmitted through the unchanged production pipeline against ∼494 candidate host species; 207/207 parsed successfully (abstentions would be counted as errors). Source cohorts: temperate gut-prophage isolation study (134), KlebPhaCol (52), broad-host-range *Vibrio* study (18) and three single-genome studies (1 each).

**Supplementary Table 32.** **Structured-evidence perturbation on the 207-pair cohort:** field-removal and label-masking conditions (7 conditions, 1,449 inference calls).

| Condition | $n$ scored | sp@1 (%) | sp@5 (%) | Δsp@1 (pp) |
|---|---|---|---|---|
| Full profile | 207 | 52.2 | 82.1 | — |
| −25-mer | 207 | 54.1 | 79.7 | +1.9 |
| −RBP | 207 | 46.4 | 81.6 | -5.8 |
| BLASTN host label masked | 207 | 43.0 | 77.8 | -9.2 |
| −CRISPR | 207 | 38.6 | 72.0 | -13.5 |
| −BLASTN | 207 | 34.8 | 54.1 | -17.4 |
| No structured evidence | 207 | 18.8 | 31.9 | -33.3 |

Each condition modifies only the targeted evidence block; model, prompt, decoding settings and candidate catalogue are identical throughout. Δ is relative to the full profile. Every condition is scored over the complete set of 207 queries, a query being counted as incorrect whenever the confirmed host is not returned at rank 1, so all conditions share one denominator and are directly comparable to the full-profile accuracy reported in the main text.

### Supplementary Figures

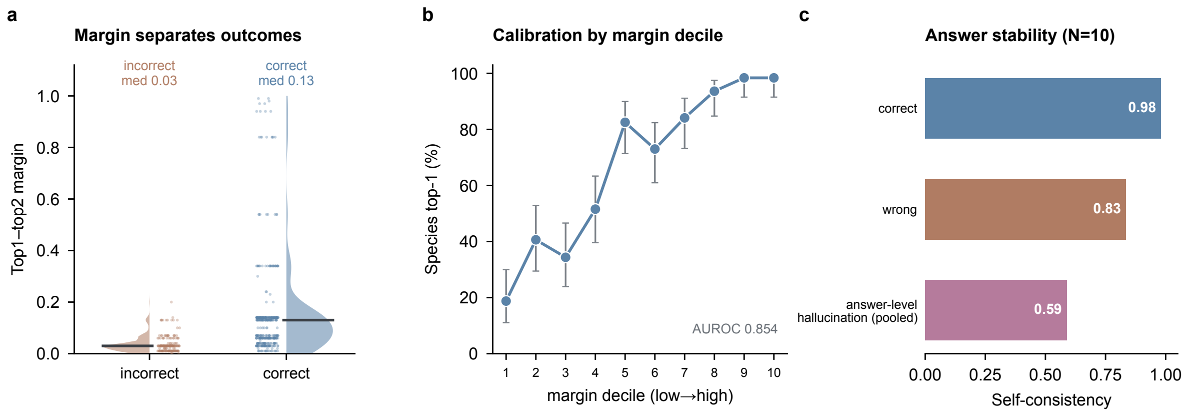

**Supplementary Figure 1.** **Reliability of the top1–top2 margin (RefSeq-634).** (**a**) Margin distribution by outcome (half-violin with jittered points and median bar): correct predictions carry a substantially larger margin (median 0.13) than incorrect ones (median 0.03). (**b**) Calibration by margin decile: observed species top-1 accuracy rises monotonically from the lowest to the highest margin decile (margin AUROC 0.854; 95% Wilson CIs). (**c**) Answer stability under resampling ($N = 10$, $T = 0.7$): correct predictions are highly self-consistent (0.98) versus 0.84 for incorrect predictions and 0.59 for the pooled answer-level-hallucination cases ($n = 19$).

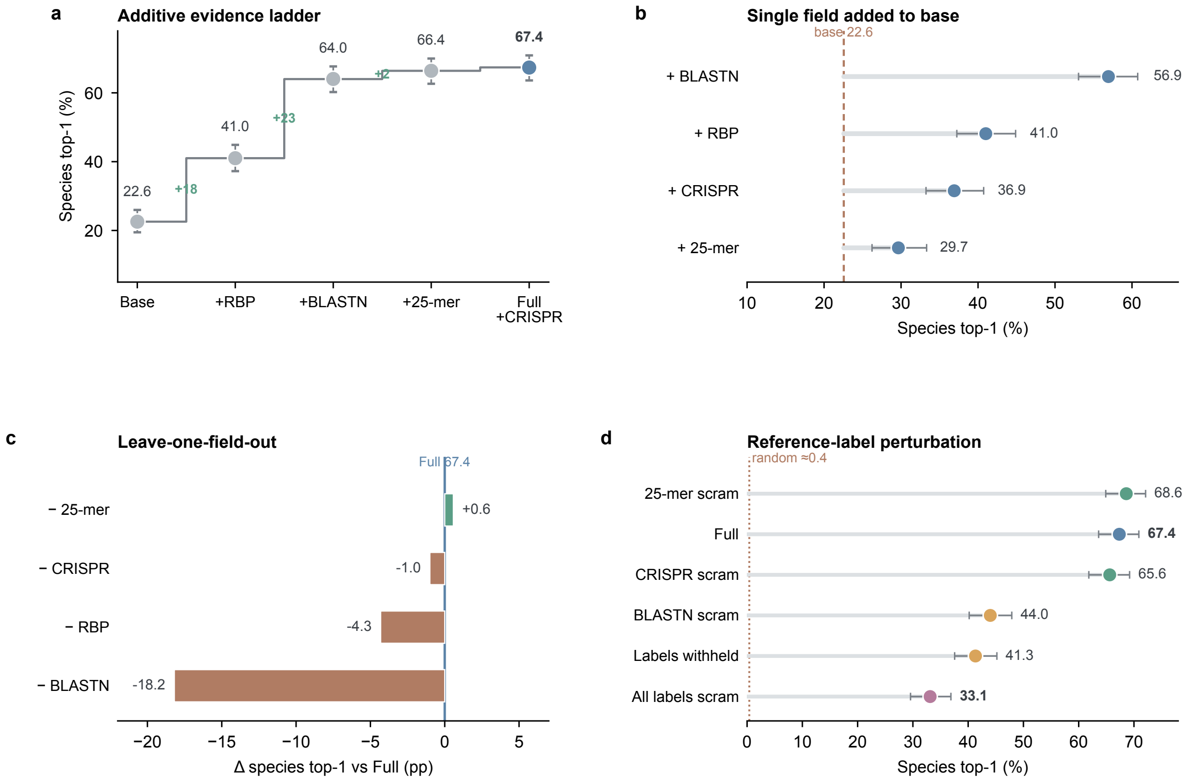

**Supplementary Figure 2. Structured-evidence perturbation (RefSeq-634, $n = 634$).** (**a**) Additive evidence ladder: species top-1 rises from an annotation-only base (22.6%) to the full profile (67.4%) as fields are added (95% Wilson CIs; green = per-step gain in pp). (**b**) Single field added to the base profile (Cleveland dots vs the base reference line): BLASTN contributes most in isolation. (**c**) Leave-one-field-out from the full profile (centred on Full): removing BLASTN is by far the most damaging (−18.1 pp). (**d**) Reference-label perturbation (ordered lollipops): withholding or scrambling host labels substantially reduces accuracy (to 33.1–41.3%, still well above the ≈0.4% random floor), whereas scrambling the low-specificity 25-mer and sparse CRISPR labels does not, showing the model tracks evidence content rather than field presence.

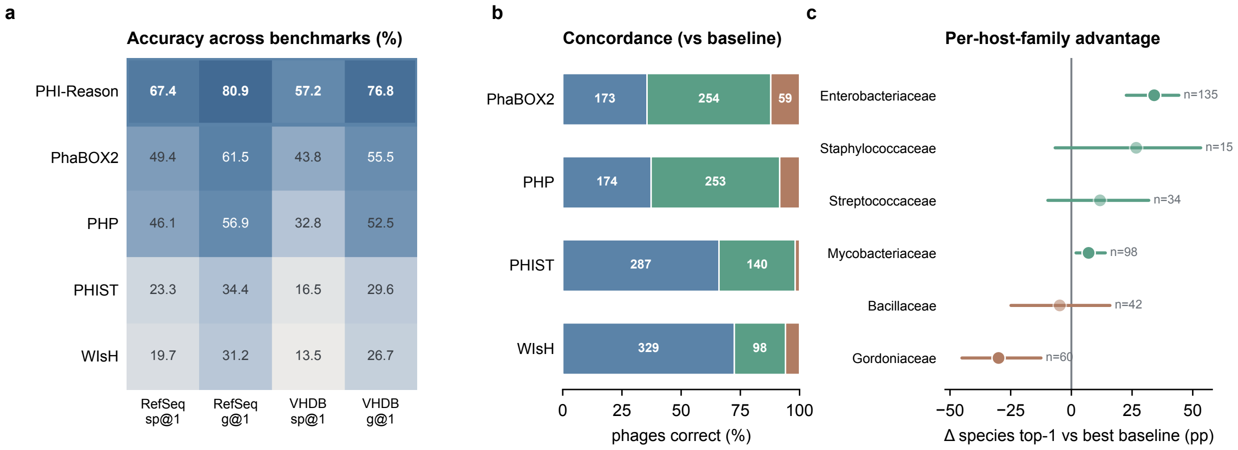

**Supplementary Figure 3. Cross-benchmark operating envelope.** (**a**) Method × benchmark accuracy heatmap (species and genus top-1 on RefSeq-634 and VHDB-3150); the PHI-Reason row is outlined. (**b**) Per-phage concordance against each baseline (100% stacked): PHI-Reason recovers many phages the baselines miss (PHI-only, blue) with few losses (baseline-only, clay). (**c**) Per-host-family advantage: PHI-Reason species top-1 minus the strongest baseline with Newcombe 95% CIs (faded points cross zero); the method leads on the larger families (e.g. *Enterobacteriaceae*) and trails on some Actinobacteria (e.g. *Gordoniaceae*).

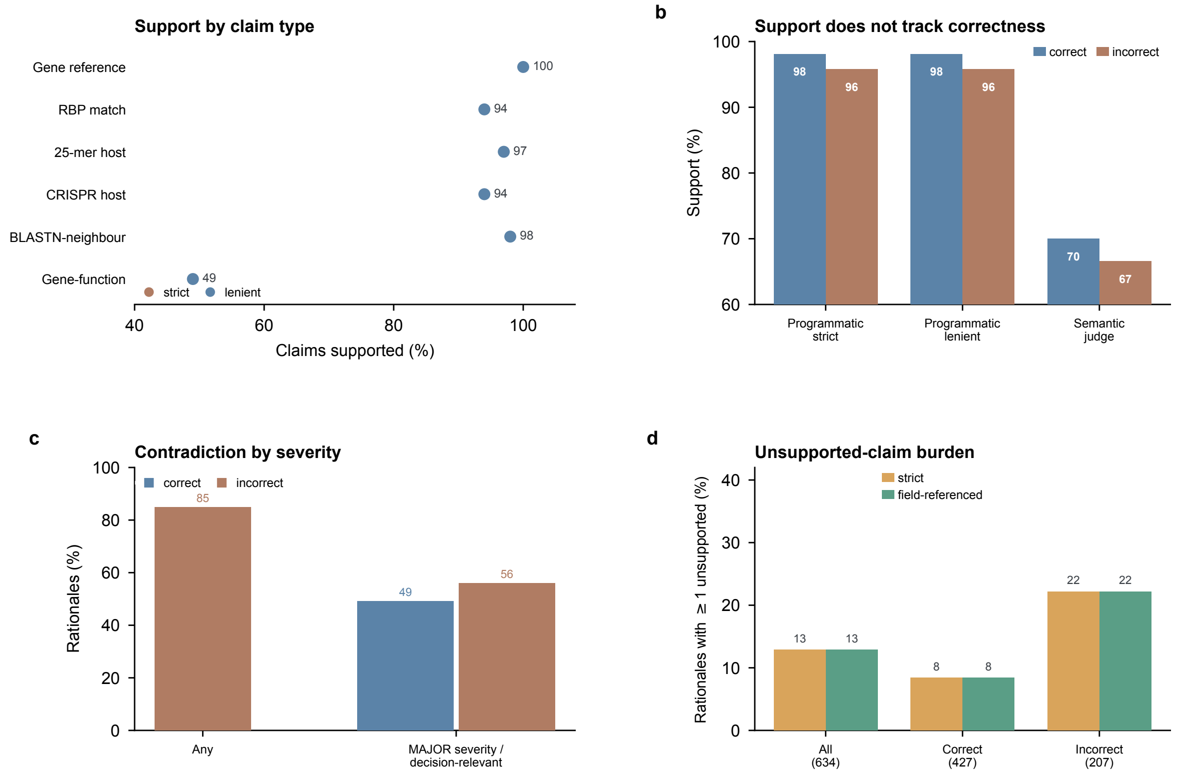

**Supplementary Figure 4. Hallucination verifier and judge — extended breakdowns (RefSeq-634; 427 correct, 207 incorrect; deployed Qwen3.6:27b backbone).** This figure provides the per-claim-type, per-severity and per-definition detail underlying main-text Fig. 3 and Supplementary Tables 16–17. (**a**) Claim-level support by claim type under the strict (cited-field) and lenient (entity-level) criteria: strict and lenient are identical for every claim type under this backbone (Supplementary Table 16). (**b**) Programmatic (strict/lenient) and blinded-judge support are similar for correct and incorrect predictions, i.e. grounding does not track correctness. (**c**) Contradiction rate by severity and outcome: MAJOR-severity contradictions, which coincide with decision-relevant contradictions under this rubric, are only modestly enriched among errors (56.0% incorrect versus 49.2% correct). (**d**) Rationale-level unsupported-claim burden under the strict versus field-referenced definitions, by prediction group.

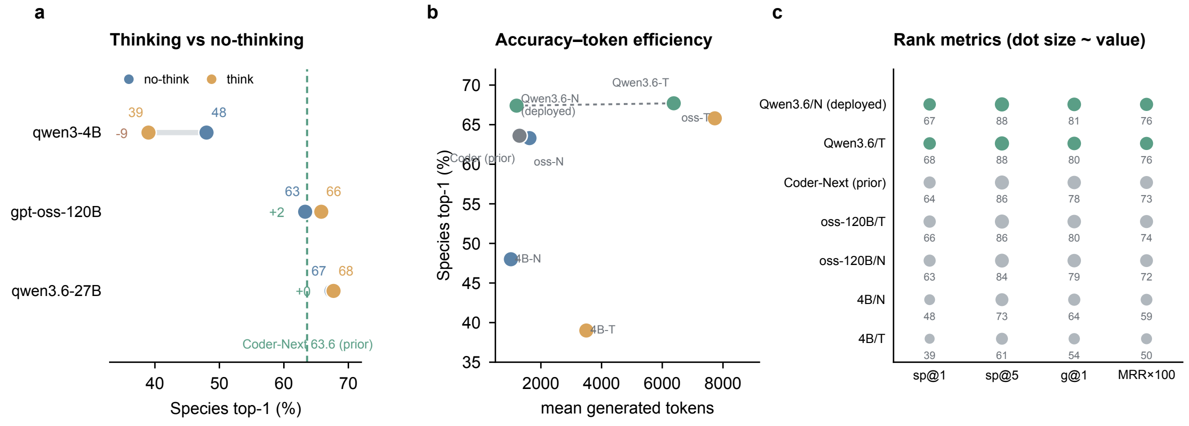

**Supplementary Figure 5. Backbone and thinking-mode behaviour (RefSeq-634, $n = 634$).** (**a**) Thinking versus no-thinking species top-1 (dumbbells) for qwen3-4B and gpt-oss-120B, with the deployed Qwen3.6-27B as a reference (dashed): enabling thinking *hurts* the small model ($48.0 \rightarrow 39.0$) but slightly helps the large one ($63.3 \rightarrow 65.8$). (**b**) Accuracy–token efficiency: species top-1 versus mean generated tokens; the deployed Qwen3.6-27B (no-thinking) defines the efficient frontier, reaching 67.4% at 1,199 tokens, ahead of Qwen3-Coder-Next and gpt-oss-120B (no-think), both at a fraction of the token cost of thinking modes (Supplementary Table 4). (**c**) Rank metrics (sp@1, sp@5, g@1, MRR) as dot small-multiples (dot size scales with value); the deployed Qwen3.6-27B leads the other higher-capacity configurations on every metric shown.

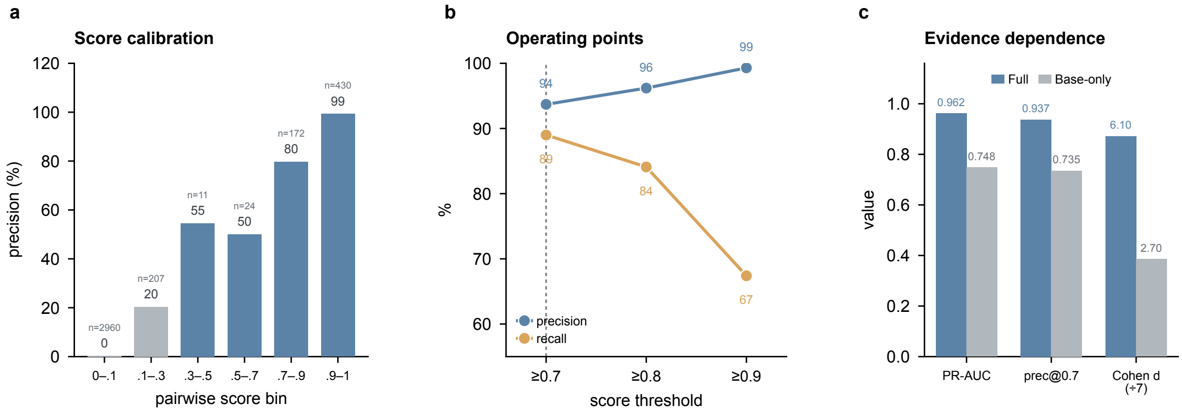

**Supplementary Figure 6. Pairwise verification (RefSeq-634; 634 positive and 3,170 negative pairs).** (**a**) Score-bin precision: precision rises monotonically with the pairwise score, from 0.3% in the lowest bin to 99% in the highest (bin sizes annotated). (**b**) Operating points: precision and recall at score thresholds 0.7/0.8/0.9. (**c**) Evidence dependence: removing the structured evidence (Base-only) sharply degrades separation (PR-AUC 0.962 → 0.748, precision@0.7 0.937 → 0.735, Cohen's $d$ 6.10 → 2.70; pairwise prompt provenance in Methods and Note S6).

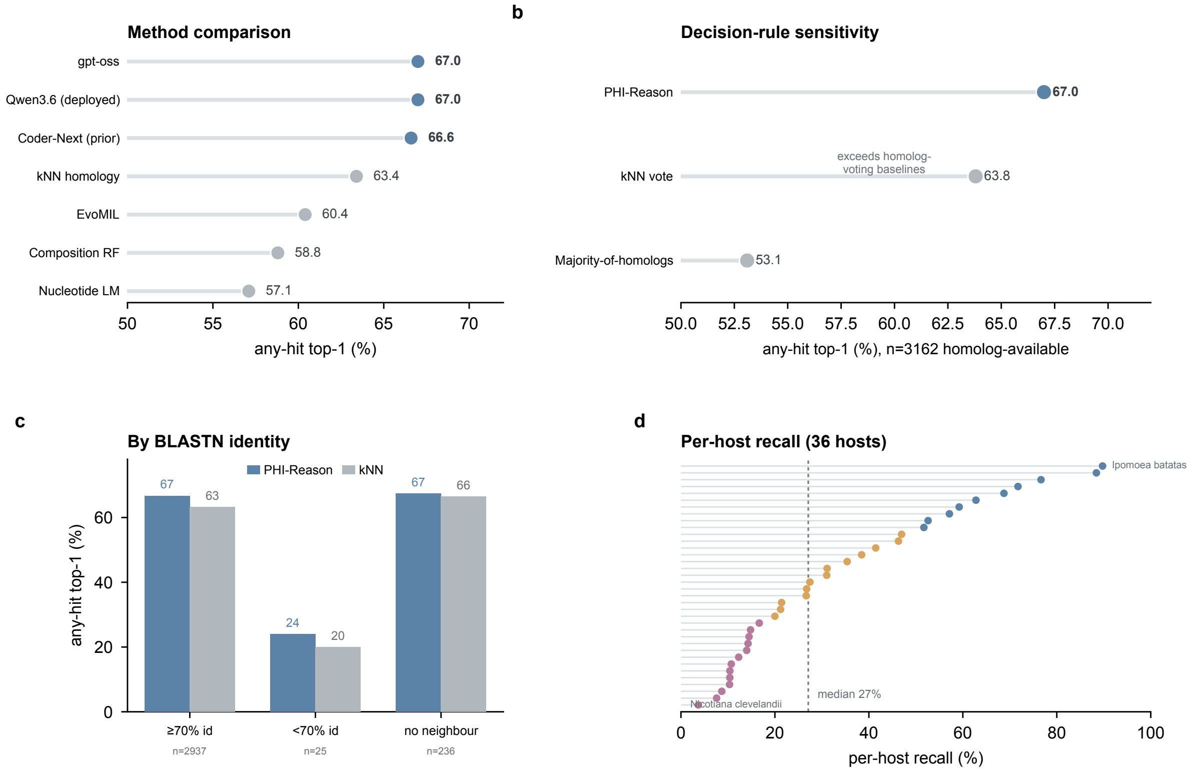

**Supplementary Figure 7. Eukaryotic virus–host case study (EvoMIL 36-host closed set).** (**a**) Method comparison (any-hit top-1): all three PHI-Reason backbones lead the learned baselines and the kNN homology baseline. (**b**) Decision-rule sensitivity on the homolog-available subset ($n = 3{,}162$): PHI-Reason exceeds both the kNN-vote and majority-of-homologs baselines, i.e. it does not collapse to a homolog-voting lookup. (**c**) Accuracy by whole-genome BLASTN identity band: PHI-Reason edges kNN across the strong-, weak- and no-BLASTN-neighbour strata. (**d**) Per-host recall for all 36 hosts (sorted); the low-recall tail is dominated by within-clade confusion (e.g. *Nicotiana*, *Cercopithecidae*), while any-hit accuracy remains high.

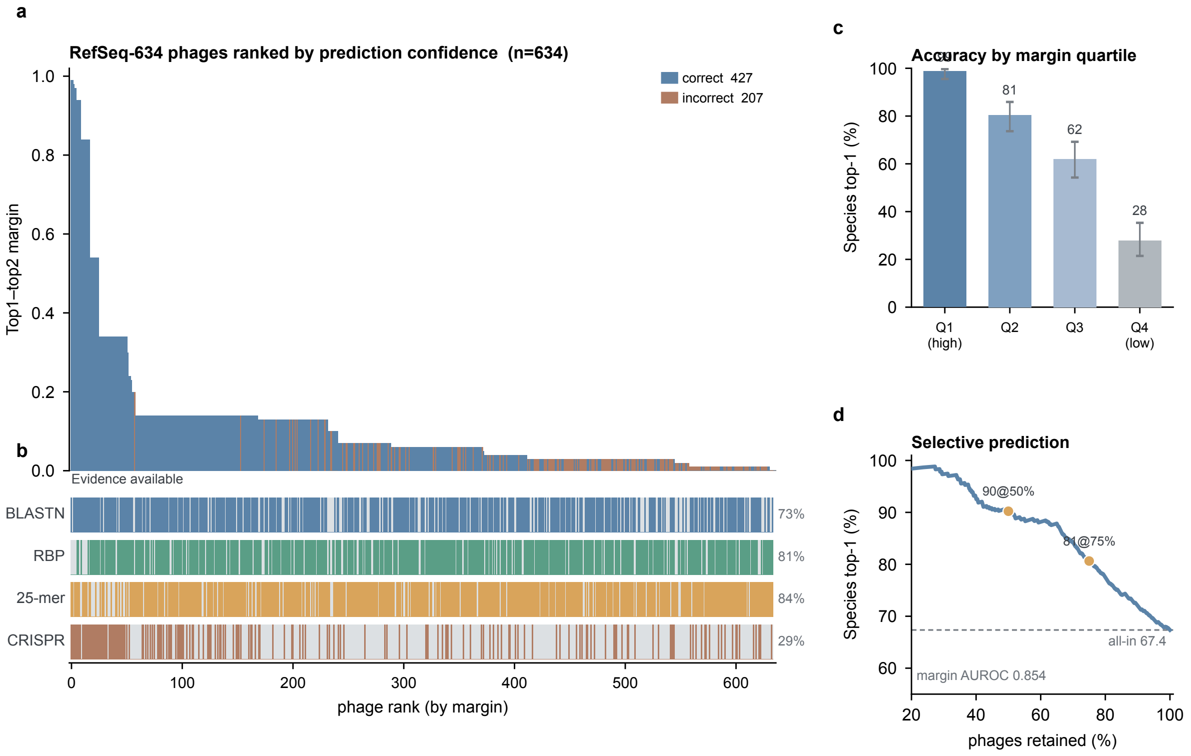

**Supplementary Figure 8. The 634-phage evidence-and-confidence landscape.** (**a**) All 634 RefSeq-634 phages ranked by top1–top2 margin; each bar is one phage, coloured by outcome. High-confidence predictions are almost all correct (blue); errors (clay) concentrate at low margin. (**b**) Evidence-availability rug aligned to (a): BLASTN, RBP and 25-mer support is dense among high-margin phages and thins toward the low-margin tail; CRISPR is sparse throughout. (**c**) Species top-1 accuracy by margin quartile (99% → 28% from the most to least confident quartile). (**d**) Selective prediction: retaining only higher-margin predictions raises accuracy (e.g. 90% at 50% coverage), the operational payoff of the margin-based reliability signal. This single view captures the method's central behaviour: retrievable evidence and prediction confidence jointly organise the outcome landscape.

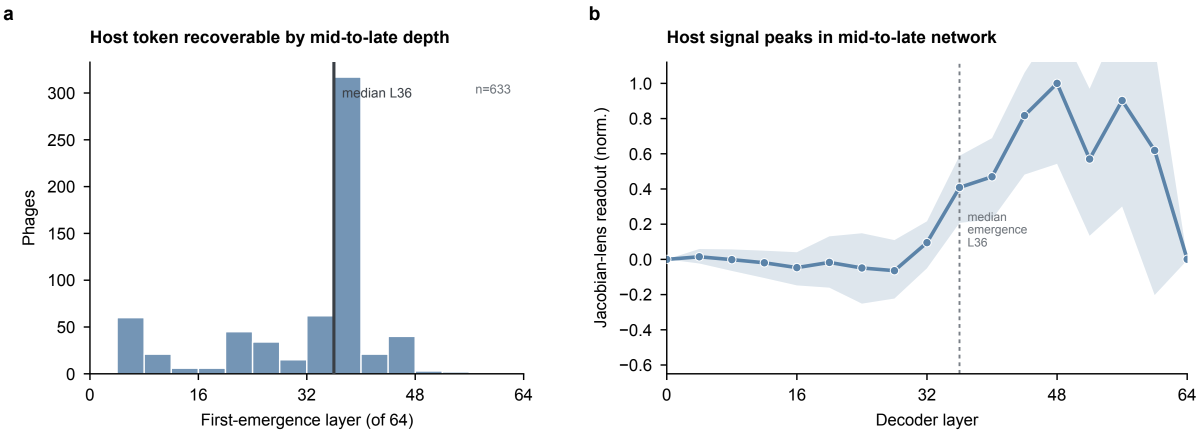

**Supplementary Figure 9. Layerwise emergence under the target-conditioned local Jacobian readout (RefSeq-634; Supplementary Note S11).** (**a**) Distribution across phages of the first decoder block at which the confirmed host-genus logit acquires an appreciable positive first-order contribution ($r_\ell$ reaches 20% of its positive peak): median layer 36 in the 64-layer model (56% of network depth). (**b**) Median normalised $r_\ell$ of the confirmed host token across decoder blocks (shaded band, interquartile range; normalised to the maximum of the median curve): the attribution rises from early blocks and peaks in mid-to-late network depth. The read direction is the ground-truth host logit; this is a descriptive, target-conditioned attribution analysis that yields no vocabulary ranking and makes no causal claim about the computation used during generation.

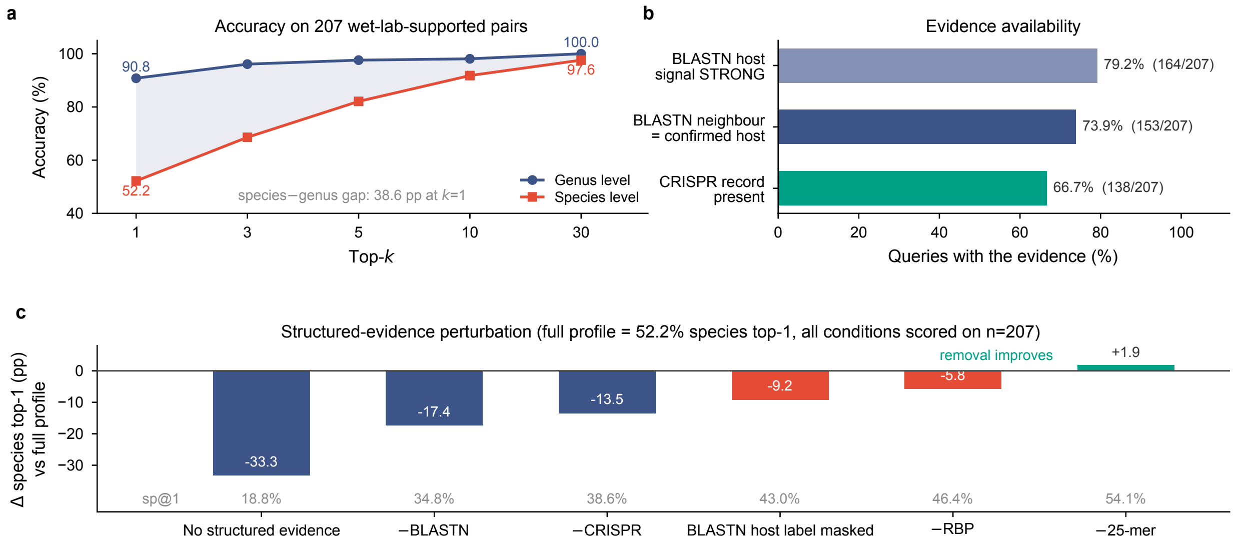

**Supplementary Figure 10. External wet-lab-supported validation on 207 phage–host pairs** (Supplementary Note S13; Supplementary Tables 31–32). (**a**) Species- and genus-level top-$k$ accuracy against ∼494 candidate host species. The 38.6 pp species–genus gap at $k$=1 closes to 2.4 pp by $k$=30: the confirmed host is almost always retrieved, and the residual error is a ranking problem within the correct genus rather than a retrieval failure. (**b**) Evidence actually retrieved across the cohort. (**c**) Field-removal and label-masking perturbation, as the change in species top-1 relative to the full profile (grey row gives the absolute sp@1 of each condition). All conditions are scored over the full set of 207 queries. Removing all structured evidence costs 33.3 pp; masking only the host *labels* on otherwise intact BLASTN records costs 9.2 pp, i.e. 53% of the 17.4 pp lost by deleting those records outright, so both the host annotation and the alignment statistics carry signal. The 25-mer channel is the single condition whose removal *improves* accuracy (teal, +1.9 pp).

### Supplementary Notes

#### Supplementary Note S1 | Gram-type and reference-coverage stratification

Full-profile RefSeq-634 sp@1 by host Gram type and BLASTN-neighbour availability (Wilson CIs, Supplementary Table 7): Gram-positive 71.4% versus Gram-negative 63.3%; neighbour present 72.7% ($n = 469$) versus absent 52.1% ($n = 165$), difference 20.6 pp (95% CI 12.0–29.1). The RBP gain is larger for Gram-negative (+30.3 pp) than Gram-positive (+9.3 pp; Table 7b).

#### Supplementary Note S2 | Adversarial reference-label scrambling controls

Host labels were withheld or permuted in the BLASTN, 25-mer and CRISPR blocks, retaining non-label content and the RBP field (full values, Supplementary Table 6; ordered effects, Supplementary Figure 2). Withholding labels gave 41.3% (−26.0 pp); scrambling all blocks gave 33.1% (−34.2 pp, above the ≈0.4% random floor for 223 candidates). BLASTN-label scrambling alone was the most proportionally damaging single-field perturbation (−23.3 pp).

#### Supplementary Note S3 | Reasoning-trace citation dimensions

Rationales were parsed into five citation channels (RBP, BLASTN, 25-mer, CRISPR, taxonomy/annotation); cross-backbone frequencies (Supplementary Figure 5) are correlational, not causal.

#### Supplementary Note S4 | Cross-model reasoning traces and thinking mode

Across seven backbone–mode configurations (Supplementary Table 4), enabling thinking increased output length (GPT-oss-120B 1,624→7,731 tokens; Qwen3-4B 1,012→3,497; Qwen3.6-27B 1,199→6,378) without a consistent accuracy gain (+2.5, −9.0, +0.3 pp respectively). BLASTN-citation rate ranged from 86.4% (Qwen3-Coder-Next) to 9.5–15.5% (Qwen3-4B), with the deployed Qwen3.6-27B intermediate at 80.4–82.0%.

#### Supplementary Note S5 | Shared failures and evidence-concordance stratification

*Shared failures.* CBI ($n = 149$, missed by all seven configurations) and BMD ($n = 196$) overlapped in 89 phages (OR 5.24, 95% CI 3.54–7.75, Fisher's exact $p = 4.6 \times 10^{-17}$; OR 5.49 on the 532-phage valid-prediction subset; Supplementary Table 8).

*Evidence concordance.* Profiles were classified by whether host-linking channels (BLASTN, RBP, 25-mer, CRISPR) agreed and supported the confirmed genus. Genus accuracy was below 15% for every backbone on the $n = 64$ conflicting/no-support profiles, versus 98–100% (three higher-capacity configurations) on the $n = 198$ concordant profiles. Because strata are defined using the confirmed host, this is an association, not a prospective reliability test.

#### Supplementary Note S6 | Pair-level verification

Full-profile pairwise PR-AUC 0.962, ROC-AUC 0.992, Cohen's $d = 6.10$; the candidate-density control raised listwise sp@1 from 67.4% to 95.7%, close to pairwise Hits@1 (97.2%; McNemar not recomputed). The original pairwise-prompt script could not be located; the Qwen3.6:27b prompt used here was rebuilt from the Methods schema and was not verified verbatim against it, so numerical comparison with pairwise results reported for other backbones should be treated as approximate (Supplementary Tables 9–10, Supplementary Figure 6).

#### Supplementary Note S7 | Prompts, output schemas and decoding settings

Schematic prompts, JSON schemas, decoding parameters, retry and parse-failure policy are summarised in Supplementary Table 22; the complete unabridged prompts, schemas and parser are provided in the code repository (Code availability).

#### Supplementary Note S8 | Evidence verification and uncertainty analyses

##### S8.1 Evidence digest construction

The evidence digest was parsed from the exact model-input profiles, without rerunning BLASTN or RBP retrieval under separate thresholds (digest fields and thresholds, Supplementary Table 21: RBP identity $\geq 40\%$/coverage $\geq 5\%$/e-value $\leq 10^{-5}$, top-5 genera per gene; BLASTN identity $\geq 70\%$/coverage $\geq 3\%$; 25-mer $k = 25$; CRISPR $\geq 90\%$ identity/length, $\leq 2$ mismatches).

##### S8.2 Programmatic verifier: strict and lenient matching

3,365 field-referenced claims (5.3 per rationale), 96.4% strict field-specific support, identical to the lenient entity-level value for every claim type (Supplementary Table 16; rationale-level burden, Supplementary Figure 4d).

##### S8.3 Semantic judge and contradiction detection

The blinded semantic judge (Claude Opus 4.8, fixed rubric) flagged 51.4% of rationales with $\geq 1$ decision-relevant (MAJOR) contradiction (49.2% correct versus 56.0% incorrect). Targeted re-auditing found unsupported specific details – invented gene counts, schema fields, neighbour-phage identifiers, candidate-host genera – not captured by the programmatic checks.

##### S8.4 Judge prompt and schema

The blinded semantic judge (Claude Opus 4.8, `claude-opus-4-8`, Anthropic API) received each rationale with the profile fields and returned per-claim support labels and a contradiction flag; the host prediction was withheld. Verbatim prompt and rubric are in the repository (Data Availability); the host-list prediction prompt is in Supplementary Table 22.

##### S8.5 Score margin and selective prediction

Margin AUROC 0.854 (95% CI 0.824–0.883; RefSeq-634) / 0.823 (95% CI 0.761–0.877; Balanced-200); mean margin 0.146 (correct) versus 0.036 (wrong) versus 0.030 (answer-level hallucination, $n = 19$; Supplementary Table 18, Supplementary Figure 1). Joint evidence-margin organisation across all 634 phages: Supplementary Figure 8.

##### S8.6 Resampling self-consistency

Self-consistency ($N = 10$, $T = 0.7$) AUROC 0.672: 0.981 (correct) versus 0.835 (wrong) versus 0.589 (pooled answer-level hallucination, $n = 19$; Supplementary Table 18).

##### S8.7 Answer-level error-source decomposition

Off-evidence 19/634 (3.0%; 95% CI 1.9–4.6%); the 207 errors partition into 157 non-decisive, 48 evidence-incomplete, 2 recoverable (Supplementary Table 17).

##### S8.8 Statistical analyses

Methods summarised in Supplementary Table 23.

#### Supplementary Note S9 | Data provenance, curation and leakage controls

*Dataset construction.* Four benchmarks (sources, versions, de-duplication, taxonomy handling: Supplementary Table 24): RefSeq-634 (held-out CHERRY-1940 partition), VHDB-3150 (3,150 evaluation phages), Hi-C MetaHiC (52 host species, 406 phages), EvoMIL (36 eukaryotic hosts). All reference corpora were built leave-test-out and query-de-leaked (Supplementary Tables 5, 21).

*Leakage controls.* The backbone's pretraining corpus is undisclosed, so genome-level exposure cannot be excluded a priori; we bound rather than deny it. (i) Profiles carry no accession or isolate name – headed by a content hash, with reference neighbours hashed and flagged "not in DB." (ii) Withholding host labels drops sp@1 to 41.3%, and scrambling all labelled blocks to 33.1% (Note S2): accuracy tracks evidence content, not presence. (iii) The annotation-only Base floor is 22.6% (Table 2). (iv) Without a BLASTN neighbour, sp@1 is 52.1% versus 72.7% with one – degrading gracefully rather than as label recall would predict. (v) On the eukaryotic transfer, PHI-Reason exceeds the kNN-vote and majority-of-homologs rules rather than collapsing to homolog lookup (Table 14b).

*Reporting.* Generation-cutoff rate 0/634 (mean 1,199 tokens, max 1,940; 4,096-token budget, 40,960-token context) – no profile was truncated. Per-item audits: Source Data 1–3; prompts, schemas, parsers, catalogues and code: repository (Data Availability); model file and compute: Supplementary Table 25.

#### Supplementary Note S10 | Eukaryotic virus–host portability case study

We tested portability to a distinct domain without interface changes, using the EvoMIL 36-host eukaryotic virus–host benchmark as a compact probe (the framework was not tuned to this task). Qwen3-Coder-Next, gpt-oss:120b and the baselines were compared on the shared 3,186-virus subset of the full 3,198-virus set; Qwen3.6:27b was evaluated on the full set and is reported separately (Supplementary Tables 12, 14). Each virus maps to one or more true hosts from a fixed 36-host catalogue; any-hit top-1 credits a match to any of them.

*Controls and schema.* Fold-aware five-fold cross-validation kept same-species viruses in one fold; retrieval excluded the query, same-species entries and near-identical ($\geq 95\%$) neighbours, with identifiers hash-masked. Only the evidence schema changed: phage-specific modules (RBP, 25-mer, CRISPR) were replaced by virus–host fields (Baltimore class, segmentation, taxonomy, BLASTP homology; Supplementary Table 13), with interface, backbone and decoding fixed. A floor profile without homology or taxonomy reached 19.2–35.1% any-hit top-1 (Qwen3.6:27b 19.2%, Qwen3-Coder-Next 23.7%, gpt-oss:120b 35.1%), 32–48 pp below the corresponding full profiles (Supplementary Table 14).

*Results.* With the full profile, PHI-Reason reached any-hit top-1 67.0% (strict 61.9%) with Qwen3.6:27b on 3,198 viruses; on the shared 3,186-virus set, 67.0% (strict 61.7%) with gpt-oss:120b and 66.6% (strict 61.5%) with Qwen3-Coder-Next (Supplementary Table 14), exceeding the evaluated baselines (EvoMIL 60.4%, kNN homology 63.4%, nucleotide-LM 57.1%, composition 58.8%). Under the Qwen3-Coder-Next backbone, 89.9% of predictions matched the single strongest homolog and 99.0% fell within a neighbour's host set (Source Data 3), so the ported interface operated largely as structured homology-following. BLASTN-identity-stratified and no-neighbour subsets (Supplementary Table 15) and per-host recall (Supplementary Figure 7; Source Data 3) delimit the regime in which portability holds for that backbone.

#### Supplementary Note S11 | Layerwise emergence under a target-conditioned local Jacobian readout

For each RefSeq-634 phage (Full profile), we computed the target-conditioned local Jacobian attribution $r_\ell = \langle \partial z_{\text{host}} / \partial h_\ell,\, h_\ell \rangle$ at the query position preceding host-name generation – the first-order sensitivity of the confirmed host-genus logit to each decoder-block activation (reverse mode, frozen parameters, matched `bf16` checkpoint, sampled across the 64 blocks of the deployed Qwen3.6:27b backbone, three-quarters of which use linear rather than full attention; Methods). Unlike the original context-averaged Jacobian lens, this is a per-input, single-target attribution with no vocabulary ranking; the confirmed genus enters only as the differentiation target. One phage's curve never reached the emergence threshold and was excluded from emergence-layer statistics only.

The confirmed host-genus logit first acquired an appreciable positive contribution at a median layer of 36 of 64 (56% of depth; Supplementary Figure 9), stable across relative thresholds ($\tau = 0.1/0.2/0.3/0.5$: median layer 32/36/36/44), and crossed later for correct (median 36) than incorrect (median 20) predictions. In the mid-to-late window (layers 32–53), correct predictions showed stronger mean positive attribution than incorrect (0.88 versus 0.36); we do not attribute this to a causal mechanism.

These are descriptive differences in a linear, non-causal readout, not evidence of an internal decision. Because the read direction is the ground-truth host logit and host genera appear verbatim in the candidate catalogue and evidence fields, early attribution cannot separate lexical availability from evidence integration; occurrence-matched lexical nulls and perturbation-matched analyses are left to future work.

#### Supplementary Note S12 | Prospective leak-controlled case study on recently deposited phage genomes

*Design.* To test whether evidence-grounding extends to genuinely unseen genomes, we assembled 14 GenBank phages whose host falls within the RefSeq-634 223-species catalogue and processed each through the identical frozen-model interface (same backbone, prompt, decoding, candidate presentation and full evidence profile). This is a temporal evidence-attribution control, not a powered accuracy estimate: 14 genomes carry wide Wilson intervals; the aim is to show correct calls track the supplied named evidence, arguing against a purely memorisation-driven account.

*Leakage control.* Accession, host, release/submission dates and reference-neighbour status are in Supplementary Table 27. Using a conservative cutoff of 2025-09-30, ten genomes (including all four main-text vignettes) were released after it with no near-full-length reference duplicate; four are flagged as *supporting* rather than strictly leak-free (two released shortly before the cutoff, two submitted in 2024 under embargoed release). Illustrative cases are drawn only from the ten cleanly post-cutoff genomes.

*Evidence attribution.* With the deployed Qwen3.6:27b backbone, the full profile recovered the correct genus for 11/14 genomes (95% CI 52–92%) and species for 6/14 (21–67%). Parametric-only fell to genus 2/14, species 1/14 (the one residual species-correct call, *E. faecalis*, being a textbook pair recoverable from priors); removing only RBP gave genus 5/14, species 4/14, intermediate between full and parametric-only. On the eleven neighbour-absent genomes, the full profile still reached genus 8/11, species 3/11 versus 1/11 and 0/11 parametric-only, so alignment-free, homology and CRISPR fields carried host-discriminative signal without nucleotide-level matches. Restricting to the ten cleanly post-cutoff genomes preserved the contrast (full 7/10, 3/10 versus 2/10, 1/10). Four cases (⋆, Supplementary Table 27) illustrate the range: *Streptococcus* phage P16 was correctly called under the full profile from concordant CRISPR (234 spacer matches, 100% identity) and RBP evidence pointing to *S. thermophilus*, but misassigned to an unrelated Gram-positive genus parametric-only; *Escherichia* phage Henuyfy22N was correctly identified as *E. coli* from RBP homology alone, without a labelled BLASTN neighbour or CRISPR record; the novel jumbo *Streptomyces* phage DeluluLabubu was misassigned under every evidence condition, as RBP homology (pointing to *Clostridium*/*Bacillus*) conflicted with 25-mer sharing (pointing exclusively to *Lactobacillus*) and the model followed the 25-mer signal; and *Klebsiella* phage Skif1059 was misassigned under every evidence condition, as tail-fibre homology to Enterobacteriaceae receptors was outweighed by spurious alignment-free similarity to unrelated Firmicutes, including a wrong Gram-status call, identifying it as a systematic hard case. *Vibrio* phage Va260-JW1 (not starred) was non-monotonic—species-correct only with RBP withheld (norbp), genus-correct but species-incorrect under both full and parametric-only—illustrating that not every case follows a clean evidence-dependence ordering at this small sample size. These results show that, for this backbone, evidence-supported correctness and evidence-conditioned failure both persist on prospective genomes, extending the dependence on named retrievable evidence documented on RefSeq-634.

#### Supplementary Note S13 | External wet-lab-supported validation and structured-evidence perturbation on 207 phage–host pairs

*Cohort.* We assembled 207 phage–host associations with experimental host support from six studies published in 2025–2026, in two collection rounds (136 and 71 genomes) that screened 23 candidate publications in total. Ground-truth hosts were taken verbatim from the source publications (positive-labelled species per accession) and were not re-derived computationally. Every genome was resubmitted through the unchanged production submission pipeline, with no cache reuse, so retrieval, profile construction and inference all ran under one frozen snapshot against approximately 494 registered candidate host species. All 207 submissions parsed successfully (0 abstentions, 0 parse failures). The six contributing studies, with the number of genomes each supplied, were: a temperate human-gut prophage isolation study (134; `10.1038/s41586-025-09614-7`); the KlebPhaCol *Klebsiella* resource (52; `10.1093/nar/gkaf1122`); a broad-host-range *Vibrio* phage study (18; `10.1093/ismejo/wraf063`); a hospital-specific *Enterobacter cloacae* cocktail design (1; `10.1038/s41564-025-02130-4`); a *Staphylococcus* curved-contractile-tail phage report (1; `10.1128/mbio.03829-25`); and a deep-sea prophage-encoded chitinase study (1; `10.1093/ismejo/wraf004`). Three studies therefore supplied 204 of the 207 pairs, so per-host accuracies below are dominated by the host panels of those cohorts rather than by an independently sampled cross-section of phage–host biology.

*Reference corpora (frozen snapshot).* All reference resources were frozen for the duration of the evaluation and are recorded verbatim in the run configuration. The candidate catalogue presented to the model comprised 494 host species, taken from a registry of 571 host summaries at freeze time. Whole-genome nucleotide neighbours were retrieved from a host-labelled phage-genome corpus of 1,940 genomes, extended to 6,425 genomes on the resubmission path; self-matches were excluded at query time by skipping the query accession rather than by physically rebuilding the database. Receptor-binding-protein homology was assessed by DIAMOND against 19,081 structurally annotated proteins drawn from 8,525 phage accessions and spanning 239 host species. Alignment-free comparison used exact 25-mer sharing against a pool of 723 host genomes (703 original plus 20 backfilled when the genome-archival defect described in the run log was repaired). CRISPR spacers were taken both from a precompiled public spacer database and from spacers called with PILER-CR directly on the registered host genomes. Because the receptor-binding-protein library covers 239 host species against a 494-species candidate catalogue, fewer than half of the candidates can receive a same-species hit in that channel even in principle; the coverage of the remaining channels is reported per query in Supplementary Table 31c.

*Overall accuracy.* Species-level top-1/3/5/10/30 accuracies were 52.2%, 68.6%, 82.1%, 91.8% and 97.6%; genus-level values were 90.8%, 96.1%, 97.6%, 98.1% and 100.0% (Supplementary Table 31). The 38.6-percentage-point gap between genus-level and species-level top-1 accuracy locates most residual error within closely related species of a correctly identified genus, matching the within-genus error structure documented on RefSeq-634 in the main text.

*Per-host-species breakdown.* Accuracy varied widely by confirmed host (Supplementary Table 31). *Klebsiella pneumoniae* ($n = 52$) reached 90.4% species-level top-1, whereas *Bacteroides caccae* ($n = 33$) reached 27.3% with 100.0% genus-level top-1 and *Bacteroides thetaiotaomicron* ($n = 8$) 0.0% with 100.0% genus-level top-1, so the largest species-level deficits occurred in host groups whose confirmed species was consistently out-ranked by a congener. *Evidence availability.* Of the 207 queries, 138 (66.7%) carried a CRISPR record and 153 (73.9%) had a labelled BLASTN neighbour matching the confirmed host; 164 profiles carried a STRONG BLASTN host signal. Among confirmed-host BLASTN hits the median identity was 96.8% and the median query coverage 31%.

*Field-removal perturbation.* Each of the 207 profiles was re-scored under seven conditions that modify only the targeted evidence block, leaving every other byte, the model, the prompt and the candidate catalogue unchanged (1,449 inference calls in total; Supplementary Table 32). Every condition is scored over the complete set of 207 queries, with a query counted as incorrect whenever the confirmed host is not returned at rank 1, so the conditions share one denominator and are directly comparable to the full-profile accuracy. Ordering the conditions by species-level top-1 accuracy gives: full profile 52.2%, −25-mer 54.1%, −RBP 46.4%, BLASTN host label masked 43.0%, −CRISPR 38.6%, −BLASTN 34.8%, no structured evidence 18.8%. Relative to the full profile, the losses were −17.4 pp (BLASTN), −13.5 pp (CRISPR) and −5.8 pp (RBP), whereas removing the 25-mer block *increased* accuracy by +1.9 pp. Removing all specialised evidence cost −33.3 pp, so roughly two-thirds of the species-level accuracy was contributed by retrieved evidence rather than by unaided model priors. Masking only the BLASTN host labels, while retaining the quantitative identity and coverage values, cost −9.2 pp, i.e. 53% of the loss incurred by deleting the entire block; both the host identity and the alignment statistics therefore carried signal.

*Directed label counterfactual.* For every query with a single dominant host-labelled evidence record, the host label of that record was replaced by a same-genus, same-family or strongest-competing decoy while all numeric and structural fields were held byte-identical. Across 185 evaluable queries the top-1 prediction switched to the introduced decoy in 0 cases (0%) and the median change in true-host rank was 0. Retrieved host labels therefore informed but did not dictate the output, consistent with the partial loss observed under label masking.

*Limitation.* The 207-pair cohort carries only positive host labels; the 59 nominated pairs available for one-to-one scoring were all positives, and only 47 species-level negative calls could be derived from strain-level host-range tables. Discrimination metrics requiring negatives (ROC-AUC, MCC) are therefore not reported for this cohort.
